# Structure-function coupling in the human brainstem

**DOI:** 10.64898/2026.04.05.716603

**Authors:** Asa Farahani, Subhranil Koley, Justine Y. Hansen, Firdaus Fabrice Hannanu, María Guadalupe García-Gomar, Kavita Singh, Simone Cauzzo, Filip Milisav, Zhen-Qi Liu, Vincent Bazinet, Marta Bianciardi, Bratislav Misic

**Author notes:** These authors contributed equally to this work (Asa Farahani, Subhranil Koley, Justine Y. Hansen). These authors contributed equally to this work (Marta Bianciardi, Bratislav Misic).

## Abstract

The nuclei of the brainstem modulate neuronal activity throughout the central nervous system. Yet connectome reconstructions typically do not include the brainstem because it is notoriously difficult to image. As a result, the influence of the brainstem on structure-function coupling in the brain is unknown. Here we use high-resolution 7 Tesla magnetic resonance imaging (MRI) to reconstruct structural and functional brain connectomes encompassing cortex and 58 brainstem nuclei spanning the midbrain, pons, and medulla. We identify structural connectional profiles of individual brainstem nuclei to the cortex and find that they align with a spectrum of functions, spanning sensory and motor processing to higher-order cognition. Structural and functional connectivity in brainstem-augmented connectomes are positively correlated, and pairs of regions with direct anatomical projections display greater functional connectivity than pairs without. Structure-function coupling is heterogeneous across brainstem nuclei, with strong coupling in both modulatory and relay nuclei. Collectively, this work presents an initial step toward understanding how the brainstem shapes structure-function relationships in the brain.

## INTRODUCTION

Anatomical projections between neuronal populations promote synchrony and the emergence of patterned neural activity. This structure-function relationship can be measured *in vivo* using techniques such as diffusion-weighted magnetic resonance imaging (MRI) and functional MRI. Early studies emphasized global relationships between structural and functional connectivity^1–7^, while more recent studies have found that structure-function coupling is also spatially heterogeneous^8–12^. Despite improvements in acquisition, preprocessing, and sample size, most extant studies focus on cortical structure-function coupling, excluding the contributions of the brainstem. Exclusion of the brainstem is partly attributable to technical difficulties associated with imaging deep brain structures. These difficulties include the presence of physiological noise and motion artifacts arising from large vasculature and cerebrospinal fluid flow surrounding the deep structures, susceptibility-related signal dropout, and challenges in parcellating small brainstem nuclei in low-resolution MR images^13–22^. Nevertheless, brainstem nuclei modulate the relationship between cortical structure and function via direct and indirect neuronal projections and diffuse neurotransmitter release^23–29^. Hence, including the brainstem and its connections is necessary to capture a complete picture of brain-wide structure-function coupling^10^.

Recent technical and analytical advances make it increasingly possible to image brainstem nuclei *in vivo* and include them in connectome reconstructions. These include ultra-high-field MRI^30–32^, advanced image acquisition techniques^31,33,34^ , and brainstem-specific pre-processing pipelines designed to eliminate physiological noise^35–37^. Concurrent development of *in vivo* atlases of brainstem nuclei in stereotactic space provides an anatomical reference for brainstem imaging^38–44^. Together, these advances have enabled a shift from reconstructing specific brainstem fiber bundles with limited and predefined brainstem seeds and cortical targets^45,46^ and studying pathways involved in specific circuitry (e.g., pain circuitry^47^), toward reconstructing large-scale networks of structural and functional connectivity that span multiple brainstem nuclei and encompass nearly the entire cortex and subcortex^48,49^.

Here we use high-resolution diffusion-weighted and resting-state functional MRI acquired at 7 Tesla in 19 healthy adults to reconstruct large-scale structural and functional networks spanning 58 brainstem nuclei and the cerebral cortex. We characterize the organization of structural connectivity between brainstem and cortex, examine nucleus-specific cortical connectivity profiles, and assess the correspondence between structural and functional connectivity within and between the brain-stem and cortex.

## RESULTS

Structural connectivity of the cortex and the brainstem was estimated using diffusion-weighted MR images acquired on a 7 Tesla scanner from 19 unrelated healthy participants (mean ± SE age: 29.74 ± 1.21 years; 10 males, 9 females). Structural connections were defined based on specified cortical and brainstem regions (see *Methods* for details on data acquisition, data preprocessing, and structural network reconstruction). Cortical regions were defined according to the 400 regions in the Schaefer parcellation^53^, and brainstem nuclei were defined according to the 58 nuclei in the Brainstem Navigator atlas (Table. S1, 50 bilateral and eight midline nuclei; Figs. 1a,b and S1; atlas available at https://www.nitrc.org/projects/brainstemnavig)^38–44^.

**Figure 1.**
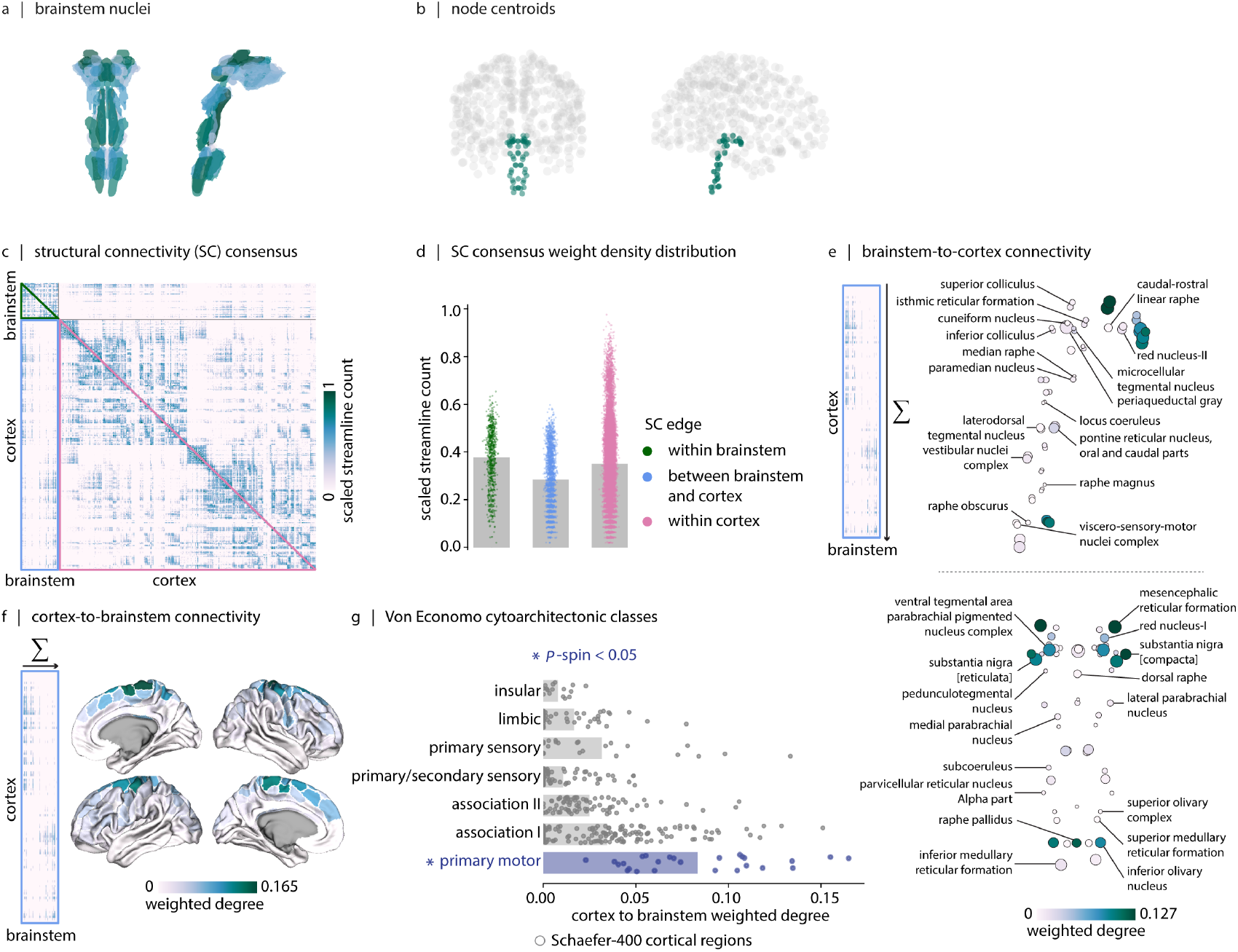
Structural connectivity of brainstem and cortex. (a) Sagittal and coronal views of the thresholded (35%) probabilistic template for all 58 brainstem nuclei in the Brainstem Navigator atlas (https://www.nitrc.org/projects/brainstemnavig). These nuclei are further displayed on the MNI template in Fig. S1. (b) Sagittal and coronal views show parcel coordinate centroids for cortical regions (gray points, 400 cortical regions) and brainstem nuclei (green points, 58 brainstem nuclei). (c) Group-consensus weighted structural connectivity (SC) matrix (58 brainstem nuclei and 400 cortical regions, 458 regions × 458 regions). The green triangle highlights the brainstem to brainstem connections, the blue rectangle highlights the connections between cortex and brainstem, and the pink triangle highlights the cortex to cortex connections. (d) Density distributions of non-zero structural connectivity edges within brainstem (green dots), between brainstem and cortex (blue dots), and within cortex (pink dots). Structural connections within brainstem are stronger than structural connections between brainstem and cortex (two-tailed, *p*_*perm*_ = 9.99 × 10^−4^, *N*_*perm*_ = 1 000). Structural connections within cortex are also stronger than structural connections between brainstem and cortex (two-tailed, *p*_*perm*_ = 9.99 × 10^−4^, *N*_*perm*_ = 1 000). (e) Brainstem-to-cortex weighted degree is calculated by summing the structural connectivity of each brainstem nucleus across all cortical regions. Sagittal and coronal views of brainstem nuclei are shown. Node sizes are proportional to the number of voxels per brainstem nucleus, and node colors reflect weighted degree. To test whether brainstem nuclei hubness rankings depend on differences in volume, we regressed out the voxel-count effect from the weighted degree—the ranking of nuclei is largely preserved, suggesting that hubness is not driven by volume (Fig. S3). (f) Cortex-to-brainstem weighted degree is calculated by summing the structural connectivity of each cortical region across all brainstem nuclei. Brain cortical maps are displayed on fs-LR midthickness surfaces. (g) Cortex-to-brainstem weighted degree binned according to the seven cytoarchitectonic classes defined by von Economo (Fig. S4)^50–52^. Each dot represents a cortical region defined according to the Schaefer-400 parcellation^53^. The brainstem shows strong structural connectivity with the primary motor cortex (two-tailed, *p*_*spin*_ = 1.998 × 10^−3^, *N*_spin_ = 1 000). In Fig. S5, we also bin the cortex-to-brainstem weighted degree according to the Yeo 7-networks^54^. For completeness, we also show the cortex-to-cortex weighted degree in Fig. S6. Mean weighted degree patterns for both structural and functional connectivity across all different network compartments are shown in Fig. S8.

### Structural connectivity of brainstem and cortex

Fig. 1c shows the group-consensus structural connectivity (SC) matrix derived from diffusion-weighted imaging. The consensus structural connectivity matrix is constructed using an edge consistency- and length-based thresholding approach (see *Methods*)^57^. Structural connections within brainstem and within cortex are stronger than structural connections between brainstem and cortex (two-tailed, *p*_*perm*_ = 9.99 × 10^−4^, *N*_*perm*_ = 1 000, Fig. 1d). Fig. S2a shows the inter-subject variability of the structural connectome. Overall, structural connectivity patterns between brainstem and cortex exhibit greater inter-subject variability compared to within-brainstem and within-cortex structural connectivity patterns.

With the consensus structural connectivity in place, the first question we ask is which brainstem nuclei are most strongly connected to cortex, and which cortical regions are most strongly connected to brainstem? Throughout the manuscript, we refer to the connectivity profile of individual brainstem nuclei with cortex as “brainstem-to-cortex” connectivity and vice versa as “cortex-to-brainstem” connectivity. Importantly, the brainstem-augmented structural connectome is symmetric, and the terms “brainstem-to-cortex” and “cortex-to-brainstem” indicate only the axis along which connectivity values are summarized, and not anterograde or retrograde anatomical directionality. The overall structural connectivity of each brainstem nucleus to cortex is quantified as the sum of SC values across all cortical regions (i.e., “weighted degree”) (Fig. 1e). When brainstem nuclei are ranked by weighted degree, the substantia nigra, mesencephalic reticular formation, raphe pallidus, ventral tegmental area, and inferior olivary nucleus are among the nuclei with the highest connectivity to cortex (brainstem-to-cortex hubs). Given that brainstem nuclei vary in regional volume, we further remove the voxel-count effect from the weighted degree and find that the ranking of nuclei is largely preserved (Fig. S3).

The overall structural connectivity of each cortical region to brainstem is quantified as the sum of SC values across all brainstem nuclei (i.e., “weighted degree”) (Fig. 1f). Cortical connectivity to brainstem is concentrated in primary motor cortex (cortex-to-brainstem hubs). To provide an anatomical description of this pattern, we compute mean cortex-to-brainstem weighted degree within each cortical cytoarchitectural class defined by the histological von Economo atlas (Fig. S4)^50–52^. This analysis shows a significant enrichment of cortex-to-brainstem structural connectivity within the primary motor cytoarchitectonic class (two-tailed, *p*_*spin*_ = 1.998 ×10^−3^, *N*_spin_ = 1 000; Fig. 1g). Notably, the motor cortex contains regions of the somato-cognitive action network (SCAN) interleaved with effector-specific motor regions^58^. Whether the observed enrichment reflects stronger connectivity with SCAN regions, effector-specific motor regions, or both, is an important follow-up question. In Fig. S7, we also show the subcortical regions to brainstem nuclei structural connectivity. The brainstem is strongly structurally connected to the thalamus proper and hypothalamus. The accumbens area is the region with the weakest structural connectivity to brainstem.

### Connection profiles of brainstem nuclei and cortical functional specialization

In the previous section we considered global connectivity profiles, aggregated over all cortical or brainstem nodes. Here we also examine the connectivity profiles of individual brainstem nuclei with cortex. To account for both direct monosynaptic and indirect polysynaptic influences, we estimate the weighted communicability of cortex to each brainstem nucleus. Communicability between two nodes is defined as the weighted sum of all paths and walks between those nodes^59–61^. Communicability does not exclusively represent direct tractography-derived connections, instead, it provides a measure of communication potential by incorporating both direct and indirect paths through the broader network architecture. This is particularly relevant when relating structural connectivity to functional activation patterns, since functional interactions are further mediated by indirect anatomical pathways rather than by direct or shortest-path connections only. Consistently, prior work has shown that diffusion models of polysynaptic connectivity, particularly communicability measures improve the correspondence between structural and functional connectivity and can better predict behavior and cognition than direct structural connectivity alone^11,61–64^.

Cortical connectivity patterns vary across brainstem nuclei, not only in overall connection strength but also in the spatial distribution of dominant cortical targets. Fig. 2a shows cortical structural communicability patterns for three example nuclei including the superior colliculus, raphe pallidus, and ventral tegmental area (averaged pattern across left and right nuclei for superior colliculus and ventral tegmental area). We specifically focus on these three examples because prior animal work consistently demonstrates their distinct anatomical connectivity and functional specialization. The superior colliculus is connected to visual areas and has a role in visually guided behavior^65–68^. The raphe pallidus has diffuse connections to motor and sensory regions and has a role in motor and sensory regulation^69^. The ventral tegmental area is connected to frontal cortex and has roles in positive and negative reinforcement, decision making, and working memory^70–73^. Indeed, superior colliculus exhibits structural communicability with visual cortical regions, whereas raphe pallidus exhibits structural communicability with the motor and the frontal cortical regions and ventral tegmental area exhibits structural communicability with the frontal cortical regions. Together, these findings show heterogeneous, nucleus-specific cortical targeting patterns across the brainstem. For further details on the correspondence between the reconstructed connectivity profiles and current anatomical knowledge for these selected nuclei see *Supplementary information* (also see Fig. S10).

**Figure 2.**
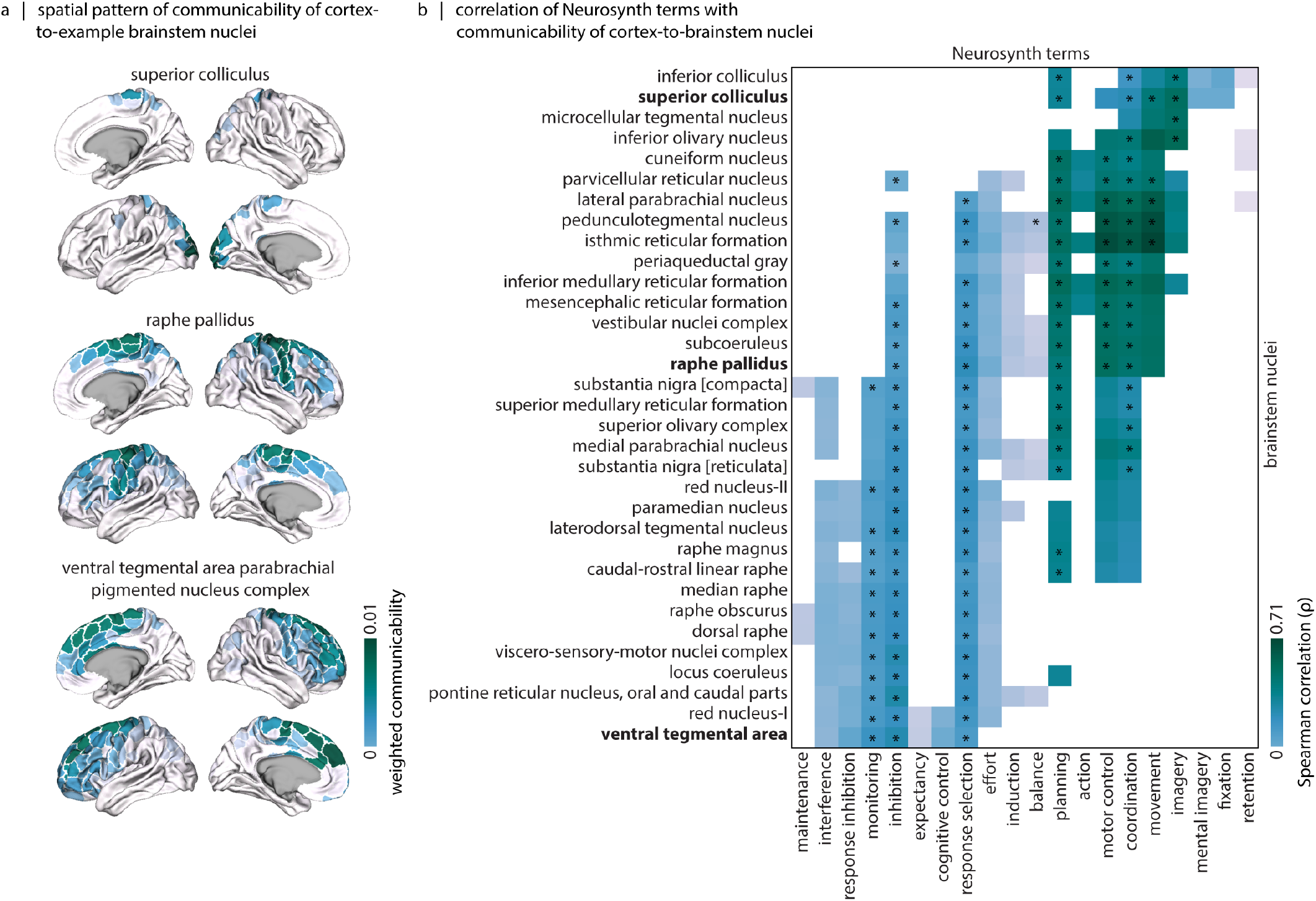
Connection profiles of brainstem nuclei. (a) Structural communicability between selected brainstem nuclei and cortex. Cortical maps are displayed on fs-LR midthickness surfaces. Connection profiles of all other brainstem nuclei are shown in Fig. S9. (b) The heatmap visualizes Spearman correlation values (*ρ*) between structural communicability pattern of each nucleus and Neurosynth meta-analytic cortical maps. Neurosynth functional/cognitive terms for which at least one nucleus exhibited a positive correlation exceeding thresholds derived from spatial autocorrelation–preserving permutation nulls are shown on the *x*-axis (*p*_spin_ *<* 0.05, *N*_spin_ = 1 000). We further apply False Discovery Rate (FDR) correction to the associations calculated per functional term; the associations that survive correction are indicated with an asterisk (*). Bold nucleus names denote nuclei whose cortical communicability patterns are shown in panel (a). Communicability patterns are averaged to obtain a single map for each pair of bilateral nuclei. A list of the 123 Neurosynth terms from the Cognitive Atlas^55,56^ is provided in Table. S2. We further examine the correspondence between structural communicability pattern of each brainstem nucleus and a series of task-fMRI contrast maps from the Human Connectome Project (HCP); these results are shown in Fig. S11.

We pursue this idea further and ask whether connections between brainstem nuclei and cortex differentiate classes of sensory, motor and cognitive functions. We cross-correlate the cortex-to-brainstem structural communicability pattern of each brainstem nucleus with the cortical maps from the Neurosynth meta-analytic atlas^55,74^ (Fig. 2b). This analysis links the heterogeneity in brainstem nuclei structural connectivity to functional specialization of the nuclei. Structural communicability patterns for nuclei including superior colliculus and inferior colliculus show spatial correspondence with Neurosynth maps associated with visual and oculomotor functions, including “imagery” and “fixation”. Structural communicability patterns for nuclei including cuneiform nucleus, isthmic reticular formation, raphe pallidus, and pedunculotegmental nucleus show strong spatial correspondence with Neurosynth maps associated with motor functions, including “movement”, “motor control”, and “coordination”. Structural communicability patterns for nuclei including ventral tegmental area and red nucleus (subregion 1) show spatial correspondence with Neurosynth maps associated with higher-level cognition, including “inhibition”, “monitoring”, and “interference”. Altogether, we find that the projection patterns of brain-stem nuclei cover a wide spectrum of cognitive functions, motivating more detailed exploration of structure-function correspondences of these nuclei.

### Structure-function coupling in brainstem-augmented connectomes

In the following sections we assess structure-function coupling in connectomes that include the brainstem. In addition to the diffusion–weighted MRI data described above, we analyze resting-state functional MRI (fMRI) acquired on the same 7 Tesla MR scanner (see *Methods* for details on fMRI data acquisition, data preprocessing, and functional network reconstruction). A detailed characterization of brainstem functional connectivity is discussed in a previous report from our group^49^. Here we aim to characterize the correspondence between structural and functional connectivity between brainstem and cortex, and within the brainstem.

Fig. 3a shows the group-consensus weighted SC and group-averaged functional connectivity (FC) matrices. Structure-function correspondence is quantified separately within three anatomically defined network compartments: within brainstem, between brainstem and cortex, and within cortex. For each compartment, structure-function coupling is defined as the Spearman correlation between corresponding structural and functional edge weights. Structure-function coupling is weaker for connections between brainstem and cortex (*ρ* = 0.12, *p*_*rewired*_ = 9.99 × 10^−4^, *N*_*rewired*_ = 1 000) compared to connections within brainstem (*ρ* = 0.42, *p*_*rewired*_ = 9.99 × 10^−4^, *N*_*rewired*_ = 1 000) and connections within cortex (*ρ* = 0.32, *p*_*rewired*_ = 9.99 × 10^−4^, *N*_*rewired*_ = 1 000) (Fig. S12c). At the individual level, inter-subject structure-function coupling shows a similar pattern, with weaker coupling for brainstem-cortex connections and stronger coupling for connections within the brainstem and within the cortex (Fig. S12d).

**Figure 3.**
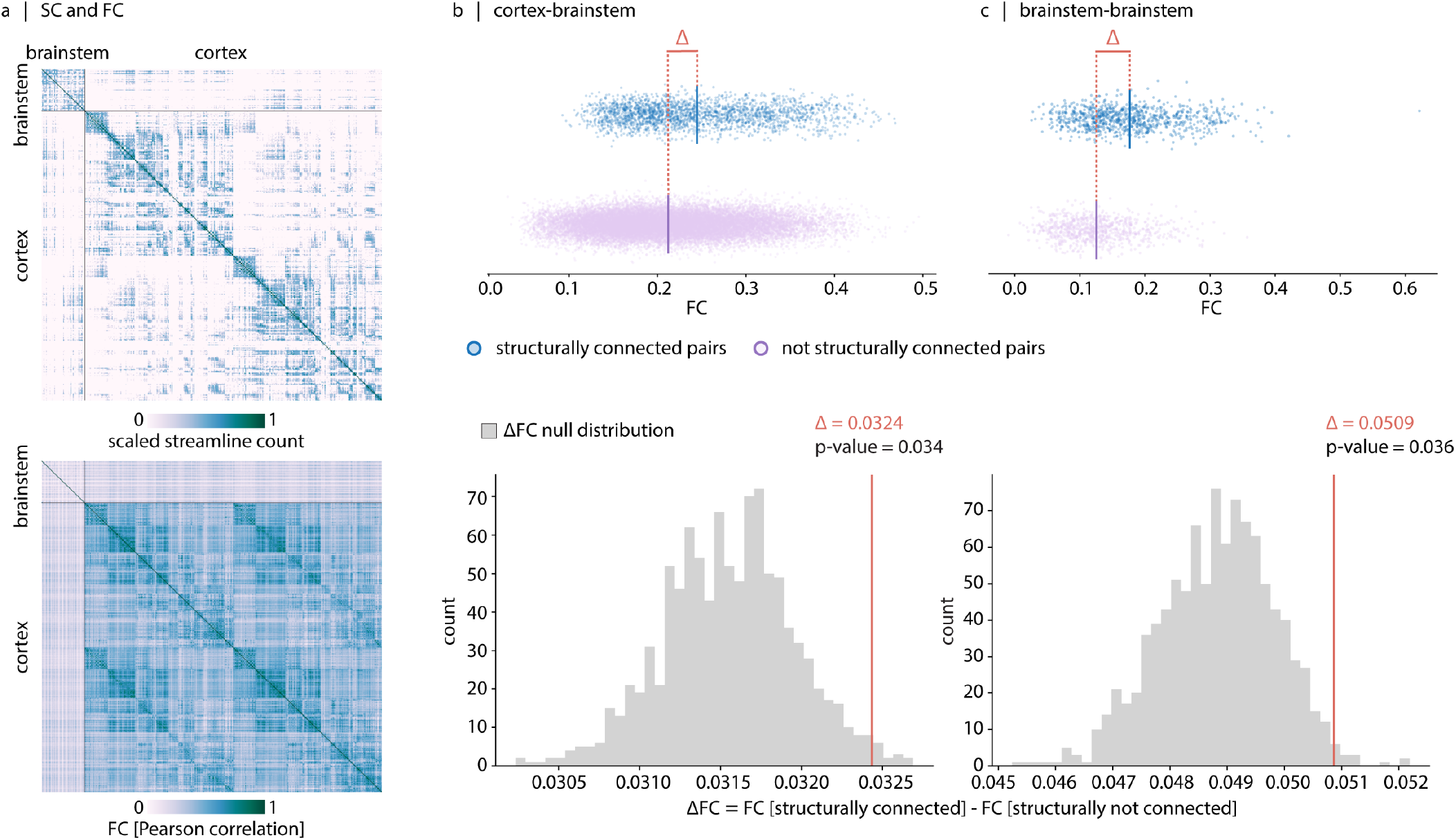
Brainstem structural connectivity supports its functional connectivity. (a) Group-consensus weighted structural connectivity matrix (top) and group-averaged functional connectivity matrix (bottom) spanning 58 brainstem nuclei and 400 cortical regions (458 regions × 458 regions). (b) Brainstem-cortex functional connectivity is stronger for structurally connected region pairs than for not-connected pairs. (c) Brainstem-brainstem functional connectivity is also stronger for structurally connected nuclei than for anatomically not-connected nuclei. For both (b) and (c), statistical significance of the difference in mean functional connectivity between connected and not-connected region pairs is assessed against a null distribution of differences (ΔFC) generated when degree- and edge length-preserving rewired structural networks (*N*_*rewired*_ = 1 000) are used to define anatomical connectivity^75^ (see Fig. S13). The FC difference (ΔFC) corresponded to Cohen’s *d* of 0.45 for the cortex-brainstem compartment and 0.79 for the brainstem-brainstem compartment. For completeness, in Fig. S14 we show that cortex-cortex functional connectivity is stronger for structurally connected region pairs than for not-connected pairs, consistent with previous reports^3,76^. For individual-level analyses, see Fig. S15.

We next examine whether the existence of direct anatomical projections is associated with stronger functional connectivity between brainstem and cortex, and within brainstem. We estimate the difference in functional connectivity between region pairs that are anatomically connected (SC ≠ 0) and those that are not (SC = 0). To assess the statistical significance of these differences, we generate 1 000 degree- and edge-length–preserving rewired structural networks and use them to construct null distributions for the differences in mean functional connectivity between anatomically connected and non-connected region pairs. In both the brainstem-cortex and brainstem–brainstem compartments, the empirical differences in mean functional connectivity exceeded those expected under the null hypothesis (two-sided, ΔFC = 0.0324, *p*_*rewired*_ = 0.034; ΔFC = 0.0509, *p*_*rewired*_ = 0.036, respectively). These results indicate that regions linked by direct anatomical projections exhibit significantly stronger functional coupling than regions lacking such structural connections (Fig. 3b,c).

The difference in functional connectivity between structurally connected and unconnected brainstem-anchored pairs is modest in magnitude. However, this effect is comparable to that observed for cortex-cortex pairs, where SC-FC correspondence is further characterized^10^ (see Fig. S14; ΔFC = 0.068, *p*_*rewired*_ = 9.99 × 10^−4^). Across all three network compartments (brainstem-cortex, cortex-cortex and brainstem-brainstem), the empirical differences exceed those expected from degree- and edge-length–preserving structural null models and are associated with effect sizes (Cohen’s *d*) greater than 0.45. Moreover, stronger functional association of structurally connected pairs is also consistently observed at the individual-subject level (Fig. S15). This indicates that the association between anatomical projections and stronger functional coupling is not driven solely by structural network topology or by group averaging. Nevertheless, structure-function correspondence remains imperfect, and the ΔFC values remain moderate, consistent with the idea that functional interactions can arise via indirect structural connections that produce coherent time courses among regions separated by two or more synapses^3,10,78,79^.

Before examining the correspondence between structural and functional connectivity of brainstem in detail, we assess how structural and functional connectivity strength is linked to the Euclidean distance between seed and target regions and to the volume of seed and target regions. Increased inter-regional distance between seed and target regions is accompanied by weaker structural connections (within cortex: *ρ* = −0.55, *p* ≈ 0; within brainstem: *ρ* = −0.16, *p* = 1.94 × 10^−6^; between brainstem and cortex: *ρ* = −0.05, *p* = 0.01). Larger regional volume of seed and target (target number of voxels × seed number of voxels) is accompanied by stronger structural connectivity, particularly within brainstem (*ρ* = 0.37, *p* = 9.89 *×* 10^−29^) and between brainstem and cortex (*ρ* = 0.25, *p* = 2.32*×*10^−36^) (within cortex: *ρ* = 0.03, *p* = 5.53 *×* 10^−5^) (Fig. S16). Increased inter-regional distance between seed and target regions is also accompanied by weaker functional connections particularly within the cortex (*ρ* = − 0.30, *p ≈* 0) and within the brainstem (*ρ* = − 0.11, *p* = 1.56 × 10^−5^). Larger regional volume of seed and target (target number of voxels × seed number of voxels) is accompanied by stronger functional connectivity, particularly within the brainstem (*ρ* = 0.65, *p ≈* 0) and between the brain-stem and cortex (*ρ* = 0.60, *p ≈* 0) (within the cortex: *ρ* = − 0.01, *p* = 2.99 × 10^−4^) (Fig. S17). Given the dependence of structural and functional connectivity on inter-regional distance and regional volume, we incorporate these geometrical features when evaluating structure–function correspondence of brainstem nuclei in the next section.

### Structure-function coupling in brainstem nuclei

Here we ask how structure-function coupling differs for individual brainstem nuclei. Given that both inter-regional distance between seed and target and their regional volume affect structural and functional connectivity between seed and target regions, correspondence between SC and FC of two regions may arise from the regional geometric features. Therefore, for each brain-stem nucleus, to assess whether SC patterns explain any variance in FC patterns beyond geometric features, we compare two predictive models of FC: a baseline regression model including only inter-regional distance and regional volume (defined as the product of seed and target voxel counts), and an extended regression model that further incorporated SC communicability weights (Fig. 4a, left). To evaluate the specificity of SC contributions, we construct 1 000 null SC networks with preserved degree- and edge-length distributions, and refit the extended model using each null network. We compare improvement in model fit using the empirical SC, relative to improvement in model fit when null networks are used.

**Figure 4.**
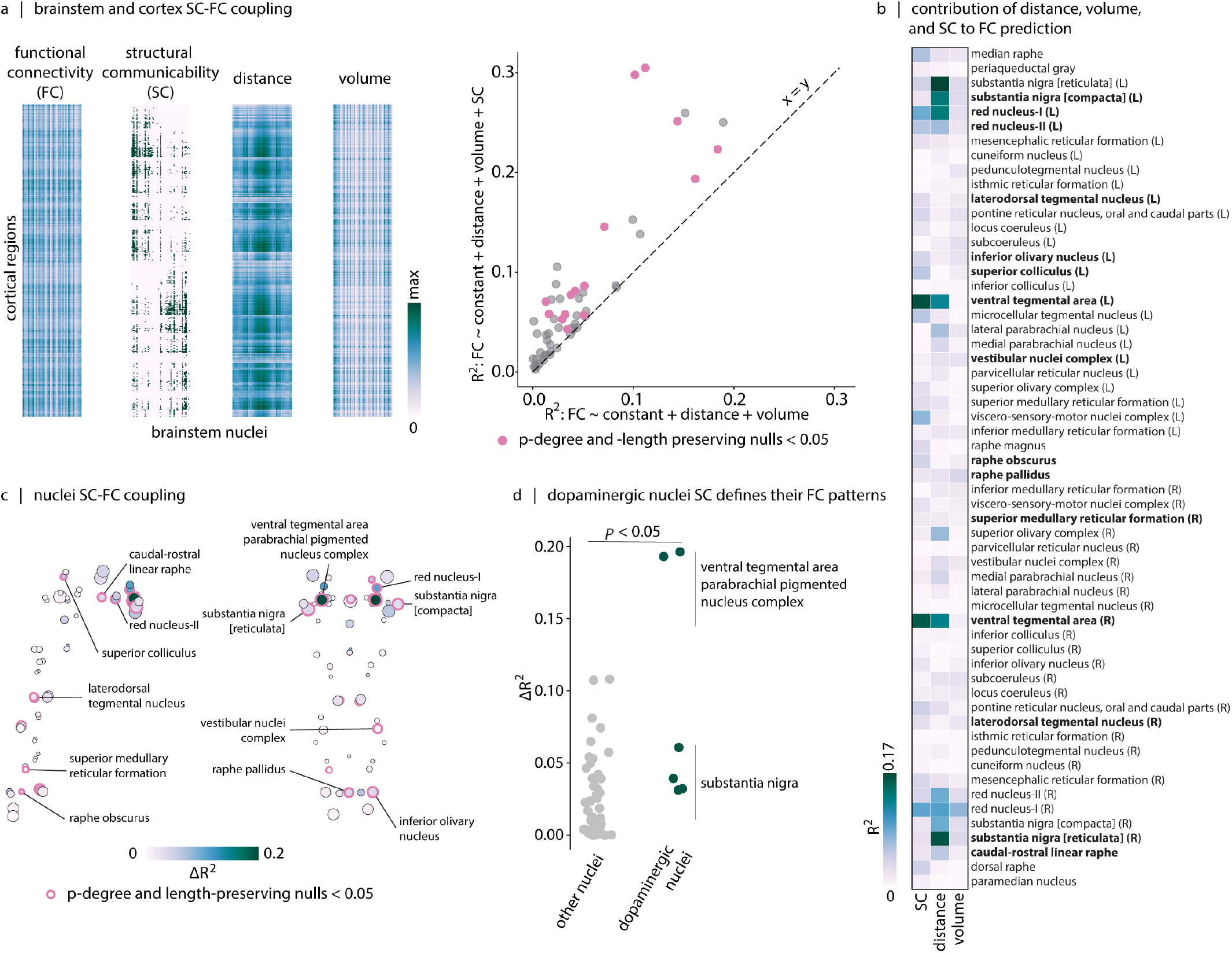
Structure-function coupling in brainstem nuclei. (a) The two leftmost heatmaps show resting-state functional connectivity (FC) and structural communicability between brainstem nuclei and cortical regions. The third heatmap shows the Euclidean distance between each brainstem nucleus and each cortical region. The fourth heatmap shows the product of the number of voxels in each brainstem nucleus and each cortical region, providing an estimate of their combined volumetric size. We construct two regression models to predict the FC profile of each brainstem nucleus with cortical regions. The baseline model includes distance and volume regressors and the extended model includes distance, volume and structural communicability profile of the nucleus as regressors. In the scatter plot, baseline model *R*^2^ values are shown on the *x* axis and extended model *R*^2^ values are shown on the *y* axis. Each dot in the scatter plot corresponds to a brainstem nucleus. We assess the significance of improvement in model performance (Δ*R*^2^) by comparing the empirical Δ*R*^2^ value to a null distribution of Δ*R*^2^ values obtained from models in which structural communicability is replaced by a communicability matrix that is obtained from degree- and length-preserving null structural networks. Brainstem nuclei showing a significant increase in *R*^2^ when empirical structural communicability is included in the model are shown in pink. (b) For each brainstem nucleus, the contributions of distance, volume and structural communicability to FC prediction (*R*^2^) are shown (quantified using the dominance analysis^77^). The nuclei written in bold are the ones that show a significant increase in *R*^2^ when empirical structural communicability is included. (c) The Δ*R*^2^ values for brainstem nuclei are mapped onto the sagittal and coronal views of the brainstem. The ventral tegmental area shows a marked improvement in FC prediction following the inclusion of structural communicability in the model (see Fig. S18). (d) Dopaminergic nuclei exhibit larger Δ*R*^2^ values, indicating stronger structure–function coupling beyond geometric constraints (inter-regional distance and volume of target and seed regions) (*p*_*perm*_ = 1.99 *×* 10^−3^, *N*_*perm*_ = 1 000).

Incorporating structural connectivity between cortex and brainstem nuclei improves prediction of functional connectivity above what can be solely explained by geometric features in 15 out of 58 brainstem nuclei (Fig. 4a, right). These nuclei include both slow-acting modulatory nuclei (such as dopaminergic ventral tegmental area and serotonergic raphe nuclei), as well as faster-acting relay nuclei (inferior olive, superior colliculus, red nucleus) (Fig. 4b,c), consistent with the notion that cortical functional connectivity is shaped by both fast sensory-motor signaling and neuromodulation^80^. The contribution of dopaminergic nuclei is particularly noteworthy (ventral tegmental area and substantia nigra; Fig. 4d), consistent with the hypothesized contribution of dopaminergic signaling to a wide spectrum of cognitive functions^81^.

For completeness, we also investigate the correspondence between structural and functional connectivity within individual brainstem nuclei. To assess whether brainstem SC explains any variance in brainstem FC of specific nuclei beyond geometric features, we apply the same modeling framework described above and compare two predictive models of FC: a baseline model including only inter-regional distance and regional volume, and an extended model that additionally incorporates SC communicability weights (Fig. S19a). Relative to degree- and length-preserving null models, empirical SC significantly improves FC prediction in 19 of 58 brainstem nuclei (Fig. S19b). The greatest correspondence is observed in a similar set of nuclei reported above (that made the greatest contribution to global structure-function coupling; Fig. S20), including ventral tegmental area, substantia nigra, several midline raphe nuclei, as well as important midbrain hubs such as the periaqueductal gray and mesencephalic reticular formation, involved in integration and arousal, respectively^82^.

### Replication analyses

Finally, we assess whether the principal structural organization and structure-function coupling patterns in the brainstem-augmented connectomes generalize beyond the 7T dataset used in this study. To this end, we perform a replication analysis using data from the Parkinson’s Progression Markers Initiative (PPMI)^83^. The PPMI dataset is acquired using 3T MRI, across multiple imaging sites, and in a participant sample with a different age range from the main cohort. We use 25 control participants from PPMI to reconstruct independent brainstem-augmented structural and functional connectomes, including the same 58 brainstem nuclei and 400 cortical parcels used in the main analysis.

First, we display the PPMI brainstem-augmented structural connectome in Fig. S21a. Consistent with the main analysis, structural connections within the brainstem and within the cortex are stronger than structural connections between brainstem and cortex (Fig. S21b). The structural organization of the brainstem-augmented connectome is preserved across datasets (Fig. S21c-f). The edge weights of the PPMI consensus structural connectome are strongly correlated with those of the main consensus structural connectome (*ρ* = 0.73; Fig. S21f). In addition, the weighted-degree profiles are highly similar across datasets for both brainstem-to-cortex connectivity (*ρ* = 0.91) and cortex-to-brainstem connectivity (*ρ* = 0.71). Again, cortex-to-brainstem weighted degree is enriched in primary motor and primary/secondary sensory cytoarchitectonic classes (Fig. S21e). These findings indicate that the main organizational features of the brainstem-augmented structural connectome are reproducible despite differences in scanner field strength, acquisition site, and participant characteristics.

We next look into the relationship between structural and functional connectivity in the PPMI dataset. Consistent with the main dataset, structural and functional connectivity are positively associated across the full brainstem-augmented connectome and within the three major network compartments (Fig. S22). Structurally connected region pairs again exhibit stronger functional connectivity than unconnected pairs in all three compartments. The observed differences in mean functional connectivity are ΔFC = 0.09 for cortex-cortex pairs (Fig. S23a), ΔFC = 0.04 for brainstem-cortex pairs (Fig. S23b), and ΔFC = 0.07 for brainstem-brainstem pairs (Fig. S23c). These effects correspond to Cohen’s *d* values of 0.86, 0.76, and 0.65, respectively. These differences exceed those expected under degree- and edge length-preserving rewired null models^75^ in all three compartments (*p*_*rewired*_ = 9.99 *×* 10^−4^, *N*_*rewired*_ = 1 000).

Last, we assess if including structural communicability improves prediction of the functional connectivity profiles beyond geometric constraints. As in the main analysis, we compare a baseline model including distance and regional volume with an extended model that additionally includes structural communicability. The spatial pattern of improvement in explained variance attributable to structural communicability is similar across datasets (*r* = 0.37; Fig. S23d,e). Consistent with the main dataset, dopaminergic nuclei show larger Δ*R*^2^ values than other nuclei (*p*_*perm*_ = 1.99 × 10^−3^ , *N*_*perm*_ = 1 000) (Fig. S23f). Together, these replication analyses show that both the structural organization of the brainstem-augmented connectome and its relationship with functional connectivity are not specific to the original 7T dataset, but are also detectable in an independent 3T multi-site cohort.

## DISCUSSION

In the present report, we use high-resolution diffusion-weighted and resting-state functional MRI data acquired using a 7 Tesla scanner together with a comprehensive brainstem atlas of 58 nuclei, to reconstruct a brainstem-augmented structural connectome and to examine the correspondence between structural and functional connectomes beyond the cerebral cortex. We identify connectional profiles of individual brainstem nuclei to the cortex and find that they align with a spectrum of functions, spanning sensory and motor processing to higher-order cognition. Finally, we characterize structure-function coupling in brainstem-augmented connectomes and study the contribution of the structural network embedding of individual nuclei to the emergence of inter-regional functional connectivity.

The study of human brainstem organization and its connections is historically based on *ex vivo* histological and fiber-dissection studies^84,85^, and is further guided by animal experiments^86–95^. These experiments have provided qualitative insights into brainstem anatomical connectivity, but the methodological heterogeneity of these studies makes cross-study comparison challenging and precludes their integration into a unified, quantitative structural connectivity matrix^96^. Recent advances in MR imaging have enabled *in vivo* structural imaging of the brainstem through the use of higher-strength scanners and sequences with contrasts that reveal borders of the small brainstem nuclei^31,38,97^. In a move toward quantitative, brainstem-augmented structural connectomes, earlier diffusion MRI studies have reconstructed major brainstem tracts^45^, or have visualized known pathways by defining specific brainstem nuclei and cortical seed regions. Here we build on these efforts by integrating brainstem nuclei into whole-brain structural connectomes using high-resolution diffusion MRI^48,98^.

We find that the brainstem connects to a wide variety of cortical regions and systems. The prominent anatomical connectivity between brainstem and motor cortex is expected from classical neuroanatomy, as the precentral gyrus gives rise to bilateral corticofugal pathways, including corticoreticular, corticorubral, and corticopontine tracts that terminate in brainstem nuclei, as well as the corticospinal tract that descends through the brainstem toward the spinal cord^87,99–101^. The connectivity of brainstem and motor cortex further reflects hardwired, evolutionary conserved motor pathways^102–104^. In addition, nuclei such as ventral tegmental area and substantia nigra are connected with frontal^89,105–107^ and somatomotor cortex^106,107^ and modulate their activity by dopaminergic signaling. These connections align with known mesocortical and nigrocortical dopaminergic pathways, possibly underlying processes such as inhibition control, performance monitoring and the integration of motor and cognitive information. Furthermore, connectivity with sensory and visual cortical areas is driven by nuclei such as superior colliculus that connect directly and indirectly with primary and extrastriate visual cortex (corticotectal tracts)^108,109^.

While the brainstem-augmented connectome introduced here recovers several structural connectivity patterns consistent with prior anatomical literature (see supplementary material, Fig. S10), we also observe some discrepancies relative to pathways reported in animal tracer and postmortem studies. For example, the ventral tegmental area is known to have direct connections to the hippocampus and the nucleus accumbens^110–114^, yet these connections do not appear as particularly strong edges in the present diffusion MRI-derived structural connectome. This discrepancy can be interpreted from two complementary perspectives. First, while tracer and postmortem studies establish the existence of specific anatomical projections, they do not always quantify projection strength relative to the full set of brainstem–cortical, brainstem–subcortical, and brainstem–brainstem pathways. Thus, a pathway may be well established anatomically while still appearing weak or inconsistently reconstructed in a diffusion MRI-derived connectome because of its relatively low macroscopic strength. Second, this discrepancy may reflect false negative connections, a known limitation of diffusion MRI tractography^115–117^. These limitations are especially relevant for sharply curving, or fanning pathways. Taken together, large-scale MRI-based approximation of brainstem connectivity should not be considered as a complete inventory of all microscopic projections. That said, diffusion MRI remains the only available *in vivo* method for mapping structural connectivity in the living human brain and is essential for comparing individuals, characterizing group differences, and constructing whole-brain models of neural communication and dynamics^115,118–120^.

White matter connections of the brainstem shape the emergence of functional connectivity. Structural and functional connectivity are positively correlated, and pairs of regions with direct anatomical projections display greater functional connectivity than pairs without. Structure-function coupling among brainstem nuclei is greater in magnitude than brainstem to cortex coupling, both at the group and individual participant level. This gradient suggests a hierarchical organization in which within brainstem circuits are strongly constrained by direct anatomical connections, while long-range brainstem-cortex projections operate in a diffuse modulatory and polysynaptic way with weaker SC–FC correspondence. Importantly, structure-function coupling is heterogeneous across brainstem nuclei. The greatest contributions are from a variety of modulatory and relay nuclei, associated with a variety of sensory-motor (e.g., superior colliculus, red nucleus, and inferior olive) and cognitive functions (e.g., ventral tegmental area, laterodorsal tegmental nucleus, and raphe nuclei), consistent with the idea that resting functional connectivity reflects the superposition of communication patterns originating from multiple circuits and systems^121,122^.

Although brainstem-cortex structure-function coupling was lower overall than within the brainstem, several nuclei, including the ventral tegmental area, raphe nuclei, laterodorsal tegmental nucleus, superior colliculus, inferior olive, and red nucleus, exhibited comparatively stronger coupling with cortical regions. These nuclei participate in organized ascending neuromodulatory and sensorimotor integration pathways, suggesting that SC–FC correspondence is higher when projections are functionally specific and embedded in structured loops. This indicates that coupling is not uniform but depends on the functional architecture of the underlying circuit. Collectively, these results contribute to an emerging literature on structure-function coupling in the brain that is at present largely cortico-centric^1,3,4,7,9–11,123–131^.

There are also some dissimilarities in the basic characteristics of brainstem-augmented structural and functional connectomes. For example, SC is generally stronger and has greater inter-subject similarity within brainstem and within cortex than between brainstem and cortex (Figs. 1d and S2a). By contrast, for FC, brainstem-cortex connectivity is neither the weakest nor the least consistent compartment across individuals (Fig. S2b). This divergence between SC and FC behavior may reflect both biological and methodological factors. From the biological standpoint, brainstem-cortex FC can be strong and relatively consistent, even when tractography-derived SC is weaker or less consistent. That is because FC is not only shaped by direct synaptic interactions but also by neuromodulatory influences, receptor architecture, and neurotransmitter release patterns^1,3,10,61,62,64,132,133^. At the same time, these differences may come down to the fact that SC and FC capture different features of brain architecture. SC measures direct physical connections that are subject to finite metabolic and spatial resources, whereas FC is a statistical construct that represents a multitude of direct and indirect interactions between regions and is not subject to the same constraints^130^. As a result, SC and FC depend on brain geometry and spatial embedding to different extents^134^. For example, physical distance has a stronger effect on diffusion MRI-derived SC estimates than on FC estimates.

Because brainstem nuclei and cortical parcels are generally separated by longer anatomical distances, brainstem-cortex SC may appear weaker and less consistent than SC within cortex or within brainstem. This distance-related effect is less dominant for FC (see Figs. S17 and S16). These differences in SC and FC also manifest in their weighted degree (strength) sequences. The correspondence between cortex-to-brainstem weighted degree estimated from SC and FC is modest (*ρ* = 0.31), yet comparable to the SC and FC weighted degree correspondence within cortex (*ρ* = 0.14) (Fig. S8). This modest correspondence is in line with prior edge-wise SC-FC coupling studies^135^, and is expected given the fundamental biological and methodological differences between SC and FC. As stated above, FC is not solely shaped by direct anatomical connections, rather it is influenced by neuromodulation and broader synaptic interactions^1,3,10,61,62,64,132,133^. Moreover, SC and FC differ in their sensitivity to basic parcel-level properties, including inter-regional distance and parcel volume. In all, treating parcel volume and inter-regional distance as covariates when examining brainstem-augmented SC and FC measures may remove the effects caused by these low-level properties.

Brainstem-augmented structural connectomes hold promise for developing computational biophysical models that simulate brain dynamics, and for understanding the anatomical foundations of cortical functional organization^136–140^. In such models, the structural connectome provides the anatomical scaffold that couples local neural population dynamics across distributed brain regions and constrains the propagation of neuronal activity. However, current models typically ignore the contribution of brainstem connectivity to simulated dynamics, despite the fact that brainstem nuclei exert considerable modulatory influence on whole-brain dynamics via neurotransmitters^24,29,69,141–146^. These neuromodulators are transmitted via long-range axonal projections to distributed cortical and subcortical brain regions, where they excite, inhibit or change the gain of post-synaptic neurons^80,147^. These adjustments in post-synaptic activity alter how signals are routed through the structural network of the brain, and ultimately how they are integrated^10^. We therefore envisage that increasingly accurate and biologically grounded modeling of neural dynamics will benefit from inclusion of widespread and heterogeneous structural connectivity profiles of brain-stem nuclei into biophysical models.

In a similar vein, brainstem-augmented structural connectomes provide a field of view beyond the cerebral cortex for investigating network spreading mechanisms in neurodegenerative diseases^148^. A growing body of evidence supports the existence, aggregation and region-to-region propagation of misfolded proteins in diseases such as Parkinson’s disease (PD)^149–153^, multiple system atrophy (MSA)^154,155^, Alzheimer’s disease (AD)^156,157^, frontotemporal dementia (FTD)^158^, and amyotrophic lateral sclerosis (ALS)^159,160^. Prion-like spreading of misfolded proteins along anatomical connections facilitates the spread of pathology from initial disease epicenters to anatomically distal regions^161–165^. Recently-developed models of disease pathological spread, such as susceptible–infected–removed^150,158,160,166^ , stochastic epidemic spreading models^167^, and diffusion models^168^ use structural connectomes to explain and forecast *in vivo* disease pathology^169^. Postmortem studies have shown that the brainstem is among the most severely affected structures in neurodegenerative diseases such as PD and ALS, and exhibits substantial amounts of misfolded-protein aggregates^170–175^. Incorporating brainstem-augmented structural connectomes into spreading models should enable more biologically realistic simulations of disease progression and provide a framework for understanding how brainstem nuclei and their anatomical connections shape pathology-specific trajectories across disease stages. Even in primarily supratentorial-dominated diseases, such as AD and FTD, brainstem-augmented connectomes could enable the identification of when and which brainstem nuclei become vulnerable based on their anatomical connectivity profiles^176,177^.

The present work should be considered alongside several methodological considerations. First, both structural and functional imaging of the brainstem are vulnerable to physiological noise arising from nearby pulsatile vasculature and cerebrospinal fluid filled spaces. Low signal-to-noise ratio increases uncertainty in fiber orientation estimation^115,178^ that can potentially bias structural connectivity estimates. Furthermore, low signal-to-noise ratio can contaminate BOLD signals with non-neuronal fluctuations, complicating the interpretation of brainstem functional connectivity^18^. We attempted to ameliorate this using extensive physiological noise correction. The densely packed nature of brainstem nuclei organization can also make their precise localization particularly difficult. Creating subject-specific brainstem nuclei labels from dedicated high-resolution multi-contrast 7 Tesla MRI of each subject would potentially enable more precise localization of nuclei, and a more thorough characterization of their structural and functional connectivity.

Second, fiber tracking models applied to diffusion-weighted images may generate systematic errors when reconstructing complex physical fiber configurations, including crossing, bending, and fanning fibers. Tractography methods preferentially reconstruct large fiber bundles with relatively straight and short trajectories, while under-representing more complex geometries^115,178–180^. This issue is particularly predominant in the brainstem, where fiber bundles are densely packed. To address this limitation we incorporated probabilistic tracking with anatomical constraints (ACT) and filtering (SIFT), which mitigates the large fiber bundle bias and increases the biological plausibility of streamlines^181,182^. Both limitations illustrate that more research is necessary to optimize tractometry in the brainstem^48,98^. A promising future direction would be to map the spatial distribution of streamline endpoints from individual brainstem nuclei and compare them against known anatomical connectivity constraints, following the approach of Dalton et al.^183^, which would provide a more direct validation of the reconstructed pathways.

Third, brainstem nuclei with larger parcel volumes, or the nuclei that are physically close to one another, tend to have stronger structural connectivity. An important question is whether this effect reflects genuine biological factors (larger neuronal populations giving rise to more axonal projections) or methodological factors (improved tractography performance in larger nuclei due to increased signal stability). We controlled for the effect of spatial proximity using null models that preserve edge length, and for the effect of size by statistically controlling for nucleus volume.

The fourth limitation is partial-voluming inherent to MR imaging, and this effect is especially problematic for small and closely adjacent brainstem nuclei^184,185^. A subset of brainstem nuclei included in the Brainstem Navigator atlas, such as the median raphe (14 ± 2 mm^3^), raphe magnus (16 ± 3 mm^3^), locus coeruleus (∼12 mm^3^ per hemisphere), superior olivary complex (16-20 mm^3^), right pedunculotegmental nucleus (24.8 ± 2.5 mm^3^), and parvicellular reticular nucleus alpha (∼19–20 mm^3^), are small relative to the 1.7 mm isotropic DWI voxel volume (∼ 4.91 mm^3^) used in this study for tractography (see Table. S4 for volumes of all 58 brainstem and 8 diencephalic nuclei compiled from the published brainstem atlas literature^38–44^). These nuclei are therefore more vulnerable to partial volume effects. However, the core findings of this study, the structure-function coupling in dopaminergic, serotonergic, and relay nuclei, are driven by the set of nuclei that are substantially larger and well-sampled at this resolution. Furthermore, we also included nucleus volume as a covariate in all regression models, ensuring that any bias introduced by differences in nucleus size does not affect our structure-function coupling results. Continued improvements in precision MRI and ultra-high-resolution scanning are needed for minimizing these effects and increasing the anatomical specificity of nucleus-level analyses.

In summary, there is growing effort in neuroscience to move beyond a cortico-centric framework toward a more integrative view of brain and central nervous system organization. The constellation of nuclei that constitute the brainstem is a focal point for numerous basal and cognitive functions, but they remain an underexplored domain of the wider brain connectome. The present work presents an initial step toward understanding how the brainstem features in the broader question of structure-function relationships in the brain.

## METHODS

### Diffusion MRI data acquisition

7 Tesla diffusion MRI data were acquired using a Siemens Magnetom scanner from 20 healthy volunteers (mean age ± SE: 29.35 ± 1.21 years; 10 males, 10 females). A custom-built volume transmit coil combined with a 32-channel receive coil was used to enhance sensitivity in deep brainstem nuclei relative to standard commercial coils^186^. To minimize head motion during scanning, participants were positioned supine with foam padding placed around the head and neck and were instructed to remain as still as possible throughout the acquisition. All participants provided written informed consent in accordance with a protocol approved by the Institutional Review Board of Massachusetts General Hospital. For analysis, diffusion MRI data from 19 participants (mean age ± SE: 29.74 ± 1.21 years; 10 males, 9 females) were included. One participant was excluded due to poor signal sensitivity in the midbrain and thalamus, signal dropout, increased field inhomogeneities and spatial distortions in the anterior pons, and reduced fiber orientation distribution function (FOD) glyph amplitude compared with the remaining subjects. Full details of data acquisition and preprocessing are described in Garcia-Gomar *et al*. and Singh *et al*.^48,98^.

High-resolution (1.7 mm isotropic) diffusion-weighted imaging (DWI) data were acquired with a 2D single-shot spin-echo echo-planar imaging (EPI) approach, unipolar diffusion encoding with 82 contiguous slices to achieve full brain coverage, 66.8 ms echo time, 7.4 s repetition time, anterior/posterior phase encoding direction, 1 475 Hz/Px bandwidth, partial Fourier of 6*/*8, 60 diffusion directions, b value of 2 500 s/mm^2^, and acquisition time 8^*′*^53^*′′*^. Additionally, for distortion correction we acquired 7 interspersed “b0” images (T2-weighted, non-diffusion-weighted EPI, b value 0 s/mm^2^) with opposite phase-encoding direction, and matched geometric distortion and spatial resolution to the DWIs.

### Functional MRI data acquisition

Brainstem functional magnetic resonance imaging (fMRI) data were acquired, preprocessed, and previously presented by Cauzzo *et al*.^36^ and Singh *et al*.^37^. The study protocol was approved by the Institutional Review Board of Massachusetts General Hospital. All participants provided written informed consent in accordance with the Declaration of Helsinki. Twenty unrelated healthy participants (mean ± SE age: 29.35 ± 1.21 years; 10 males, 10 females) participated in an eyes-closed resting-state 7 Tesla session (Magnetom, Siemens Healthineers, Erlangen, Germany). Participants were compensated at a rate of 35 USD per hour. During the session, three 10–minute runs were acquired. Importantly, brainstem-specific custom protocols were developed for the 7 Tesla MRI acquisition and processing, which we describe below. Complete acquisition and processing details are provided in Cauzzo *et al*.^36^ and Singh *et al*.^37^.

Briefly, imaging was performed using a custom-built 32-channel receive coil and a volume transmit coil at 7 Tesla. For each participant, three runs of functional gradient-echo echo-planar imaging (EPI) data were collected with an isotropic voxel size of 1.1 mm, matrix size of 180 × 240, GRAPPA factor of 3, 123 interleaved slices (sagittal slice orientation), 0.82 ms echo-spacing time, 32 ms echo time, 2.5 s repetition time, flip angle of 75^°^, simultaneous-multi-slice factor of 3, 210 repetitions (volumes), anterior/posterior phase-encoding directions, 1 488 Hz/Px bandwidth, and acquisition time 10^*′*^07^*′′*^. Participants’ alertness was verbally confirmed between runs. Foam pads were used to minimize head motion and earplugs were provided. To account for physiology–related signal fluctuations, timing of cardiac and respiratory cycles was recorded using a piezoelectric finger pulse sensor (ADInstruments, Colorado Springs, CO, USA) and a piezoelectric respiratory bellow (UFI, Morro Bay, CA, USA). To correct for geometric distortion, a field map with 2.0 mm isotropic resolution was acquired. In addition, a T1-weighted multi-echo MEMPRAGE anatomical image was acquired for each participant using a 3 Tesla Siemens Connectom MRI scanner equipped with a custom-built volume transmit coil and a 64-channel receive coil. Images were obtained with an isotropic voxel size of 1 mm, field of view of 256 × 256 ×176 mm^3^, GRAPPA factor of 3, sagittal slice orientation, flip angle of 7^°^, echo time of 1.69, 3.5, 5.3, and 7.2 ms, 2.53 s repetition time, 1.5 s inversion-time, 650 Hz/Px bandwidth, anterior/posterior phase encoding direction, and acquisition time 4^*′*^28^*′′*36^.

### Diffusion MRI data preprocessing

DWIs were denoised using local principal component analysis filter^187^, motion and distortion corrected (FSL, topup/eddy). Diffusion tensor (FSL, dtifit) invariants, such as fractional anisotropy (FA) map and non-diffusion-weighted S0 image (T2-weighted), were then computed. To run the tractography, we mapped 58 brain-stem and 8 diencephalic nuclei from the Illinois Institute of Technology (IIT) MNI (IIT-MNI) space to native FA/S0 image space. To accomplish this, we first constructed an optimal group template from all subjects’ FA/S0 images in native S0 space using the Advanced Normalization Tool (ANTs)^188^; subsequently we coregistered the optimal FA/S0 group template to the FA/S0 template in IIT-MNI space using affine transformation followed by non-linear warp. Finally, the transformation matrices from those two steps (from single subject’s FA/S0 to optimal group template and from optimal group template to the FA/S0 template in IIT-MNI space) were concatenated and inverted to compute the full coregistration transformation, which were applied to brainstem nuclei labels to bring them from IIT-MNI space to single subject native space^36,37^. The 400 cortical labels of Schaefer parcellation in MNI space were transformed to single-subject native space using the concatenated inverse transformation of the following steps aligning native space images to MNI space: native FA/S0 images to optimal FA/S0 group template to IIT-MNI to MNI. Subcortical regions of interest (i.e., bilateral thalamus proper, caudate, putamen, pallidum, amygdala, hippocampus, accumbens) were parcellated from the preprocessed MEMPRAGE images using Freesurfer^189^. We employed FSL-based registration (FLIRT boundary-based affine registration) to bring these subcortical parcellations to native S0 space. Finally, the hypothalamic atlas label from Pauli *et al*.^190^ was registered from MNI space to native S0 space by concatenating the transformation from MNI space to IIT-MNI space to the inverse registration described above mapping images from IIT-MNI space to native space.

### Functional MRI data preprocessing

Physiological noise correction was applied separately to each resting-state fMRI run using a custom-built Matlab implementation of RETROICOR^35^ adapted to the slice acquisition sequence. Functional images were subsequently slice-time corrected, reoriented to a standard orientation, and coregistered to the MEMPRAGE anatomical image. Coregistration was implemented in AFNI using a two-step approach consisting of an affine alignment and a boundary-based, edge–enhancing nonlinear registration^191^. The accuracy of fMRI alignment to MNI template space was visually inspected for each participant to ensure that template-defined brainstem nuclei were well aligned with individual anatomy. Next, multiple nuisance regressors were regressed out from the fMRI time series, including six regressors describing rigid-body motion, a regressor describing respiratory volume per unit time convolved with a respiration response function^192^, a regressor describing heart rate convolved with a cardiac response function^193^, and five regressors capturing cerebrospinal fluid (CSF) signal. CSF regressors were derived using principal component analysis applied to a mask of the brainstem-surrounding ventricles, specifically, the lower part of the third ventricle, the cerebral aqueduct, and the fourth ventricle. Therefore, by design, the time-series of brainstem nuclei are not correlated with the signal from the surrounding ventricles^194,195^. The denoised data were then converted to percent signal change by dividing by the temporal signal mean, multiplying by 100, and bandpass filtering between 0.01–0.1 Hz. Finally, any residual temporal mean was removed, and the three runs were concatenated for subsequent functional network reconstruction.

### Brainstem nuclei segmentation

Brainstem nuclei were defined according to the Brainstem Navigator, a previously developed probabilistic atlas of *in vivo* brainstem (*n* = 58, 8 midline, 50 bilateral) and diencephalic (*n* = 8, 4 bilateral) nuclei^38^. The segmentation of the hypothalamus (*n* = 1 midline region) was from Pauli et al.^[190]^. All nuclei are defined in MNI152NLin6Sym space at 1–mm^3^ spatial resolution (matrix size: 182 × 218 × 182). Here we provide a brief overview of the atlas reconstruction; full methodological details are available in previous publications^38–44^. Briefly, 12 healthy participants (mean ± SE age: 28 ± 1 years; 6 males, 6 females) underwent 7 Tesla MRI imaging, during which a T2-weighted image and a diffusion-weighted image were acquired. Diffusion data were used to compute voxel-wise diffusion fractional anisotropy (FA). The T2-weighted and FA images have high contrast for brain-stem nuclei and were used to segment the nuclei. In Bianciardi *et al*.^38^, three raphe nuclei (median raphe, dorsal raphe, raphe magnus), the periaqueductal gray, the substantia nigra, and the red nuclei were segmented. For each nucleus, the segmentation was done using a semi-automated approach by clustering either the FA or the T2-weighted image. Identified clusters corresponded to an individual nucleus or, in some cases, corresponded to multiple nuclei. In cases where the identified cluster included several nuclei, manual separation of nuclei was performed based on established anatomical brain-stem knowledge. This procedure resulted in a binary mask for each nucleus at the individual level. Subsequently, labels were aligned to MNI152 space, averaged across individuals, and converted to probabilistic maps, where a value of 100% indicated that a voxel was consistently identified as belonging to a given nucleus across all 12 individuals. For all nuclei in the Brainstem Navigator atlas, the semi-automatic and manual segmentations ensured that no nuclei overlapped; however, slight overlaps may occur in the group-averaged probabilistic templates, depending on the probability threshold applied.

Later in Bianciardi *et al*.^39^, the mesopontine tegmental nuclei, including cuneiform, pedunculotegmental nuclei, oral pontine reticular nuclei, paramedian raphe, and caudal-linear raphe were segmented. They used the same semi-automated approach, with two additional validation steps: (1) nuclei were also manually segmented by a neurosurgeon using the T2-weighted and FA images, and the mesopontine tegmental anatomical landmarks; and (2) segmented nuclei were compared with a post-mortem histologically-defined atlas^196^. All subsequent nuclei segmented in later publications were delineated manually by multiple experts. Final individual-level labels were defined as the intersection of independent segmentations provided by the experts. The probabilistic templates were generated by averaging the segmentations across participants. Segmentations were validated against the Paxinos histological atlas^196^. Specifically, in Garcia-Gomar *et al*.^40^, the inferior and superior colliculi and superior olivary complex, as well as the medial geniculate nucleus (MGN) and lateral geniculate nucleus (LGN) which are part of the thalamus, were segmented. In Singh *et al*.^42^, the lateral and medial parabrachial nuclei, the vestibular nuclei complex, and the medullary viscero-sensory-motor nuclei were segmented. The lateral and medial parabrachial nuclei were further validated using histological evaluation from a post-mortem brainstem specimen. In Singh *et al*.^43^, the mesencephalic reticular formation, isthmic reticular formation, microcellular tegmental nucleus, ventral tegmental area, and the caudal-rostral linear raphe nucleus complex were segmented. Last, in Garcia-Gomar *et al*.^41^, the raphe obscurus, raphe pallidus, locus coeruleus, subcoeruleus, laterodorsal tegmental nucleus-central gray of the rhombencephalon, inferior and superior medullary reticular formation, and the pontine reticular nucleus (oral/caudal part) were segmented.

### Structural network reconstruction

To construct a structural connectome for each participant, we computed global tractography using refined tissue segmentation to precisely account for brainstem gray and white matter boundaries.

#### Tissue segmentations and GM-WM boundary mask

Tissue segmentations of cortical/subcortical gray matter (CGM/SGM), white matter (WM) and cerebrospinal fluid (CSF) were used as input for global tractography. To compute them, first we coregistered the preprocessed MEMPRAGE to native DWI S0 space using “mrtransform” of MRtrix3 software package (http://www.mrtrix.org)^197,198^. Then, we performed tissue segmentation into five different tissue types (5TT, i.e., CGM, SGM, WM, CSF and a pathological tissue compartment -empty in controls) of the MEMPRAGE image in native space using “5ttgen” algorithm in MRtrix3 with the FSL backend. We used non-skull-stripped and uncropped MEMPRAGE images to ensure accurate segmentation and preserve alignment with diffusion space. The 5ttgen algorithm treats brainstem and diencephalic nuclei mostly as white matter (because in T1-weighted MRI they are not clearly visible), thus the five tissue segmentations were refined to precisely account for brainstem and diencephalic gray matter as follows. In IIT space, we first defined a brainstem and diencephalic GM mask including all the Brainstem Navigator nuclei. We then coregistered the brainstem and diencephalic GM mask and the whole brainstem masks from IIT to S0 native space. The whole brainstem mask was then expanded by computing the union with the brainstem and diencephalic GM mask to include all Brainstem Navigator nuclei boundary voxels. The brainstem WM mask was computed at the native diffusion space by subtracting the brainstem and diencephalic GM mask from the expanded whole brainstem mask. Voxels overlapping with the CSF compartment were removed from the brainstem and diencephalic GM mask, brainstem WM mask, and the whole brainstem mask in native space to avoid misclassification. The CGM segmentation was then refined by removing all Brainstem Navigator nuclei voxels, including both GM and WM, using the expanded whole brainstem mask to ensure no brainstem voxels remained in the CGM compartment. Rather, this brainstem and diencephalic GM mask was incorporated into the SGM segmentation, after enforcing mutual exclusivity with the existing SGM mask to prevent overlapping assignments. Further, the brainstem and diencephalic GM mask was removed from the WM segmentation, and the brainstem WM mask was added to the WM segmentation. Finally, to prevent any overlap between the WM and the updated SGM segmentation, the SGM mask was subtracted from the WM segmentation. The CSF and pathological tissue compartments were left unchanged. Finally, the refined CGM, SGM, WM segmentations along with CSF and pathological tissue segmentations were merged to generate a refined 5TT image in S0 native diffusion space, input for anatomically constrained tractography.

The gray-white matter interface mask was generated from the refined 5TT image in diffusion space using the 5tt2gmwmi command in MRtrix3. This step incorporated both modified CGM and SGM compartments defined in the refined 5TT image to accurately delineate the gray-white matter interface, which was subsequently used for anatomically accurate streamline seeding in anatomically constrained tractography.

#### Global tractography

Global tractography was performed using MRtrix3 (http://www.mrtrix.org)^197,198^. Voxel-wise white matter fiber orientation distributions (FODs) were estimated in diffusion native space via constrained spherical deconvolution (dwi2fod) after computing the response function from preprocessed DWIs (dwi2response)^199,200^. FODs, together with the refined 5TT segmentation and gray-white matter interface (see above), served as input for anatomically constrained tractography (ACT) using the iFOD2 algorithm (“tckgen”)^201^. Streamlines were seeded at the gray-white matter interface to ensure biologically plausible fiber initiation, and backtracking was enabled to enforce anatomically valid termination at tissue boundaries. We generated 100 million total streamlines with minimum streamline length of 1 mm, maximum angle between successive steps of 45°, and an FOD amplitude cut-off of 0.07. Subsequently, we applied “tcksift” in MRtrix3 to perform spherical deconvolution-informed filtering of tractograms (SIFT)^182^. This combined framework is hereafter referred to as ACT-SIFT-based tractography. The refined 5TT image was provided as the anatomical constraint, and SIFT was performed with reference to the FOD image. Streamlines were iteratively removed to optimize correspondence between streamline density and fiber density estimates until a final tractogram comprising 10 million streamlines was obtained for downstream analyses.

Following ACT-SIFT-based tractography, region-wise structural connectivity was quantified using an endpoint-constrained, seed-target approach. For each subject, streamline subsets were extracted from the ACT-SIFT tractograms (10 million streamlines) using “tckedit” with the -include and -ends_only options. This procedure was performed for each of the 483 predefined regions registered to diffusion native space. Only streamlines with endpoints terminating within a given region were retained, while streamlines merely passing through the region were excluded. For each seed region, streamlines terminating in every other target region were subsequently isolated using the same endpoint-restricted criteria. The number of streamlines connecting each seed-target region pair was quantified using “tckinfo” and later normalized by 10 million as the total number of streamlines in SIFT, and streamline length statistics (mean, standard deviation, median, maximum, and minimum) were computed using “tckstats”. This procedure was repeated for all seed-target combinations, resulting in a subject-specific 483 × 483 structural connectivity matrix derived from the ACT-SIFT tractograms.

### Functional network reconstruction

To construct a functional connectome for each participant, we parcellated the cortex using the 400-region Schaefer atlas^53^ and parcellated the brainstem using the 58-nucleus Brainstem Navigator atlas (https://www.nitrc.org/projects/brainstemnavig^38–44^). In all analyses, we used the probabilistic brainstem labels that were thresholded at 35%. This thresholding resulted in some overlapping voxels for bordering nuclei. Specifically, in the 8334 voxels labeled as part of a brainstem nucleus, 7922 (95%) were labeled only once, whereas 405 (4.8%) were labeled twice and 7 (0.08%) labeled three times. No voxel was assigned to more than three nuclei. Functional connectivity was quantified as the Pearson correlation between time-series for every pair of brain regions (458 regions in total). The group-averaged functional connectome was obtained by averaging individual connectomes across participants.

### Group-consensus structural connectivity

The group-representative structural connectome (shown in Fig. 1c) was constructed using a distance-dependent consensus thresholding approach^57,202,203^. This approach identified consistent structural connections across subjects and mitigated inconsistencies in individual subject tractography reconstructions^204,205^. To construct the group-consensus connectome (as shown in Fig. 1c), we first derived a binary group-consensus structural connectivity matrix while preserving the connectomes’ (1) density and (2) distribution of edge lengths across individual subjects^57,121,206^. We collated all edges from individual subject connectomes and binned the edges by their length. The number of bins was determined heuristically, as the square root of the mean binary density across subjects. Within each bin, edges were ranked by their frequency of occurrence across subjects, and the most consistently expressed connections were retained. Specifically, if the mean number of edges across subjects within a given bin was equal to *k*, the *k* edges with the highest cross-subject consistency of that length were selected. The resulting binary consensus matrix was then applied as a mask to the element-wise average of individual subjects’ weighted structural connectivity matrices, yielding the final weighted group-consensus connectome. This distance-dependent thresholding procedure yielded a network that preserved the empirical edge-length distribution while preferentially preserving connections that were consistently expressed across subjects.

### Replication dataset: Parkinson’s Progression Markers Initiative (PPMI)

We used data from 25 healthy controls from the PPMI dataset (mean age ± SD: 66.34 ± 11.94 years; 15 males, 10 females) to reconstruct the brainstem-augmented structural and functional connectomes^83^.

DWI preprocessing was performed using MRtrix3^197,198^, FSL^207^, ANTs^188,208^, and FreeSurfer 7.3.3^209^, and included Marchenko-Pastur principal component analysis (MP-PCA) denoising, Rician bias correction, Gibbs-ringing removal, motion/eddy correction (FSL topup/eddy via dwifslpreproc), and N4 bias-field correction. The diffusion tensor and FA maps were fitted within a SynthStrip brain mask, and the mean B0/FA images were registered to the IIT atlas^210^ using ANTs bivariate deformable registration. The inverse warp propagated the Brainstem Navigator v0.9 brainstem nuclei atlas (58 brainstem nuclei, thresholded at 0.35) and the Schaefer-400 cortical parcellation into native DWI space.

The structural connectome was derived using single-shell single-tissue CSD and probabilistic tractography (iFOD2) with anatomically-constrained tractography (5TT, custom-corrected so brainstem/diencephalic nuclei were reassigned to subcortical gray matter) and SIFT filtering^182^ (100M streamlines generated, 10M retained), producing a symmetric 482 ×482 matrix (58 brainstem nuclei, 24 subcortical regions, 400 cortical regions). Raw streamline counts were normalized per subject to counts-per-million, log1p-transformed, and harmonized across acquisition sites using ComBat (neuroCombat, Python)^211^, with site as the batch variable and age, sex, and session as covariates; mean-only ComBat was used for sites with fewer than two subjects. Functional connectome edges were the Fisher *z*-transformed Pearson correlations between mean regional BOLD time series, harmonized across sites with the same ComBat procedure and covariates. Further details on the PPMI dataset are available at https://www.ppmi-info.org.

### Structural communicability

Communicability (*C*_*ij*_) between two nodes *i* and *j* (*i* ≠*j*) quantified the extent to which they were connected through all possible weighted walks in a network, while accounting for both connection strength and path length^59,60^. Communicability incorporated contributions from walks of all lengths, assigning greater importance to shorter walks. For a weighted structural connectivity matrix *A*, communicability was defined using the matrix exponential of the degree-normalized adjacency matrix,

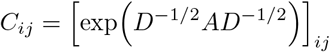

where *D* was the diagonal degree matrix with entries *D*_*ii*_ = ∑ _*k*_ *A*_*ik*_. In this formulation, each walk contributed the product of the edge weights along the walk, while normalization by node degree controlled for the influence of highly connected nodes and ensured that communicability reflected relative, rather than absolute, connectivity strength. Structural communicability was computed using the `communicability_wei` function from the `netneurotools` package^212^.

### Neurosynth meta-analytic term maps

Probabilistic measures of the association between voxels and cognitive processes were obtained from Neurosynth^55^, a meta-analytic tool that synthesizes results from more than 14 000 published neuroimaging studies by searching for high-frequency keywords that are published alongside their associated voxel coordinates (https://github.com/neurosynth/neurosynth, using the volumetric association test maps^56^). This measure of association was the probability that a given cognitive process was reported in the study if there was activation observed at a given voxel. Although 1 334 terms were reported in the Neurosynth engine, we focused our analysis primarily on cognitive functions and, therefore, limited the terms of interest to cognitive and behavioral terms. To avoid selection bias, the terms were selected from the Cognitive Atlas, a public ontology of cognitive science^74^, which included a comprehensive list of neurocognitive terms. We ended up incorporating 123 terms, ranging from umbrella terms (“attention” and “emotion”) to specific cognitive processes (“visual attention” and “episodic memory”), behaviors (“eating” and “sleep”) and emotional states (“fear” and “anxiety”). The coordinates reported by Neurosynth were parcellated according to the Schaefer-400 atlas^53^. The full list of Neurosynth functional/cognitive included terms is provided in Table. S2.

### Spin tests

To assess the effect of spatial autocorrelation on spatial associations between two cortical brain maps (Fig. 1g, 2), we used the so-called spatial autocorrelation preserving permutation tests, commonly referred to as “spin tests”^213^. Briefly, brain phenotypes were projected to spherical projection of the fsaverage surface. This involved selecting the coordinates of the vertex closest to the center of mass for each parcel. These parcel coordinates were then randomly rotated, and original parcels were reassigned to the value of the closest rotated parcel (*N* repetitions). For parcels where the medial wall was the closest, we assigned the value of the next closest parcel instead. Following these steps, we obtained a series of randomized brain maps that had the same values and spatial autocorrelation as the original map, while the relationship between values and their spatial location had been permuted. These maps were then used to generate null distributions of desired statistics. Throughout the manuscript, whenever the spatial correspondence between two brain maps was tested for, “spin test” was carried out for maps parcellated with the Schaefer-400 cortical atlas^53^. Notably, this parcellation was chosen because it divided the cortex into relatively homogeneous parcel sizes, and its number of parcels was comparable to the estimated number of distinct human neocortical areas^214,215^.

### Network randomization

Structural networks were randomized using a procedure that preserves the density, edge length distribution, and degree distribution of the empirical network^75,213^. Edges were binned according to Euclidean distance (10 bins). Within each bin, pairs of edges were selected at random and swapped, for a total number of swaps equal to the number of regions in the network multiplied by 20. This procedure was repeated *N* times to generate *N* null structural networks, which were then used to generate null distributions of network-related measure of interest.

When generating the null networks, we used Euclidean distance between brain regions as a proxy for streamline length. These measures are highly correlated (see Fig. S13). This choice was motivated by the sparsity of the empirical structural connectome, which results in a sparse length matrix that cannot be directly used in the rewiring algorithm introduced by Betzel *et al*.^75^.

## Data and code availability

All code used to perform the analyses is publicly available at https://github.com/netneurolab/Farahani_brainstemsc. The analyses rely on open-source Python packages, including NumPy (version 1.21.6)^216,217^, SciPy (version 1.7.3)^218^, pandas (version 1.3.5)^219^, seaborn (version 0.12.2)^220^, Matplotlib (version 3.5.3)^221^, statsmodels (version 0.13.5)^222^, Nilearn (version 0.10.1)^223^, NiBabel (version 4.0.2)^224^ and netneurotools (version 0.2.3)^212^.

Cortical regions are defined using the Schaefer-400 parcellation^53^. Brainstem nuclei are defined using the Brainstem Navigator atlas, which is publicly available at https://www.nitrc.org/projects/brainstemnavig^38–44^.

All preprocessed data, including the brainstem-augmented functional and structural connectomes are publicly available at https://osf.io/tqs64.

## Acknowledgments

BM acknowledges support from the Natural Sciences and Engineering Research Council of Canada (RGPIN-2017-04265), Canadian Institutes of Health Research (PJT-180439), and Canada Research Chairs Program (CRC-2022-00169). JYH acknowledges support from the Natural Sciences and Engineering Research Council of Canada. MB acknowledges support from the National Institute of Aging, NIH: R01-AG063982 and the Michael J. Fox Foundation: MJFF-022672 Award.

The funders had no role in study design, data collection and analysis, decision to publish or preparation of the manuscript.

## Disclaimer

The views expressed herein are those of the authors and do not reflect the official policy or position of the Henry M. Jackson Foundation for the Advancement of Military Medicine Inc.

The opinions and assertions expressed herein are those of the authors and do not reflect the official policy or position of the Uniformed Services University of the Health Sciences or the Department of War.

## Supplementary information

### Literature-based validation of selected brainstem connectivity profiles

#### Superior colliculus

The superior colliculus is involved in orienting the head and sensory systems toward objects of interest and is connected with structures that are related to visual, auditory, and somatosensory motor functions^225^. We observe the superior colliculus connectivity with the inferior colliculus, in agreement with the tract-tracing studies in guinea pigs and rhesus monkeys showing that the superior colliculus receives afferents from the inferior colliculus that contribute to the formation of an auditory map^226,227^. In line with a recent study in rodents, we also find superior colliculus connectivity with the para-bigeminal nucleus, this connection may have a role in defensive behavior^228^. Similarly, deep layers of the superior colliculus have been shown to connect to neurons in the periaqueductal gray in rats, as also observed in our data^229^. The superior colliculus also shows connectivity with the medial and isthmic reticular formations, which participate in horizontal saccades and head movements, as demonstrated in primate studies including rhesus and cynomolgus monkeys^226,230–233^. Consistent with the established role of the superior colliculus in visuomotor control and gaze shifts, we further observe connectivity with the visual cortex, supplementary eye fields, and thalamus, in agreement with evidence from mammalian and human studies^234–238^. We also observe superior colliculus connectivity with the substantia nigra reticulata, in agreement with anatomical evidence from mice, rats, and rhesus monkeys^226,238–241^. Additional superior colliculus connections are observed with the pedunculopontine tegmental nucleus, consistent with tract-tracing studies in rats, mice, and cats^231,242–244^. Superior colliculus connectivity with the cuneiform nucleus is also supported by tract-tracing evidence in rhesus monkeys and rats, which describes a tectocuneiform pathway from the superior colliculus to the cuneiform area, a pathway implicated in orienting and defensive-like motor responses^226,245,246^. Finally, anatomical connections between the hypothalamus and superior colliculus have been documented in mice and cats, and these pathways have been implicated in approach, appetitive, reproductive, and social attack behaviors^247,248^.

#### Ventral tegmental area with parabrachial pigmented nucleus of the ventral tegmental area nuclei complex

The ventral tegmental area with parabrachial pigmented nucleus of the ventral tegmental area nuclei complex (VTA-PBP) is involved in reward processing, motivation, cognition, drug addiction, avoidance behavior, fear conditioning, and arousal^249–253^. Here we observe VTA-PBP connectivity with prefrontal and orbitofrontal cortical areas, in line with previous findings in macaques and rodents^107,254–257^. We also observe connectivity with the pedunculotegmental nucleus and lateral parabrachial nucleus, consistent with anatomical studies in rats and squirrel monkeys^110,258–260^. In addition, prior rodent work has described anatomical connectivity between the VTA-PBP complex and the thalamus, periaqueductal gray, dorsal raphe, and hypothalamus^73,106,261–264^, which are shown to be its glutamatergic/cholinergic afferents. Furthermore, in rodents, VTA has been shown to have projections to amygdala^110,251,265^, raphe nuclei^110,266^, locus coeruleus^110,267^, mesencephalic and pontine reticular formation structures^266^, and hypothalamus^264,266,267^, subthalamic nucleus, striatal regions^263,268^, laterodorsal tegmental nucleus^73,263^ and lateral parabrachial nucleus^269^. Connections with many of these regions were also apparent in the VTA-PBP structural connectome presented in this work. In macaques, VTA connectivity to sensorimotor cortex, putamen, and caudate nucleus have also been reported^107^. VTA-PBP has also been shown to have connectivity to basal forebrain and ventral pallidum, which are GABAergic afferents of VTA^73^.

At the same time, we acknowledge that not all expected projections are recovered. In particular, based on established VTA projections to the nucleus accumbens in rat brain^110–112^, we expected to observe VTA–PBP connectivity with the accumbens, but this connection was not apparent in our tractography-derived connectome. This likely reflects known limitations of diffusion MRI tractography, including spatial resolution, crossing fibers, false negatives, and the difficulty of resolving small adjacent midbrain nuclei. Moreover, interpretation of fine-grained connectivity involving neighboring structures such as the substantia nigra and red nucleus remains challenging even in tract-tracing studies, where the contribution of anatomical adjacency versus true projection pathways can be difficult to disentangle.

#### Raphe pallidus

The raphe pallidus (RPa) is among the smallest raphe nuclei in the brain^69^ and is known to connect to autonomic, endocrine or somatic structures in the brain to enable its role in motor control, respiratory and sympathetic regulation. In line with our findings, RPa has been shown to have a widespread connectivity pattern throughout the brain, including connections to cortex, hypothalamus, and brainstem regions such as mesencephalic, pontine and medullary reticular formation, the parabrachial, the tegmental and other raphe nuclei, and spinal cord in rats^270^. Connectivity with substantia nigra and locus coeruleus, vestibular nuclei, paragigantocellular reticular nucleus, and amygdala are also reported in rat models^271–273^. Connections of raphe pallidus with periaqueductal gray are also reported in rats, however, we could not recover this connection in the data under study^272^. We further find connectivity with inferior olivary complex, ventral tegmental area, and subcortical regions such as pallidum, putamen, caudate, and sub-thalamic regions.

### Nucleus-level robustness across structural and functional measures

To provide a summary assessment of the reliability of nucleus-specific connectivity estimates, we ranked brain-stem nuclei using a composite score integrating nucleus volume, inter-subject similarity of SC and FC within the brainstem and between brainstem and cortex, and SC–FC correspondence within the brainstem and between brainstem and cortex (Fig. S24). This composite ranking provides a data-driven estimate of the relative robustness of individual nuclei and may help guide future nucleus-specific connectivity analyses.

---

**Figure S1.**
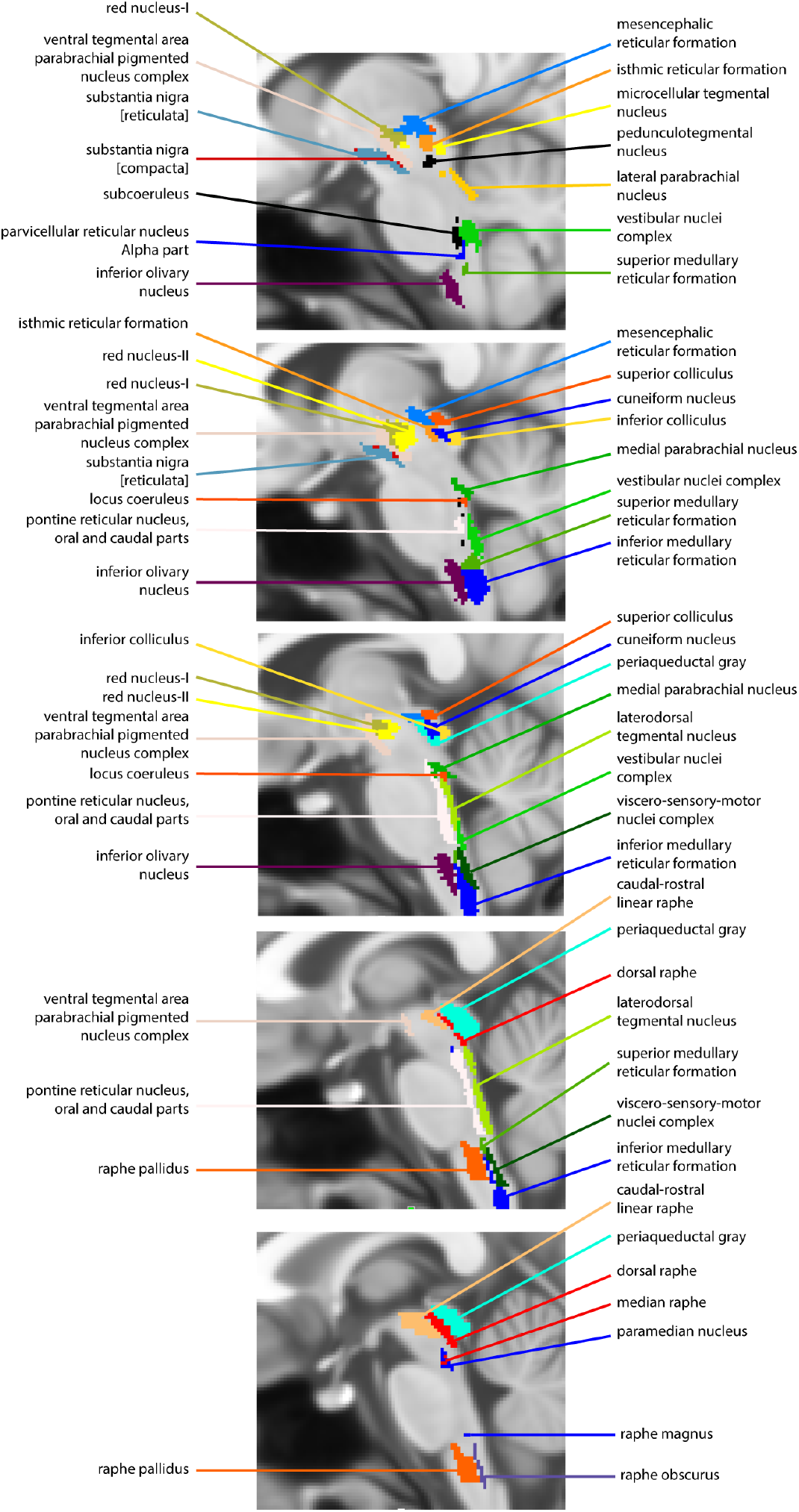
Representation of brainstem nuclei on the T1-weighted MNI template. Brainstem nuclei are displayed on the 1-mm MNI template. Each nucleus is labeled by name. Five slices encompassing the brainstem are shown to illustrate the spatial location of the nuclei across the template. The MNI template has a matrix size of 182 × 218 × 182 voxels, and the displayed slices correspond to P = 82, 84, 86, 88, and 90, C = 149, and A = 11.

**Figure S2.**
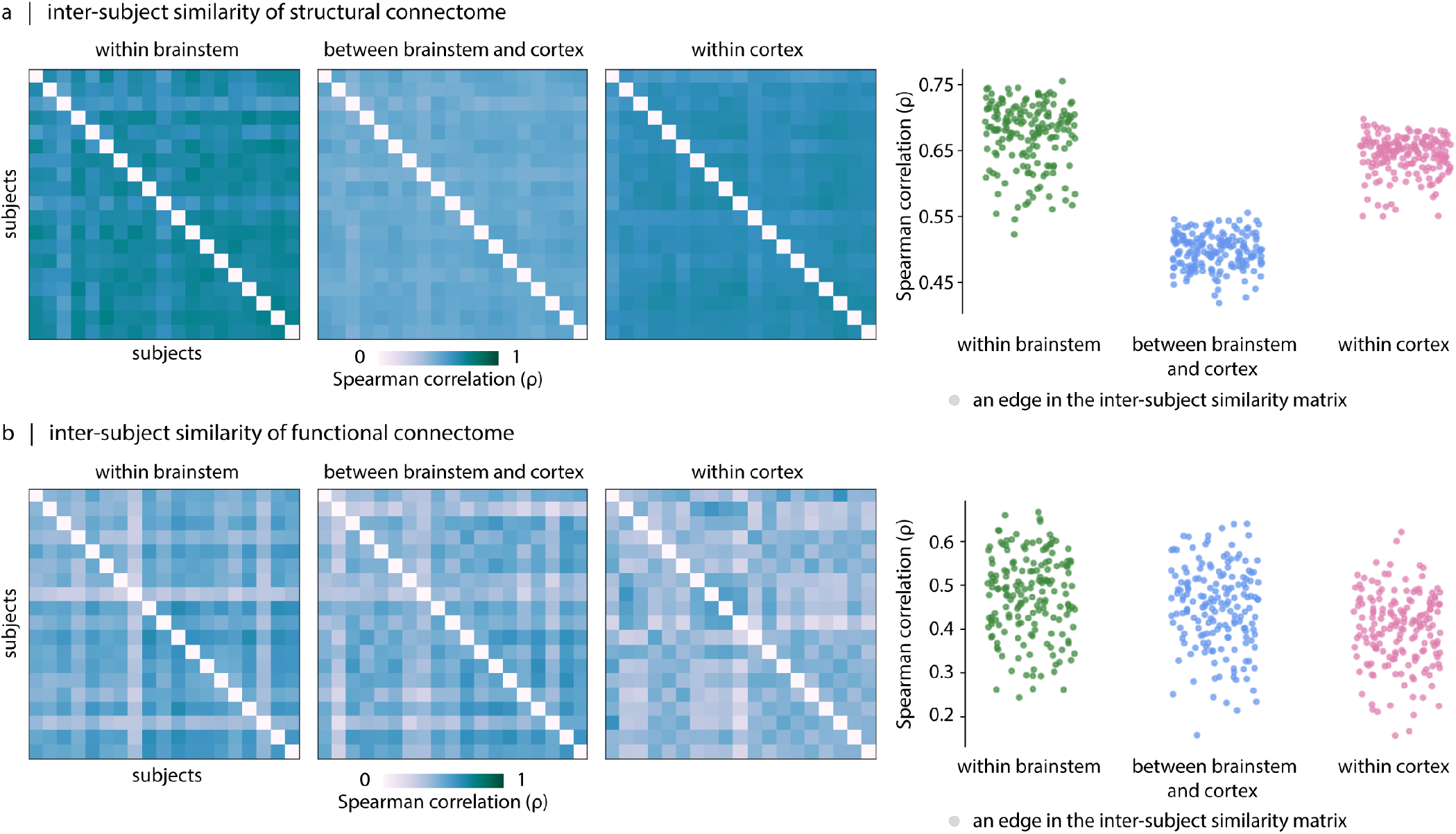
Inter-subject similarity of structural connectome (SC) and functional connectome (FC) Inter-subject similarity is quantified separately for SC and FC using Spearman correlation coefficients (*ρ*). For each pair of participants, we correlate (1) vectorized upper-triangular within-brainstem connectivity (2) vectorized brainstem-cortex connectivity, and (3) vectorized upper-triangular within-cortex connectivity. (a) The heatmaps show the resulting inter-subject similarity matrices computed from SC profiles. For SC, inter-subject similarity is higher for within-brainstem 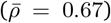 and within-cortex SC 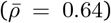 than for brainstem-cortex SC 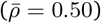 0.50). (b) For completeness, we further visualize the heatmaps corresponding to inter-subject similarity matrices computed from FC profiles. In this case, inter-subject similarity is broadly comparable across compartments with 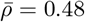 for within-brainstem, 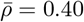 for within-cortex, and 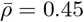 for brainstem-cortex compartment.

**Figure S3.**
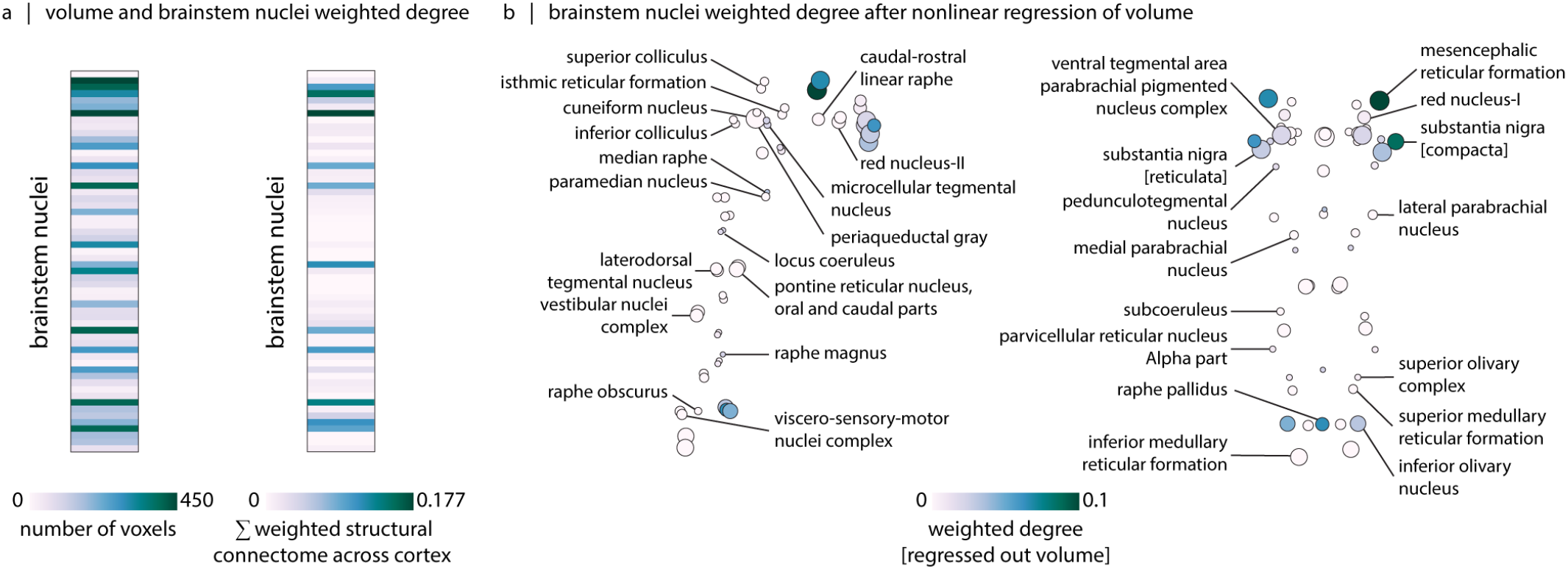
Brainstem-to-cortex connectivity after controlling for nucleus volume. After computing the weighted structural connectivity of each brainstem nucleus across all cortical regions (a, right), we regressed out the nonlinear effect of parcel volume (estimated by voxel count) (a, left) from the weighted degree. (b) Brainstem-to-cortex weighted degree after controlling for nucleus volume.

**Figure S4.**
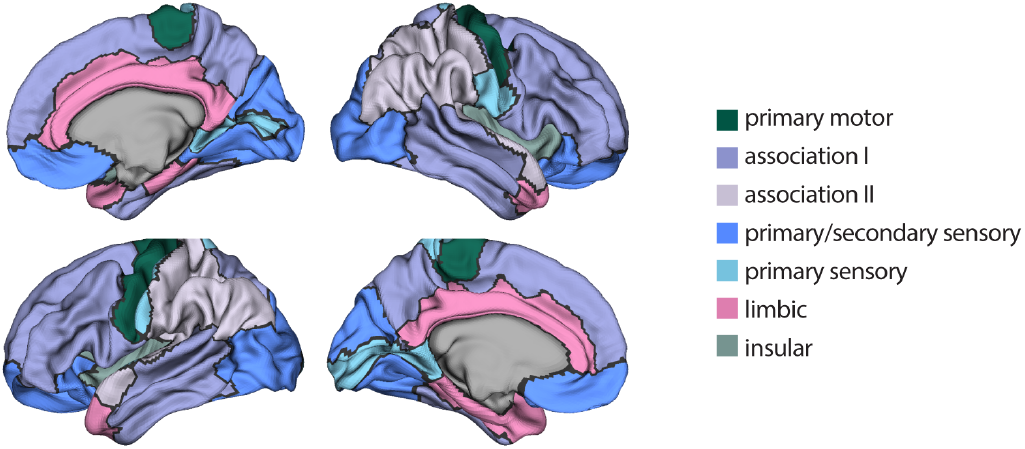
Von Economo cytoarchitectonic classes. The von Economo cytoarchitectonic parcellation^50–52^ is displayed on fs-LR midthickness cortical surfaces.

**Figure S5.**
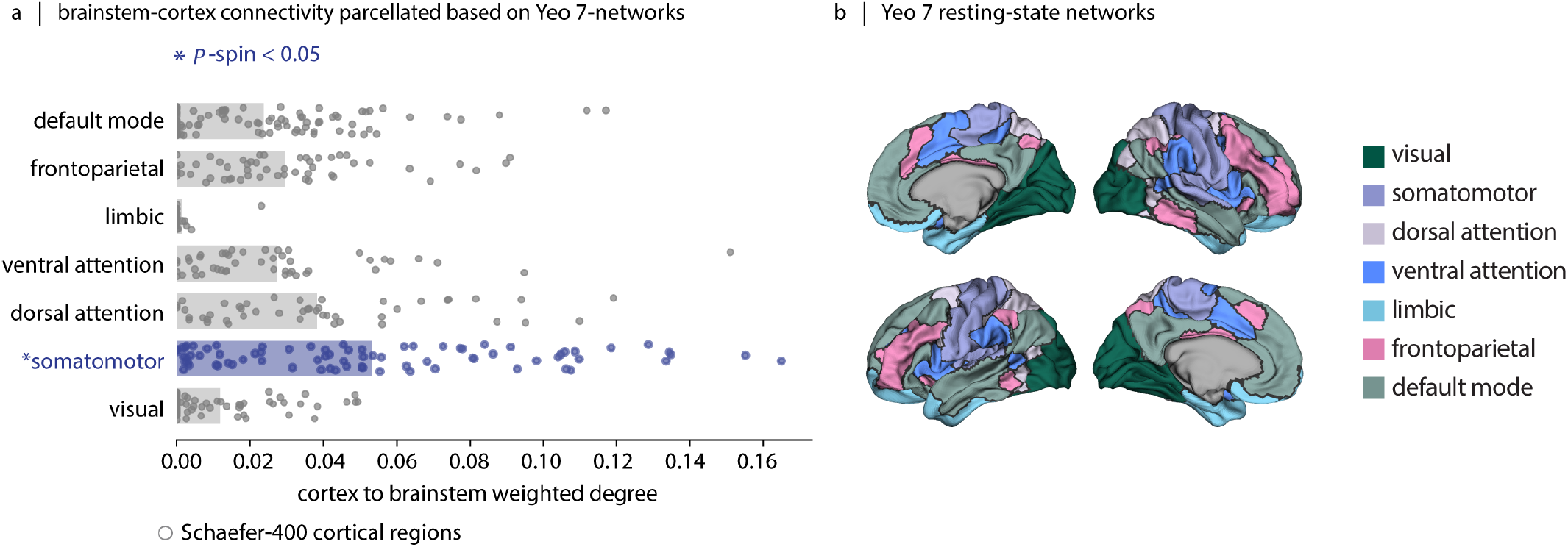
Cortex-to-brainstem weighted degree binned according to the Yeo 7-networks. (a) Each dot represents a cortical region defined according to the Schaefer-400 parcellation^53^. The brainstem shows strong structural connectivity with the somatomotor network (two-tailed, *p*_*spin*_ = 9.99 × 10^−3^, *N*_*spin*_ = 1 000). (b) The Yeo 7-networks parcellation is displayed on fs-LR midthickness cortical surfaces^54^.

**Figure S6.**
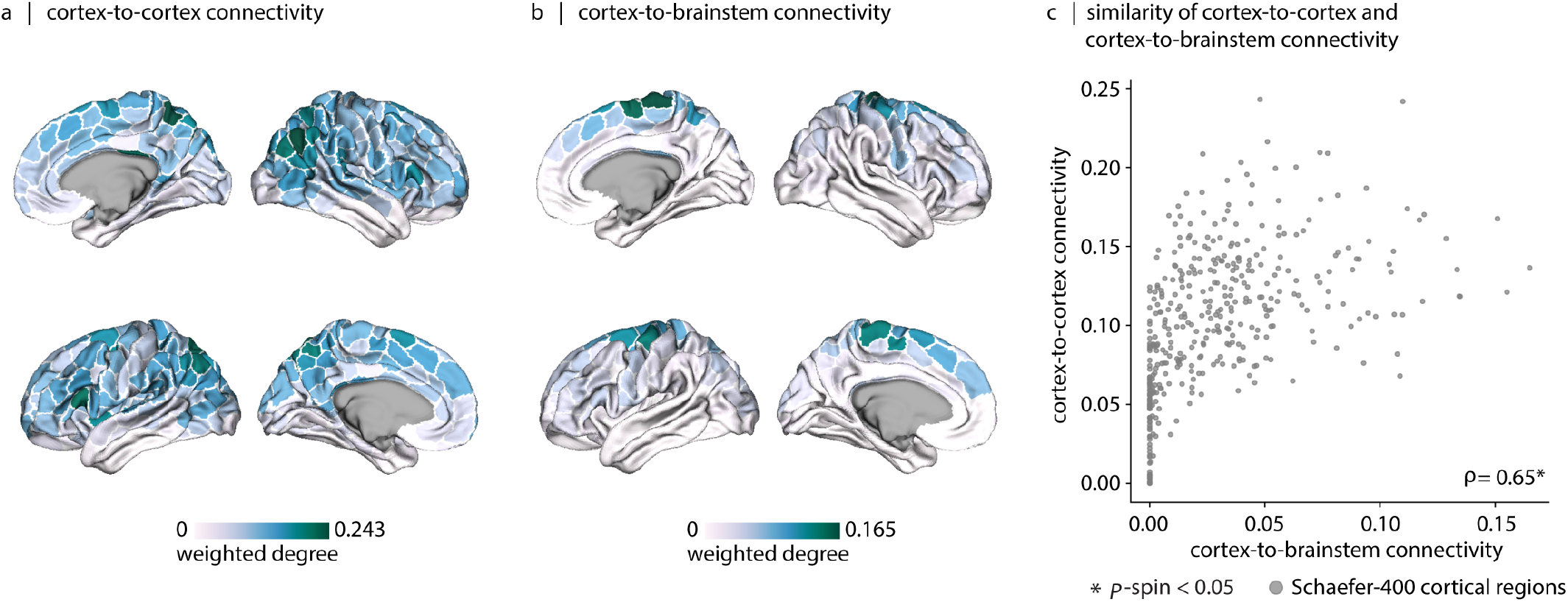
Similarity of cortex-to-cortex and cortex-to-brainstem structural connectivity weighted degree. (a) Cortex-to-cortex weighted degree. (b) Cortex-to-brainstem weighted degree. Brain cortical maps are displayed on fs-LR midthickness cortical surfaces. (c) The scatter plot shows the cortex-to-brainstem weighted degree on the *x*-axis and cortex-to-cortex weighted degree on the *y*-axis. The Spearman correlation between the two is equal to 0.65 (*p*_*spin*_ = 9.99 *×* 10^−4^, *N*_*spin*_ = 1 000).

**Figure S7.**
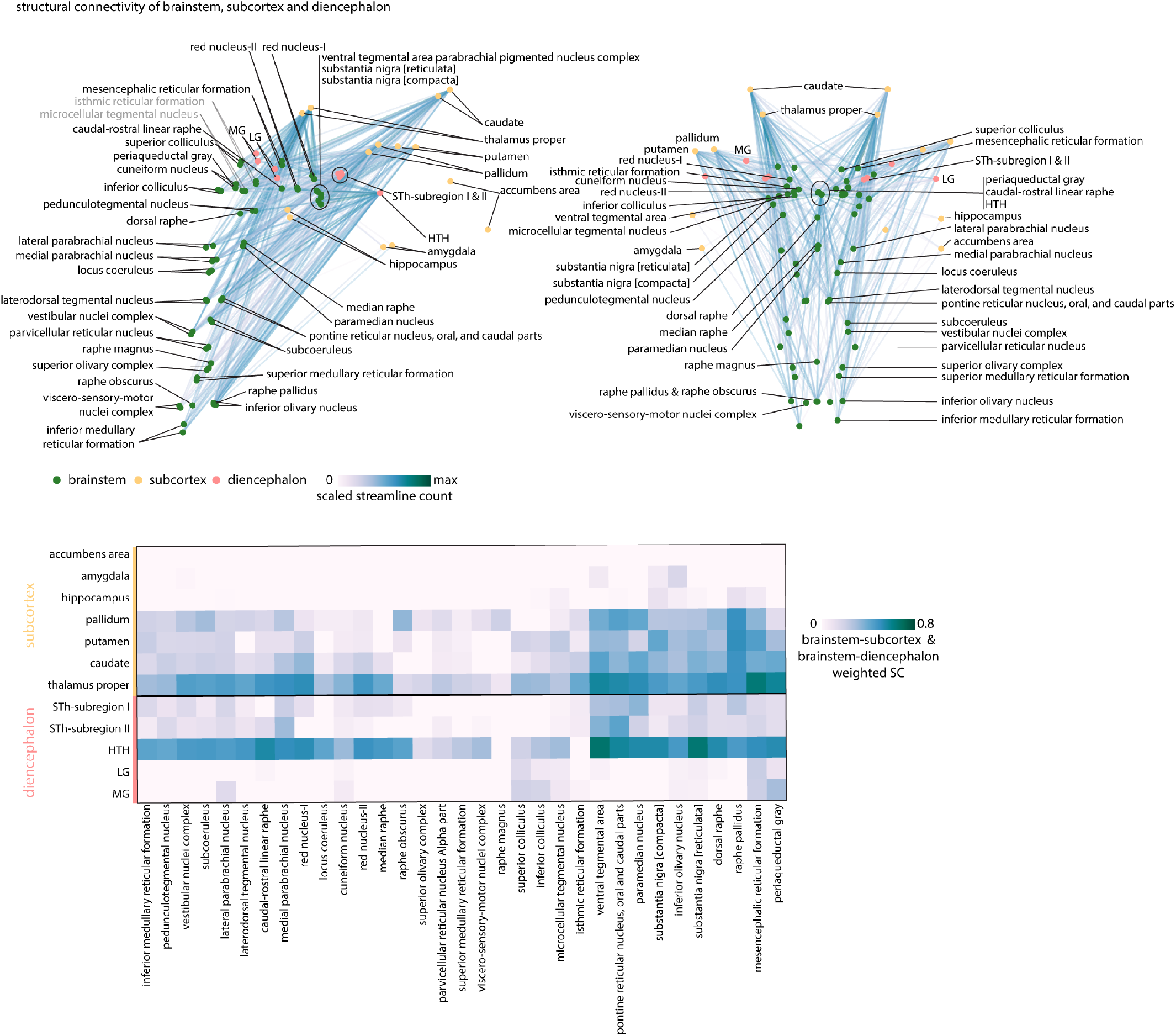
Connection profiles of brainstem nuclei with subcortex and diencephalon. Top: Sagittal and coronal views of brainstem, subcortical and diencephalon nuclei are shown. Edges show structural connections between brainstem nuclei and subcortical and diencephalic nuclei. Bottom: The heatmap shows the structural connectivity of brainstem nuclei with subcortical and diencephalic nuclei. For bilateral structures, connectivity values are averaged across the left and right hemispheres. Abbreviations used throughout this figure include STh: subthalamic nucleus, HTH: hypothalamus, LG: lateral geniculate nucleus, and MG: medial geniculate nucleus.

**Figure S8.**
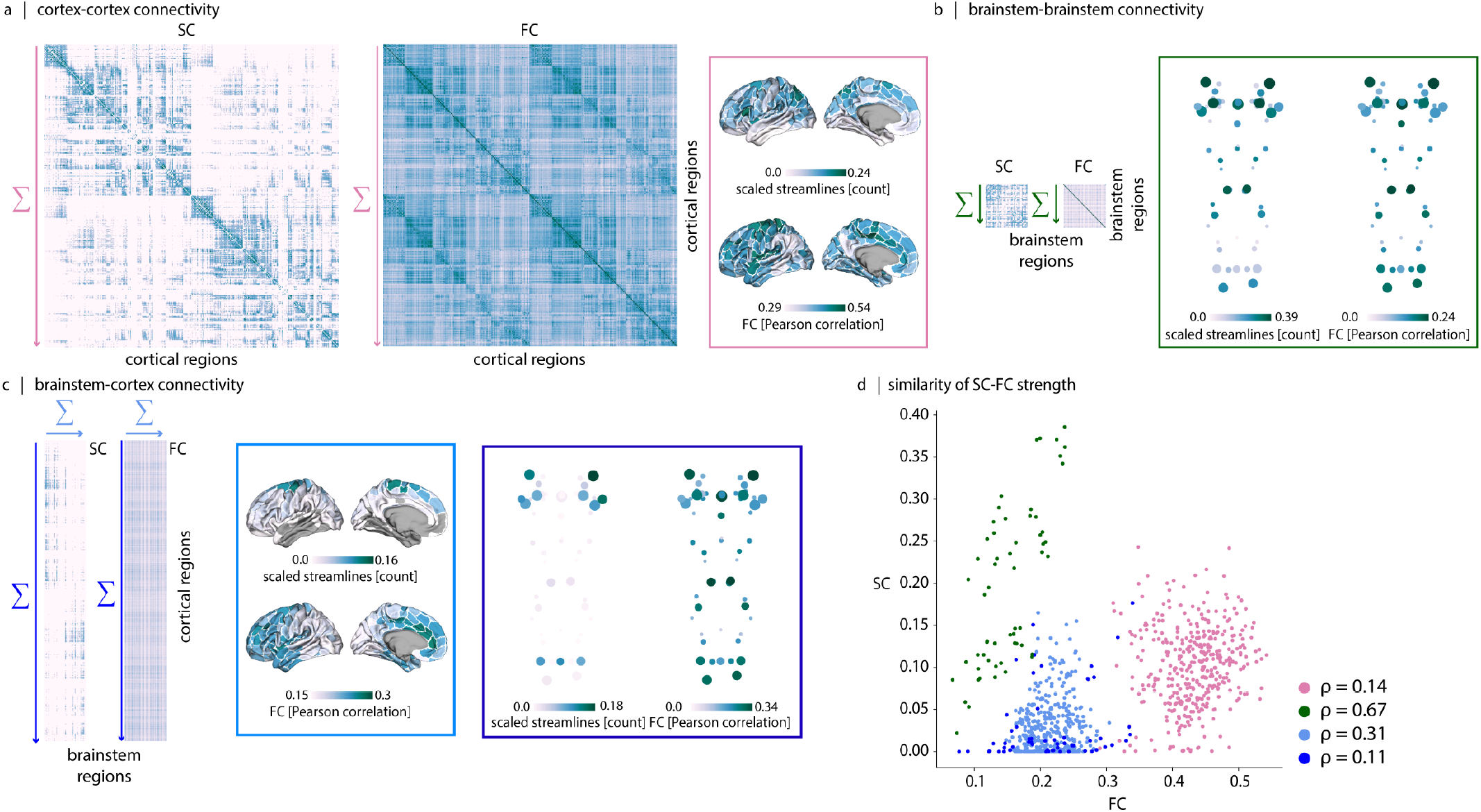
Structural and functional weighted degree patterns. (a) Cortex-to-cortex weighted degree is calculated by summing each cortical region’s SC/FC values across all cortical regions. (b) Brainstem-to-brainstem weighted degree is calculated by summing each brainstem nucleus’s SC/FC values across all brainstem nuclei. (c) Cortex-to-brainstem weighted degree is calculated by summing each cortical region’s SC/FC values across all brainstem nuclei. Brainstem-to-cortex weighted degree is calculated by summing each brainstem nucleus’s SC/FC values across all cortical regions. (d) Scatter plots show the correspondence between SC (*y*-axis) and FC (*x*-axis) weighted degree patterns across the different connectivity compartments.

**Figure S9.**
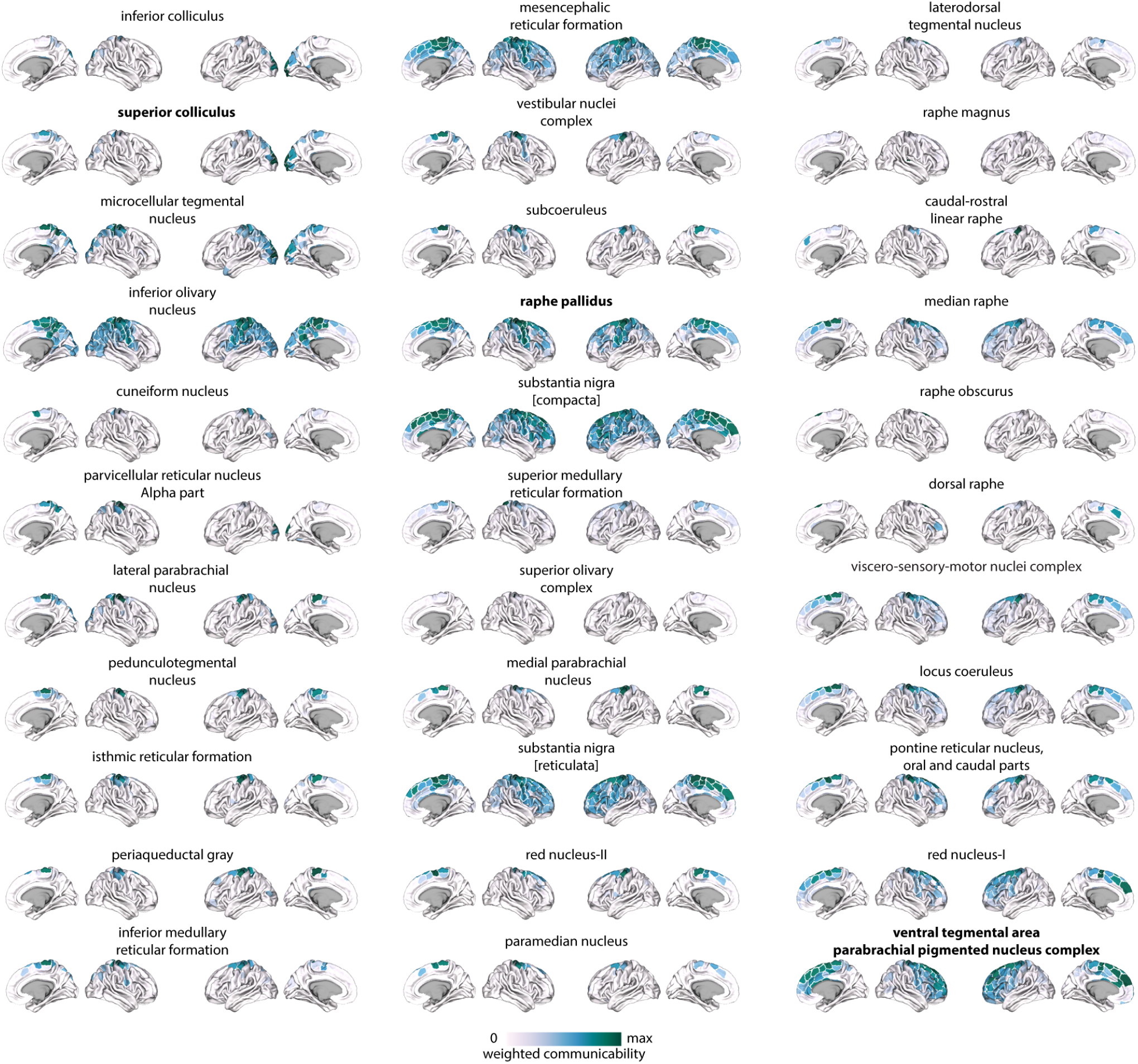
Connection profiles of all brainstem nuclei. Structural communicability profiles of each brainstem nucleus and cortex. Cortical maps are displayed on fs-LR midthickness surfaces. The connectivity patterns of the nuclei highlighted in bold are further shown in Fig. 2, and their anatomical correspondence with findings from histological and animal studies is further described in the *Supplementary information*.

**Figure S10.**
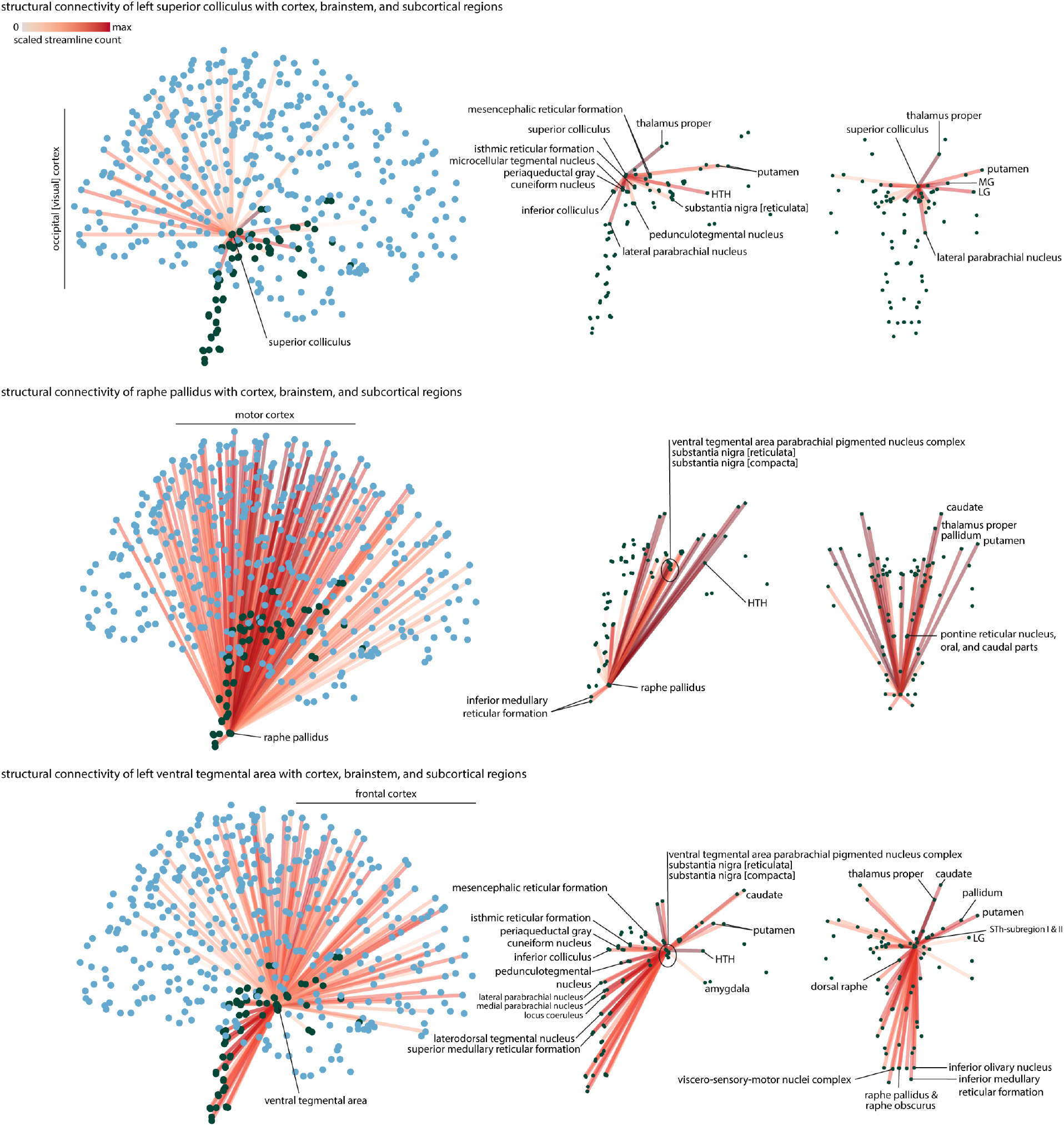
Structural connectivity profiles of superior colliculus, ventral tegmental area with parabrachial pigmented nucleus of the ventral tegmental area nuclei complex, and raphe pallidus. Edges represent reconstructed structural connections, with edge color indicating connection strength.

**Figure S11.**
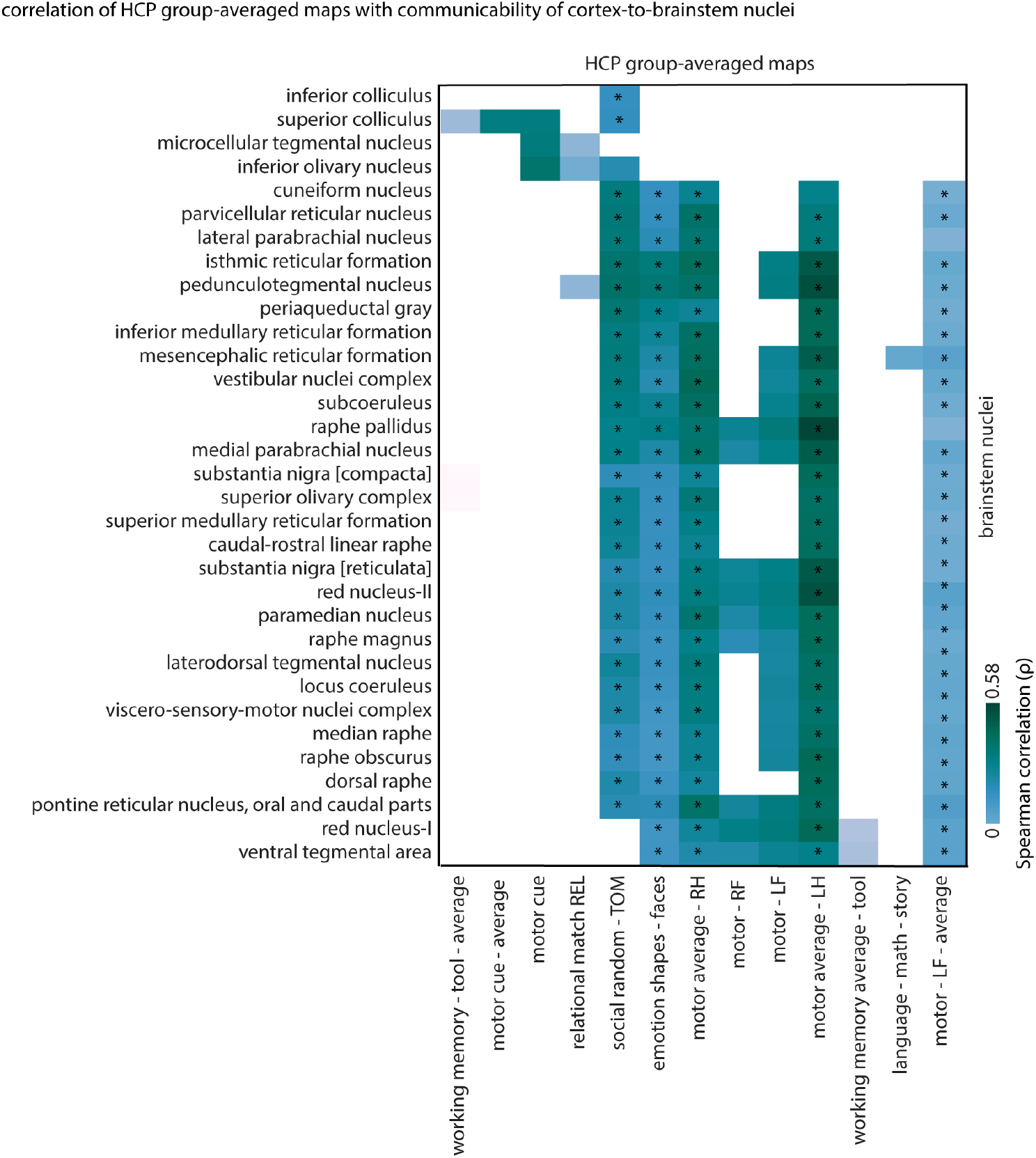
Correspondence between brainstem communicability profiles and HCP task-fMRI activation maps. The heatmap shows Spearman correlation values (*ρ*) between the structural communicability profile of each brainstem nucleus and cortical task-fMRI effect-size maps from the Human Connectome Project (HCP). HCP task-fMRI data include 86 contrasts across seven domains: working memory (WM), gambling, motor, language, social, relational, and emotion^274^. Redundant negative contrasts are excluded from the analysis, resulting in 63 task contrast maps included in this analysis. A list of all included contrast maps is provided in Table. S3. Contrasts for which at least one nucleus exhibited a positive correlation exceeding thresholds derived from spatial autocorrelation–preserving permutation nulls are shown on the *x*-axis (*p*_*spin*_ *<* 0.05, *N*_*spin*_ = 1 000). Superior colliculus connectivity is associated with the motor-cue contrast. The motor-cue contrast map is obtained by contrasting the rest period versus the cue period at the beginning of each motor task block, during which participants are visually instructed about the upcoming motor task. We also observe an association between inferior olivary connectivity and the contrast coming from the relational-processing task. This task requires participants to identify whether two pairs of objects share the same varying dimension (either shape or texture). This finding is consistent with prior evidence implicating the inferior olive in processing complex temporal sequences of visual stimuli and in eye-movement-related functions^275,276^. In addition, VTA and red nucleus connectivity profiles show associations with the higher-order working memory task. As expected, several brainstem nuclei also show associations with motor task contrasts. Note that we further apply False Discovery Rate (FDR) correction to the associations calculated per HCP-task; the associations that survive correction are indicated with an asterisk (*).

**Figure S12.**
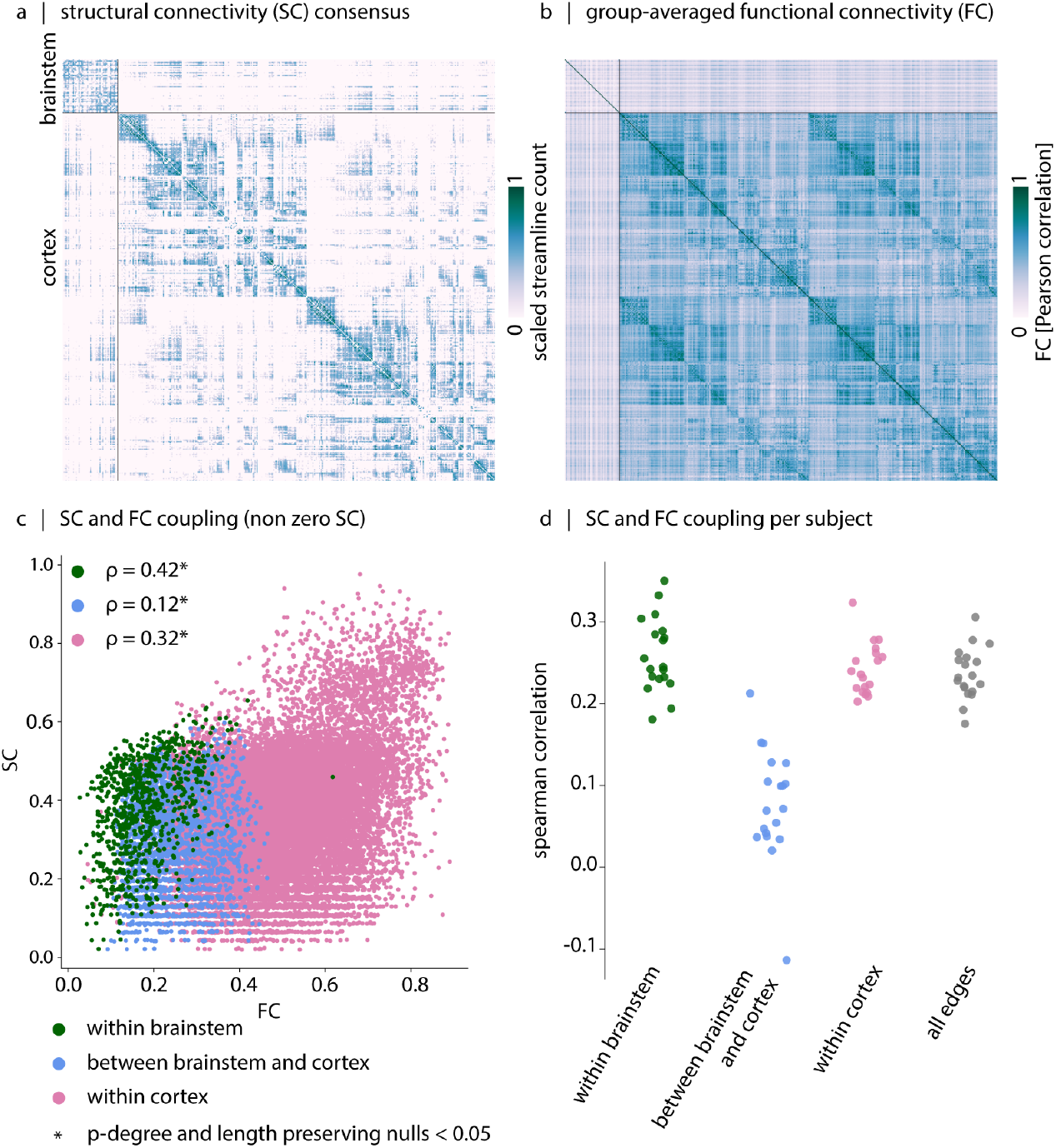
Association between structural and functional connectivity. (a) Group-consensus weighted structural connectivity (SC) matrix. (b) Group-averaged functional connectivity (FC) matrix. (c) The scatter plot shows the edge-wise relationship between SC and FC. Each dot represents a single edge. Spearman correlation coefficients are reported for each compartment (within-brainstem: *ρ* = 0.42, *p*_*rewired*_ = 9.99 × 10^−4^, *N*_*rewired*_ = 1 000; between brainstem and cortex: *ρ* = 0.12, *p*_*rewired*_ = 9.99 ×10^−4^, *N*_*rewired*_ = 1 000; and within-cortex: *ρ* = 0.32, *p*_*rewired*_ = 9.99 × 10^−4^, *N*_*rewired*_ = 1 000). Diagonal elements and edges with zero SC were excluded from correlation analyses. Reported *p*-values are obtained by comparing empirical structure-function correlations with a null distribution of correlations between degree- and length-preserving structural network nulls and empirical FC. Fisher’s *z* tests show that correlation between SC and FC is stronger within brainstem than between brainstem and cortex (*Z* = 9.884, *p ≈* 0), and it is also stronger than SC and FC correlation within cortex (*Z* = 3.461, *p* = 5.37 × 10^−4^). (d) Distribution of subject-level Spearman correlation coefficients between SC and FC, computed separately within each network compartment and across all edges.

**Figure S13.**
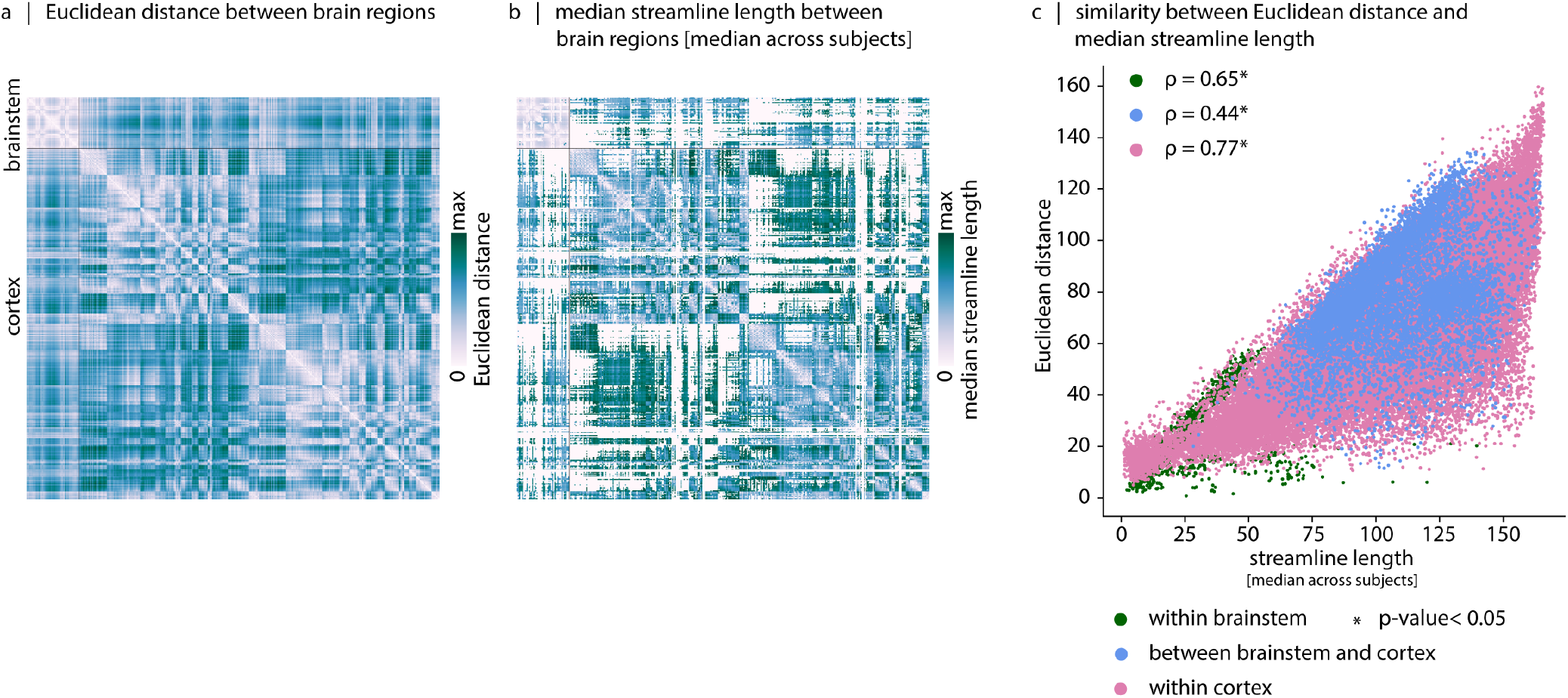
Similarity of Euclidean distance and streamline length between brain regions. (a) Euclidean distance between brain regions. (b) Streamline length between brain regions. For each edge, the median value across 19 individuals is shown. (c) The scatterplot shows the correspondence between Euclidean distance (*y*-axis) and streamline length (*x*-axis) across brain regions. Edges with a median streamline length of zero across individuals are excluded from the plot. We use the Euclidean distance matrix when generating degree- and edge length-preserving rewired structural networks^75^. This choice is motivated by sparsity of the streamline length matrix (panel b), which cannot be directly used in the rewiring algorithm.

**Figure S14.**
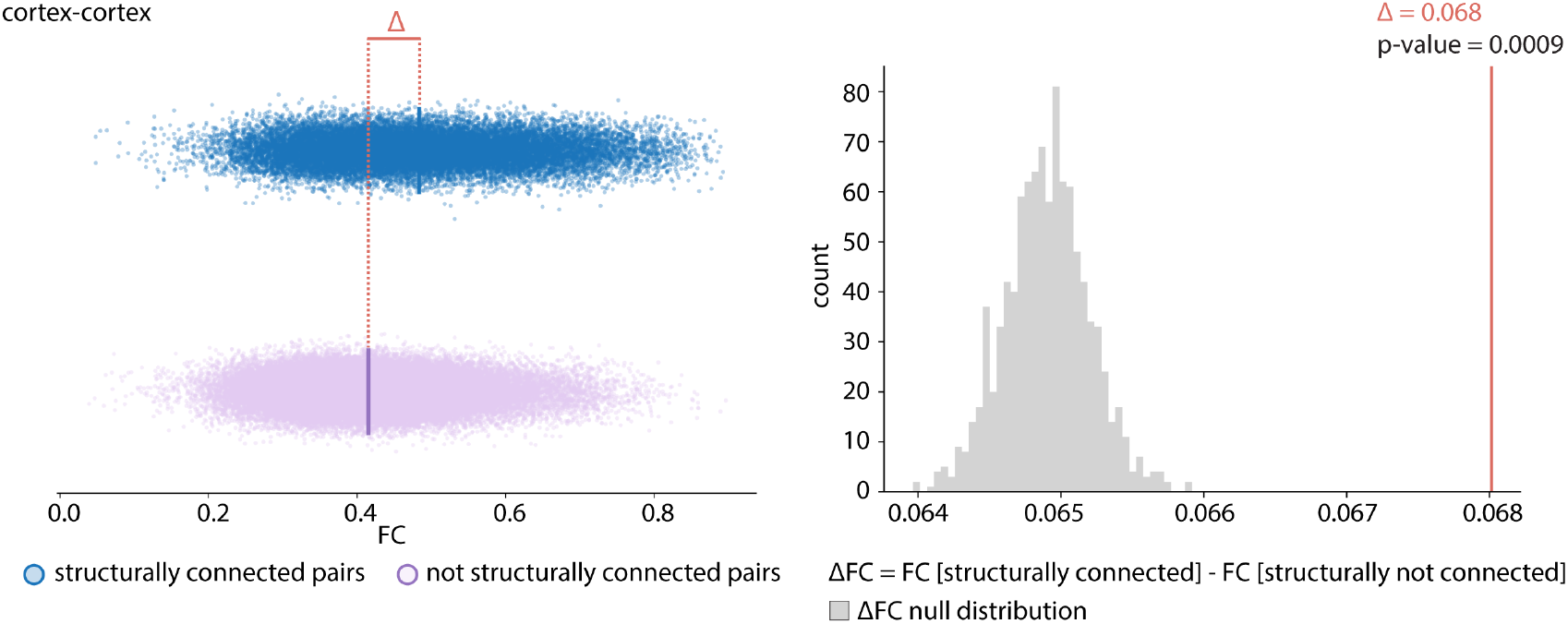
Cortical structural connectivity supports its functional connectivity. Cortex-cortex functional connectivity is stronger for structurally connected region pairs than for not-connected pairs. Statistical significance of the difference in mean functional connectivity between connected and not-connected region pairs is assessed against a null distribution of differences (ΔFC) generated when degree- and edge length-preserving rewired structural networks (*N*_*rewired*_ = 1 000) are used to define anatomical connectivity^75^. The FC difference (ΔFC) corresponded to Cohen’s *d* of 0.63 for the cortex-cortex compartment.

**Figure S15.**
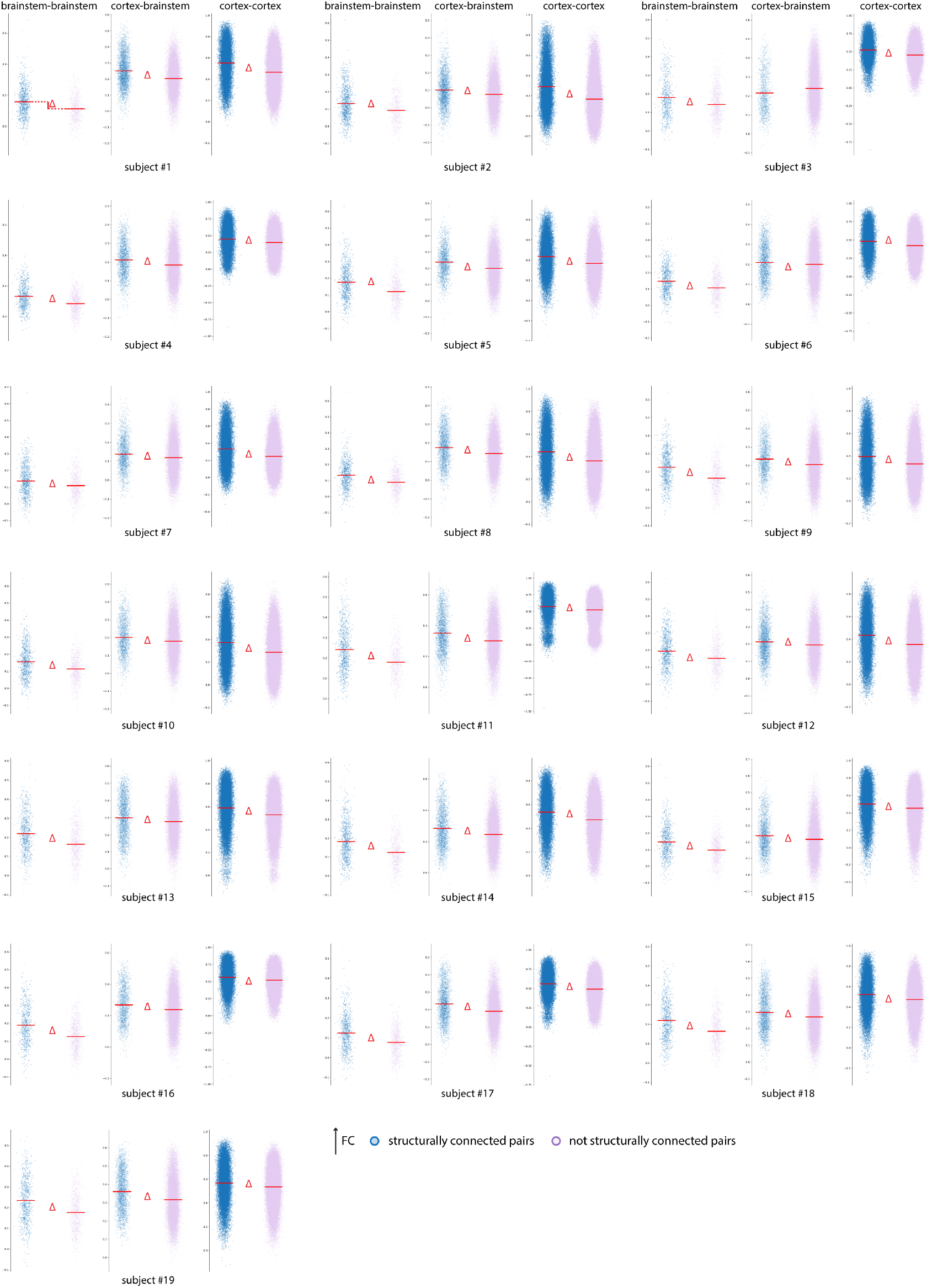
Structural connectivity supports functional connectivity in individuals. We examine whether structural connectivity supports functional connectivity at the individual level by comparing FC values between structurally connected and structurally unconnected region pairs. This analysis was performed separately for three network compartments, including within brainstem, between brainstem and cortex, and within cortex. In all three compartments, structurally connected pairs showed higher FC than structurally unconnected pairs in 18 of 19 individuals. Statistical significance was assessed using a Mann-Whitney U test comparing FC values across connected and unconnected pairs (*p*-value *<* 0.05 is indicated by a red delta (Δ) symbol).

**Figure S16.**
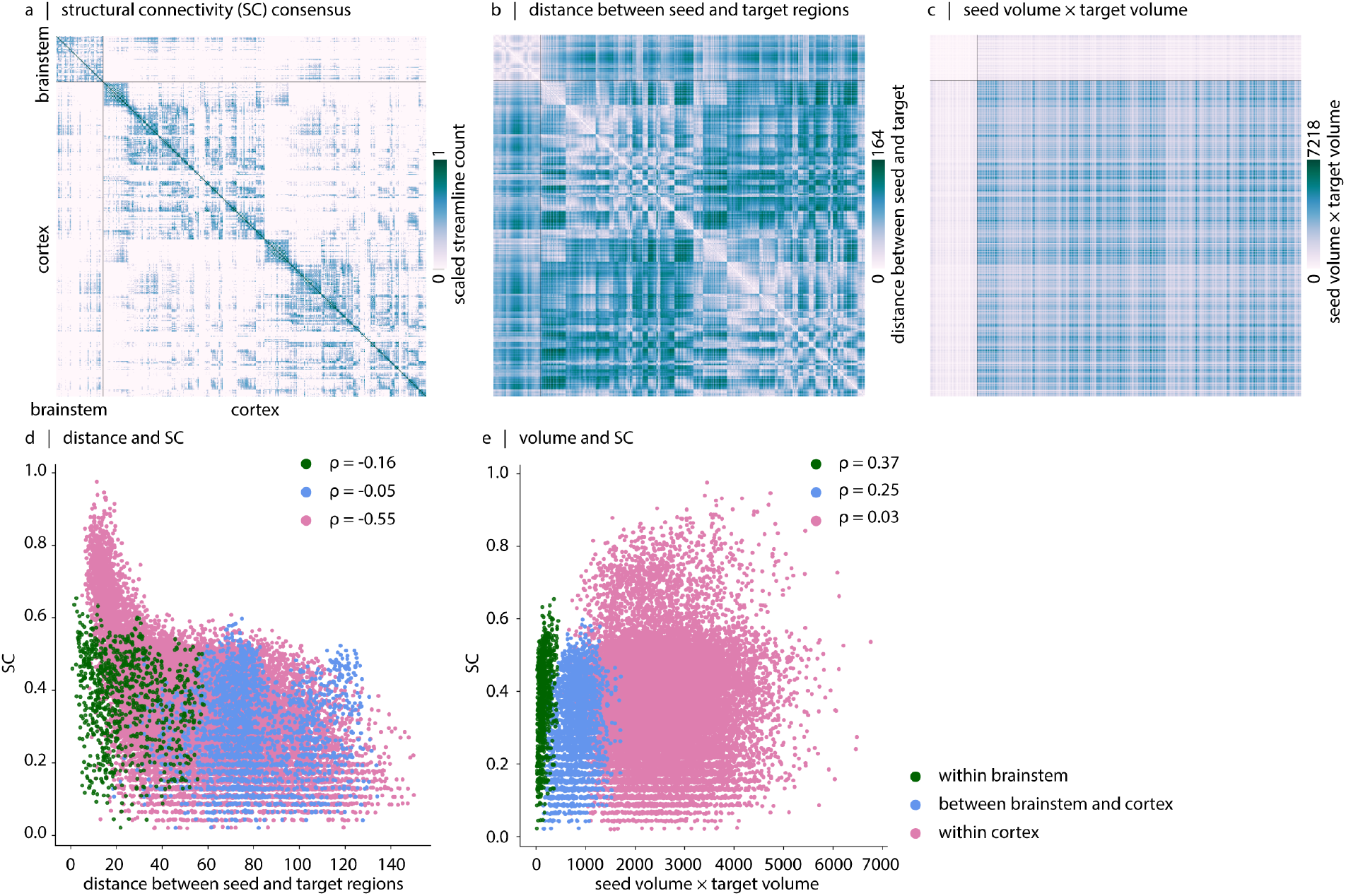
Association between structural connectivity and distance and volume of seed and target regions. (a) Group-consensus structural connectivity (SC) matrix. (b) Distance matrix. Each edge in this matrix represents the Euclidean distance between regional centroids. (c) Volume matrix. Each edge in this matrix represents the product of voxel counts for each pair of regions. (d) The scatter plot shows SC as a function of Euclidean distance between regional centroids (within-brainstem *ρ* =− 0.16, *p* = 1.94 × 10^−6^; between brainstem and cortex *ρ* = −0.05, *p* = 0.01; and within-cortex *ρ* = −0.55, *p ≈* 0). (e) The scatter plot shows SC as a function of regional volumes (within-brainstem *ρ* = 0.37, *p* = 9.89 × 10^−29^; between brainstem and cortex *ρ* = 0.25, *p* = 2.32 × 10^−36^; and within-cortex *ρ* = 0.03, *p* = 5.53 × 10^−5^). Diagonal elements and edges with zero SC were excluded from correlation analyses.

**Figure S17.**
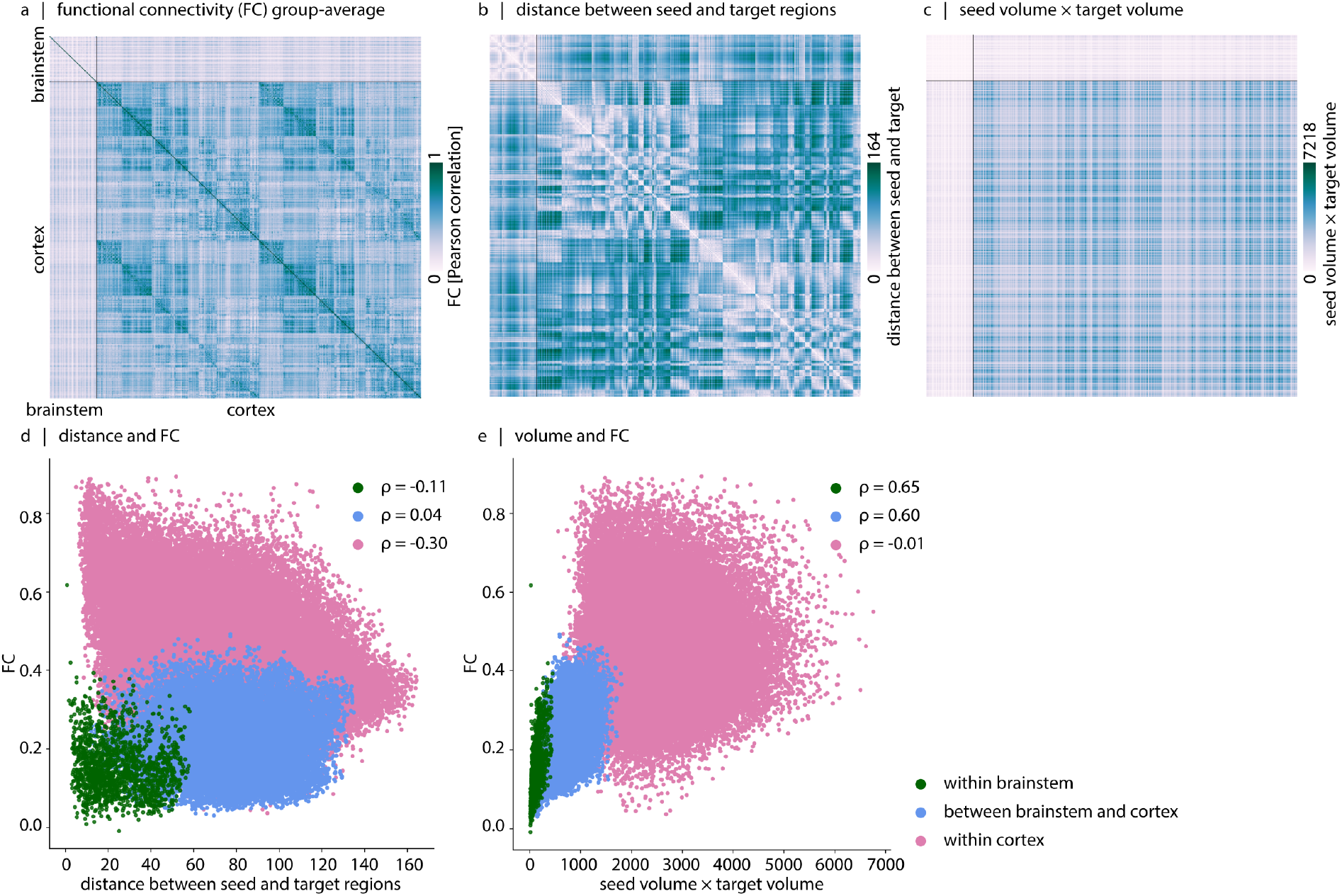
Association between functional connectivity and distance and volume of seed and target regions. (a) Group-averaged functional connectivity (FC) matrix. (b) Distance matrix. Each edge in this matrix represents the Euclidean distance between regional centroids. (c) Volume matrix. Each edge in this matrix represents the product of voxel counts for each pair of regions. Diagonal elements represent the voxel count of individual regions. (d) The scatter plot shows FC as a function of Euclidean distance between regional centroids (within-brainstem *ρ* = −0.11, *p* = 1.56 *×* 10^−5^; between brainstem and cortex *ρ* = 0.04, *p* = 4.59 *×* 10^−8^; and within-cortex *ρ* = −0.30, *p* ≈ 0). (e) The scatter plot shows FC as a function of regional volumes (within-brainstem *ρ* = 0.65, *p* ≈ 0; between brainstem and cortex *ρ* = 0.60, *p* ≈ 0; and within-cortex *ρ* = −0.013, *p* = 2.99*×*10^−4^).

**Figure S18.**
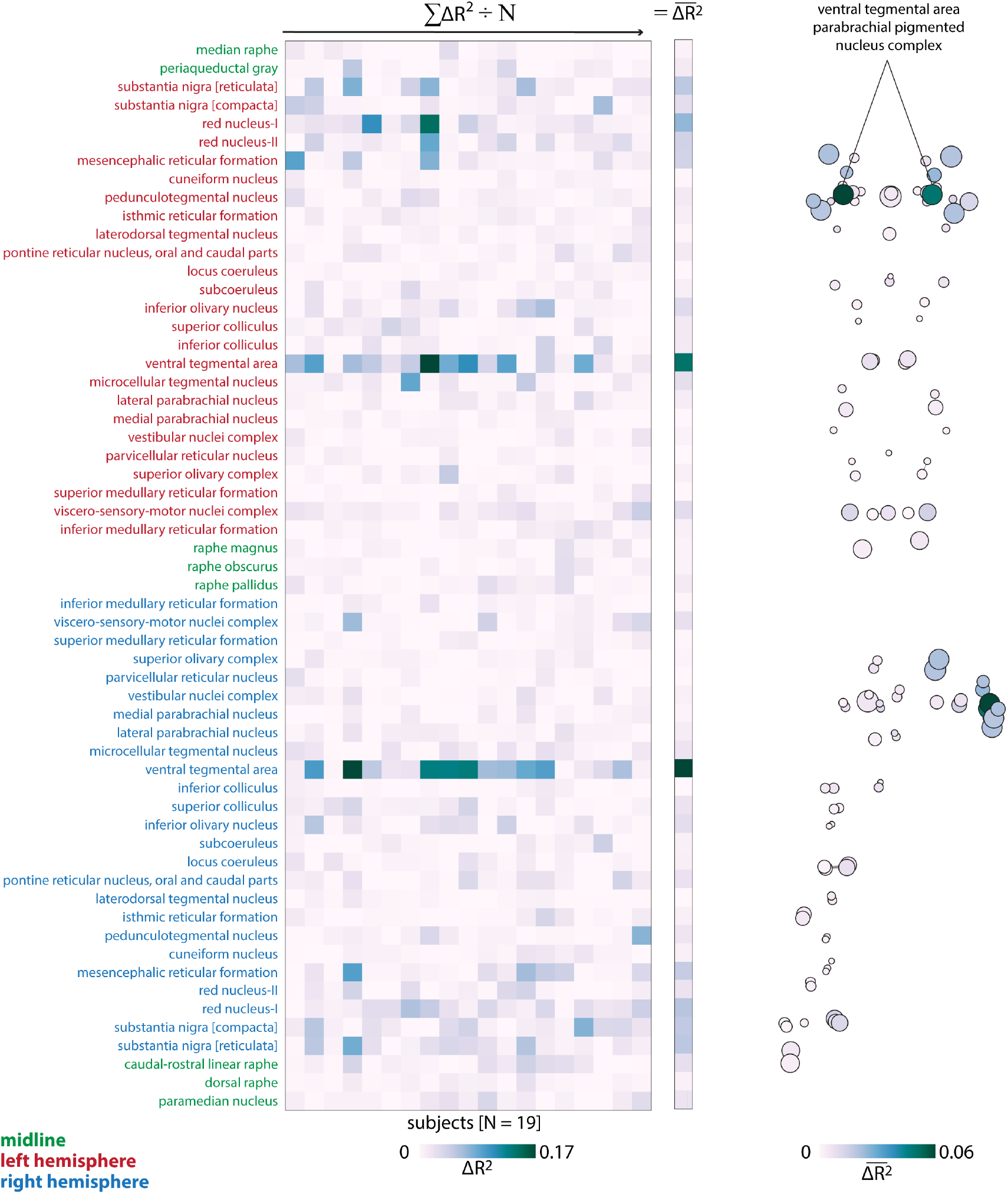
Structural and functional coupling between brainstem and cortex in individual subjects. For each individual, we replicated the analysis explained in the caption of Fig. 4. The pooled result across all 19 individuals is shown on the right column and on the brainstem template.

**Figure S19.**
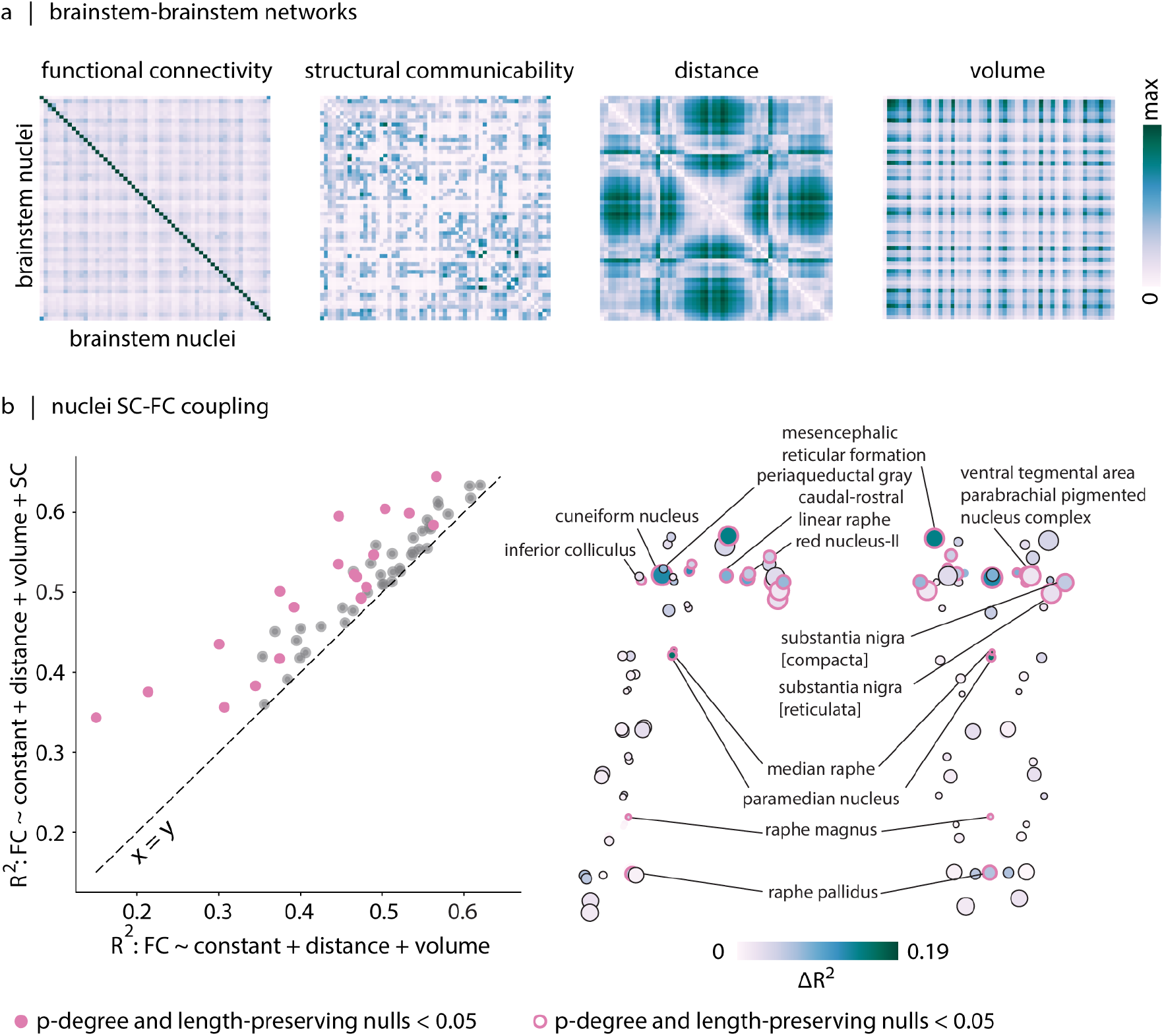
Structural and functional coupling within brainstem. (a) The two leftmost heatmaps show resting-state functional connectivity (FC) and structural communicability between pairs of brainstem nuclei. The third heatmap shows the Euclidean distance between each pair of brainstem nuclei. The fourth heatmap shows the product of the number of voxels for each pair of brainstem nuclei, providing an estimate of their combined volumetric size. (b) We construct two regression models to predict the FC profile of each brainstem nucleus with other brainstem nuclei. The baseline model includes distance and volume regressors and the extended model includes distance, volume and structural communicability profile of the nucleus as regressors. Left: In the scatter plot, baseline model *R*^2^ values are shown on the *x*−axis and extended model *R*^2^ values are shown on the *y*−axis. Each dot in the scatter plot corresponds to a brainstem nucleus. We assess the significance of improvement in model performance (Δ*R*^2^) by comparing the empirical Δ*R*^2^ value to a null distribution of Δ*R*^2^ values obtained from models in which structural communicability is replaced by a communicability matrix that is obtained from degree- and length-preserving null structural networks. Brainstem nuclei showing a significant increase in *R*^2^ when empirical structural communicability is included in the model are shown in pink. Right: The Δ*R*^2^ values for brainstem nuclei are mapped onto the sagittal and coronal views of the brainstem.

**Figure S20.**
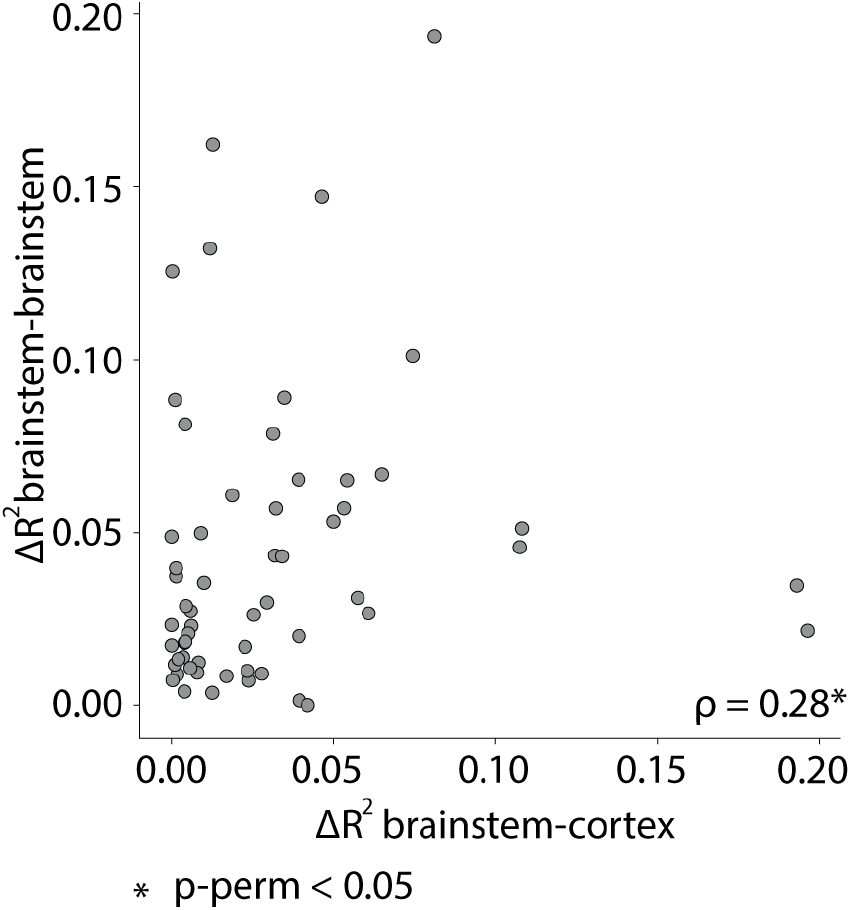
Correspondence of nuclei that show the largest improvement in FC prediction when SC is included in the model. The scatter plot shows the Δ*R*^2^ values for brainstem-cortex (*x*-axis) and brainstem-brainstem connectivity (*y*-axis).

**Figure S21.**
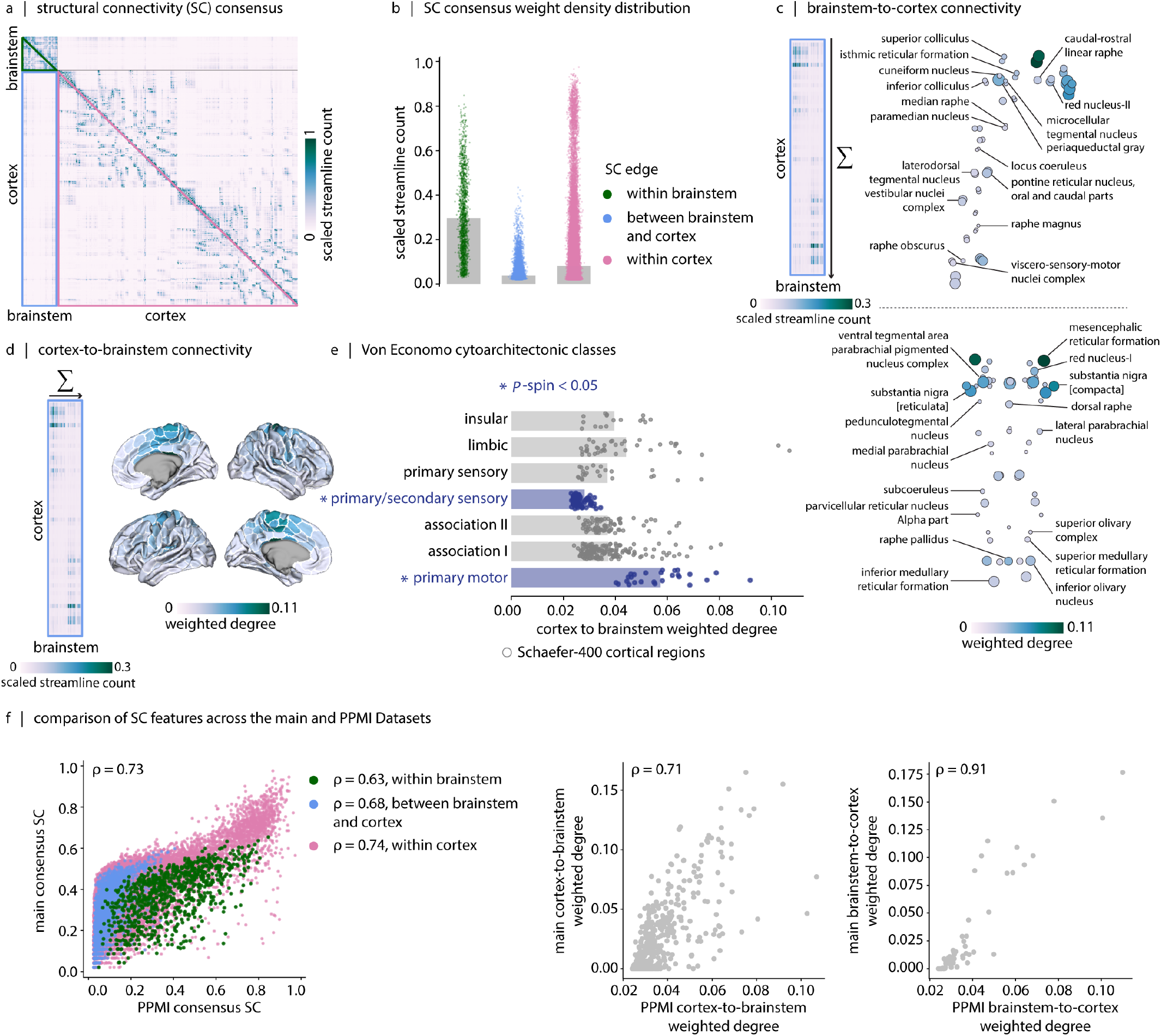
Replication of findings using the PPMI dataset: Structural connectivity of brainstem and cortex. (a) Group-consensus weighted structural connectivity (SC) matrix reconstructed from the 3T PPMI dataset (58 brainstem nuclei and 400 cortical regions, 458 regions × 458 regions). The green triangle highlights the brainstem to brainstem connections, the blue rectangle highlights the connections between cortex and brainstem, and the pink triangle highlights the cortex to cortex connections. (b) Density distributions of non-zero structural connectivity edges within brainstem (green dots), between brainstem and cortex (blue dots), and within cortex (pink dots). Structural connections within brainstem are stronger than structural connections between brainstem and cortex. Structural connections within cortex are also stronger than structural connections between brainstem and cortex (in both cases: two-tailed, *p*_*perm*_ = 9.99 × 10^−4^, *N*_*perm*_ = 1 000). (c) Brainstem-to-cortex weighted degree. Sagittal and coronal views of brainstem nuclei are shown. Node sizes are proportional to the number of voxels per brainstem nucleus, and node colors reflect weighted degree. (d) Cortex-to-brainstem weighted degree. Brain cortical maps are displayed on fs-LR midthickness surfaces. (e) Cortex-to-brainstem weighted degree binned according to the seven cytoarchitectonic classes defined by von Economo^50–52^. Each dot represents a cortical region defined according to the Schaefer-400 parcellation^53^. The brain-stem shows strong structural connectivity with the primary motor cortex and the primary/secondary sensory cortex (two-tailed, *p*_*spin*_ = 2.99 × 10^−3^, 0.029, respectively, *N*_spin_ = 1 000). (f) Cross-dataset comparison of structural connectivity profiles. The left panel compares SC edge weights between the PPMI consensus connectome and the main consensus connectome. The middle panel shows the similarity of cortex-to-brainstem weighted-degree profiles between datasets, with the PPMI dataset on the *x*-axis and the main dataset on the *y*-axis. The right panel shows the corresponding similarity of brainstem-to-cortex weighted-degree profiles across datasets.

**Figure S22.**
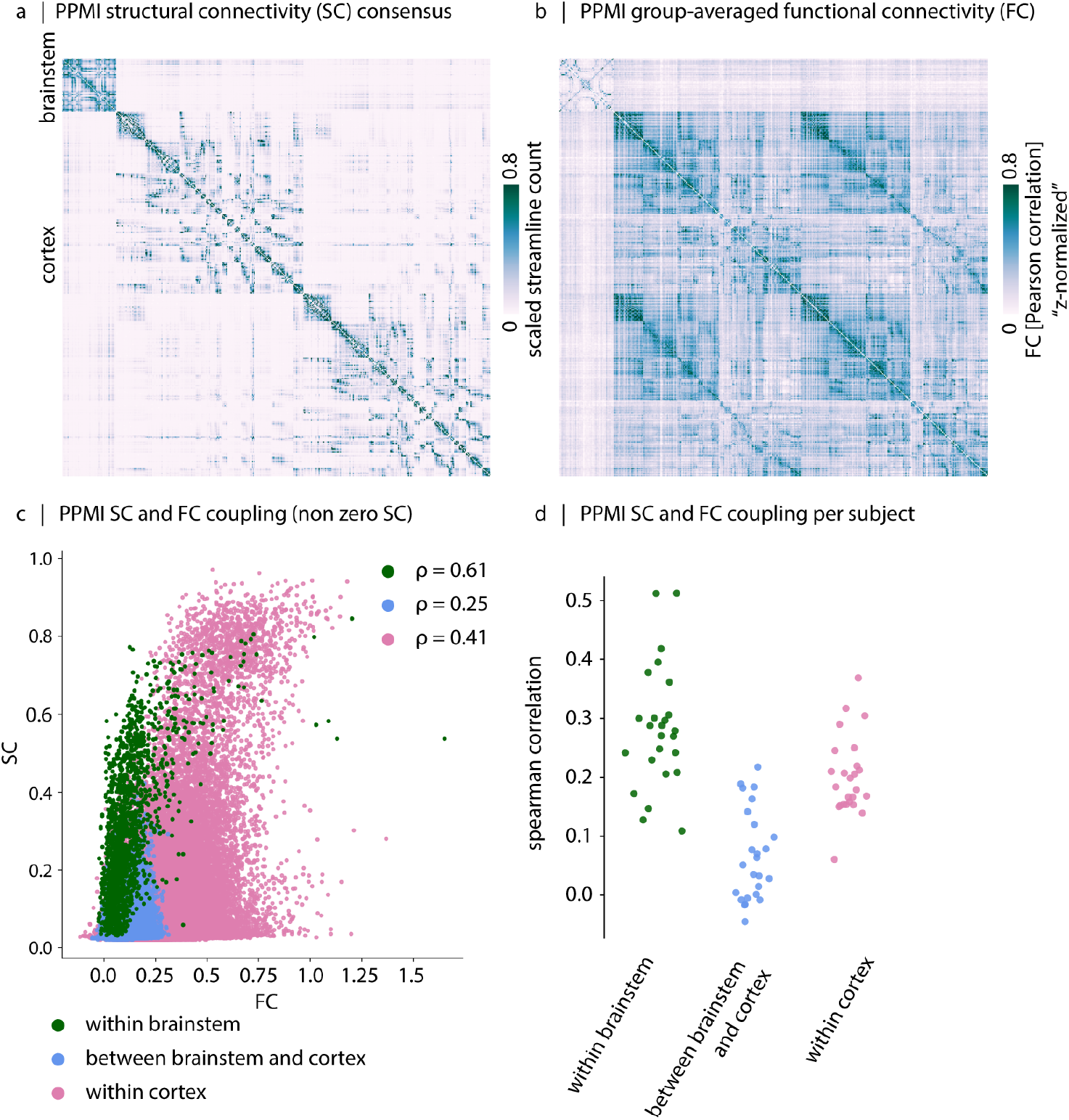
Replication of findings using the PPMI dataset: Association between structural and functional connectivity. (a) Group-consensus weighted structural connectivity (SC) matrix reconstructed from the 3T PPMI dataset. (b) Group-averaged functional connectivity (FC) matrix reconstructed from the 3T PPMI dataset. (c) The scatter plot shows the edge-wise relationship between SC and FC (*z*-transformed). Each dot represents a single edge. Spearman correlation coefficients are reported for each compartment. Diagonal elements and edges with zero SC were excluded from correlation analyses. (d) Distribution of subject-level Spearman correlation coefficients between SC and FC, computed separately within each network compartment.

**Figure S23.**
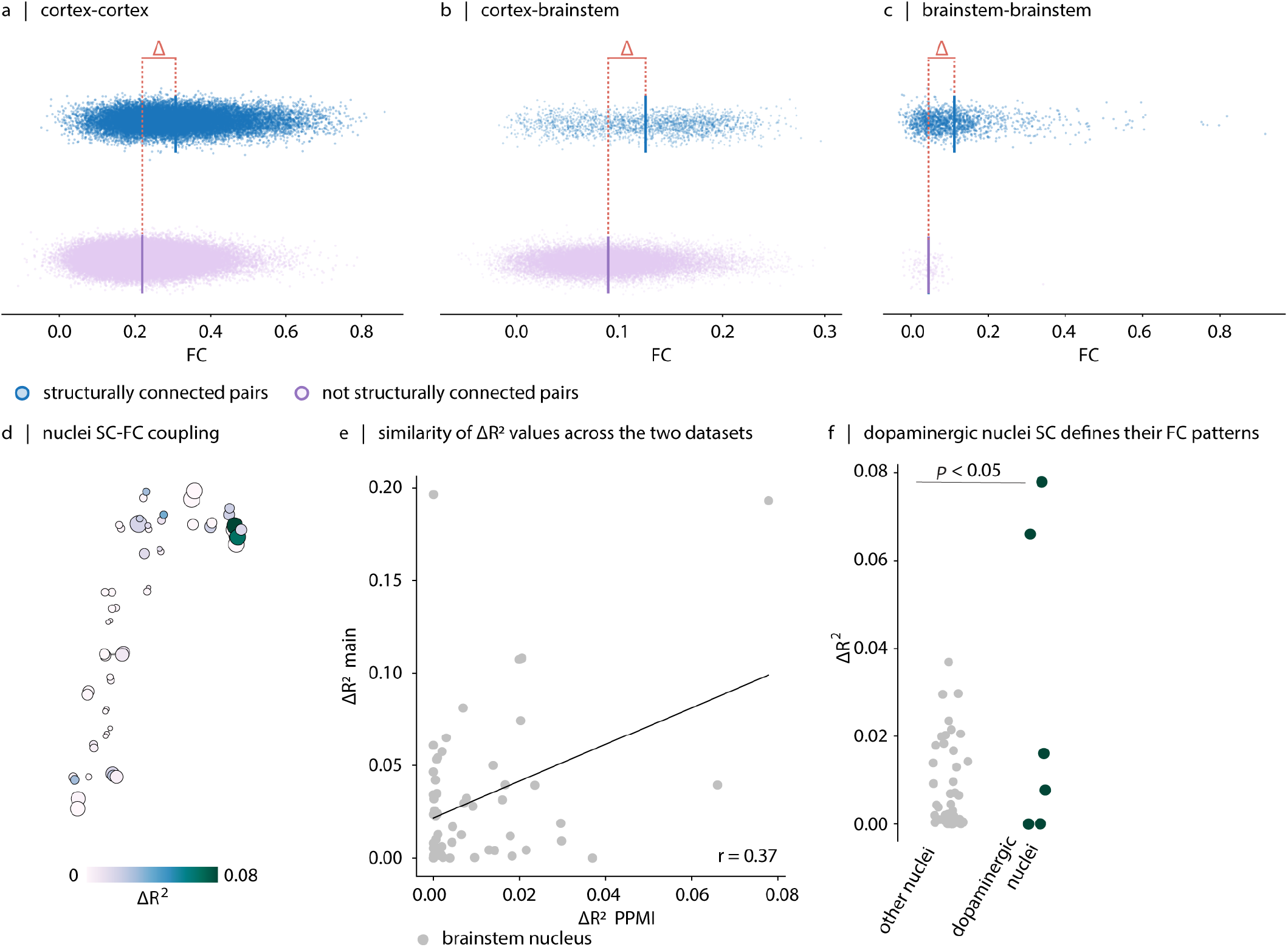
Replication of findings using the PPMI dataset: Brainstem structural connectivity supports its functional connectivity. Functional connectivity is stronger for structurally connected region pairs than for not-connected pairs for (a) cortex-cortex (b) brainstem-cortex, and (c) brainstem-brainstem connections. To facilitate comparison across datasets, the PPMI structural consensus connectome was matched in density to the main consensus connectome before analysis. Statistical significance of the difference in mean functional connectivity between connected and not-connected region pairs is assessed against a null distribution of differences (ΔFC) generated when degree- and edge length-preserving rewired structural networks (*N*_*rewired*_ = 1 000) are used to define anatomical connectivity^75^. The observed ΔFC is 0.09 for cortex-cortex, 0.04 for brainstem-cortex and 0.07 for brainstem-brainstem. *p*_*rewired*_ is equal to 9.99 × 10^−4^ for all three. The FC difference (ΔFC) corresponded to Cohen’s *d* of 0.86 for the cortex-cortex compartment, 0.76 for the cortex-brainstem compartment and 0.65 for the brainstem-brainstem compartment. (d) We construct two regression models to predict the FC profile of each brainstem nucleus with cortical regions. The baseline model includes distance and volume regressors and the extended model includes distance, volume and structural communicability profile of the nucleus as regressors. The resulting increases in explained variance attributable to structural communicability (Δ*R*^2^) for brainstem nuclei are mapped onto the brainstem. (e) Cross-dataset correspondence of brainstem Δ*R*^2^ values. Each point represents a brainstem nucleus, with Δ*R*^2^ from the PPMI dataset shown on the *x*-axis and Δ*R*^2^ from the main dataset shown on the *y*-axis. The spatial correspondence between datasets is *r* = 0.37. (f) Dopaminergic nuclei exhibit larger Δ*R*^2^ values, indicating stronger structure-function coupling beyond geometric constraints (inter-regional distance and volume of target and seed regions) (*p*_*perm*_ = 1.99 *×* 10^−3^, *N*_*perm*_ = 1 000).

**Figure S24.**
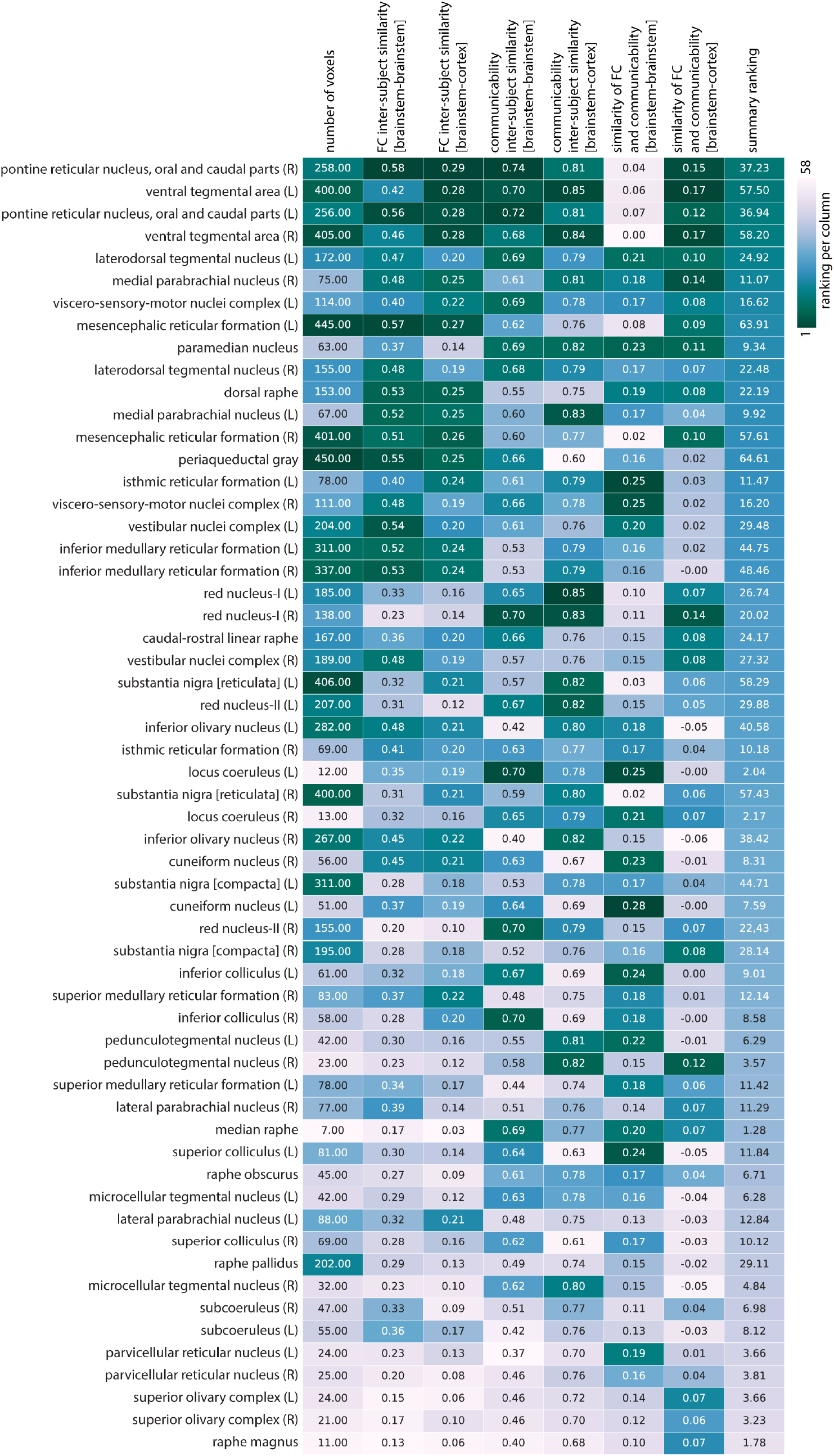
Brainstem nuclei ranking. Brainstem nuclei are ranked based on a composite score integrating nucleus volume, inter-subject similarity of SC/FC within the brainstem and between the brainstem and cortex, and the SC-FC correspondence within the brainstem and between the brainstem and cortex. Cell values indicate the number of voxels, the average correlation between the SC profile of a given nucleus across individuals (reported separately for the within-brainstem and brainstem–cortex connectivity compartments), and the correlation between the SC and FC profiles of a given nucleus for each connectivity compartment. Colors show the descending rank of each nucleus within a given column. The rightmost column shows the average rank of each nucleus across all seven measures (columns) and serves as a summary index of overall connectivity robustness.

**TABLE S1.**
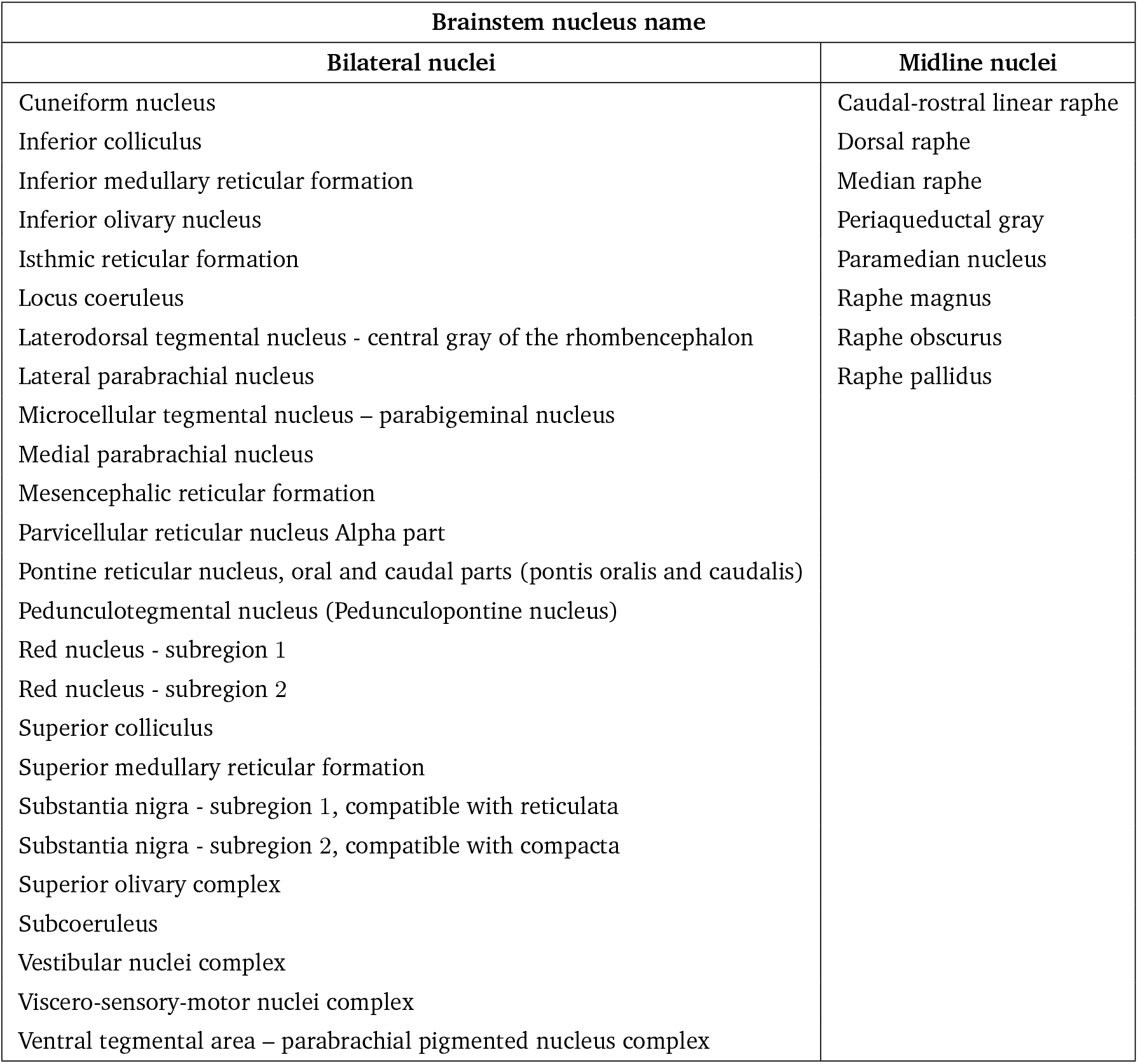
Gray matter regions labeled in the Brainstem Navigator atlas. The Brainstem Navigator atlas includes 58 brainstem nuclei (50 bilateral and eight midline nuclei)^38–44^. Atlas is available at https://www.nitrc.org/projects/brainstemnavig.

**TABLE S2.** Neurosynth terms. List of the 123 Neurosynth terms used in this study^55,56^.

| Neurosynth terms |  |  |  |
| --- | --- | --- | --- |
| action | adaptation | addiction | anticipation |
| anxiety | arousal | association | attention |
| autobiographical_memory | balance | belief | categorization |
| cognitive_control | communication | competition | concept |
| consciousness | consolidation | context | coordination |
| decision | decision_making | detection | discrimination |
| distraction | eating | efficiency | effort |
| emotion | emotion_regulation | empathy | encoding |
| episodic_memory | expectancy | expertise | extinction |
| face_recognition | facial_expression | familiarity | fear |
| fixation | focus | gaze | goal |
| hyperactivity | imagery | impulsivity | induction |
| inference | inhibition | insight | integration |
| intelligence | intention | interference | judgment |
| knowledge | language | language_comprehension | learning |
| listening | localization | loss | maintenance |
| manipulation | meaning | memory | memory_retrieval |
| mental_imagery | monitoring | mood | morphology |
| motor_control | movement | multisensory | naming |
| navigation | object_recognition | pain | perception |
| planning | priming | psychosis | reading |
| reasoning | recall | recognition | rehearsal |
| reinforcement_learning | response_inhibition | response_selection | retention |
| retrieval | reward_anticipation | rhythm | risk |
| rule | salience | search | selective_attention |
| semantic_memory | sentence_comprehension | skill | sleep |
| social_cognition | spatial_attention | speech_perception | speech_production |
| strategy | strength | stress | sustained_attention |
| task_difficulty | thought | uncertainty | updating |
| utility | valence | verbal_fluency | visual_attention |
| visual_perception | word_recognition | working_memory |  |

**TABLE S3.**
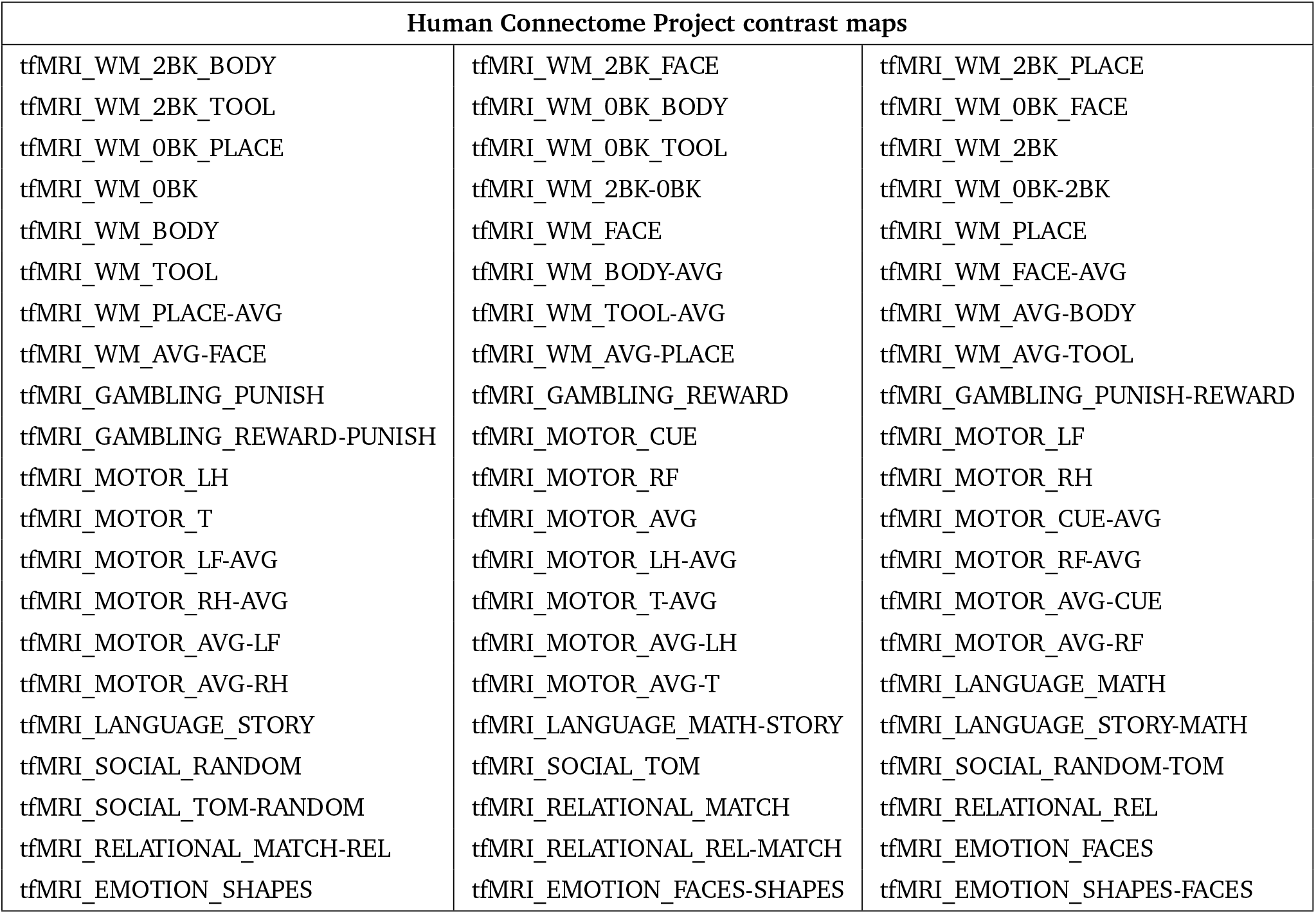
Human Connectome Project contrast maps. List of the 63 HCP contrast maps used in this study^274^.

| Human Connectome Project contrast maps |  |  |
| --- | --- | --- |
| tfMRI_WM_2BK_BODY | tfMRI_WM_2BK_FACE | tfMRI_WM_2BK_PLACE |
| tfMRI_WM_2BK_TOOL | tfMRI_WM_0BK_BODY | tfMRI_WM_0BK_FACE |
| tfMRI_WM_0BK_PLACE | tfMRI_WM_0BK_TOOL | tfMRI_WM_2BK |
| tfMRI_WM_0BK | tfMRI_WM_2BK-0BK | tfMRI_WM_0BK-2BK |
| tfMRI_WM_BODY | tfMRI_WM_FACE | tfMRI_WM_PLACE |
| tfMRI_WM_TOOL | tfMRI_WM_BODY-AVG | tfMRI_WM_FACE-AVG |
| tfMRI_WM_PLACE-AVG | tfMRI_WM_TOOL-AVG | tfMRI_WM_AVG-BODY |
| tfMRI_WM_AVG-FACE | tfMRI_WM_AVG-PLACE | tfMRI_WM_AVG-TOOL |
| tfMRI_GAMBLING_PUNISH | tfMRI_GAMBLING_REWARD | tfMRI_GAMBLING_PUNISH-REWARD |
| tfMRI_GAMBLING_REWARD-PUNISH | tfMRI_MOTOR_CUE | tfMRI_MOTOR_LF |
| tfMRI_MOTOR_LH | tfMRI_MOTOR_RF | tfMRI_MOTOR_RH |
| tfMRI_MOTOR_T | tfMRI_MOTOR_AVG | tfMRI_MOTOR_CUE-AVG |
| tfMRI_MOTOR_LF-AVG | tfMRI_MOTOR_LH-AVG | tfMRI_MOTOR_RF-AVG |
| tfMRI_MOTOR_RH-AVG | tfMRI_MOTOR_T-AVG | tfMRI_MOTOR_AVG-CUE |
| tfMRI_MOTOR_AVG-LF | tfMRI_MOTOR_AVG-LH | tfMRI_MOTOR_AVG-RF |
| tfMRI_MOTOR_AVG-RH | tfMRI_MOTOR_AVG-T | tfMRI_LANGUAGE_MATH |
| tfMRI_LANGUAGE_STORY | tfMRI_LANGUAGE_MATH-STORY | tfMRI_LANGUAGE_STORY-MATH |
| tfMRI_SOCIAL_RANDOM | tfMRI_SOCIAL_TOM | tfMRI_SOCIAL_RANDOM-TOM |
| tfMRI_SOCIAL_TOM-RANDOM | tfMRI_RELATIONAL_MATCH | tfMRI_RELATIONAL_REL |
| tfMRI_RELATIONAL_MATCH-REL | tfMRI_RELATIONAL_REL-MATCH | tfMRI_EMOTION_FACES |
| tfMRI_EMOTION_SHAPES | tfMRI_EMOTION_FACES-SHAPES | tfMRI_EMOTION_SHAPES-FACES |

**TABLE S4.**
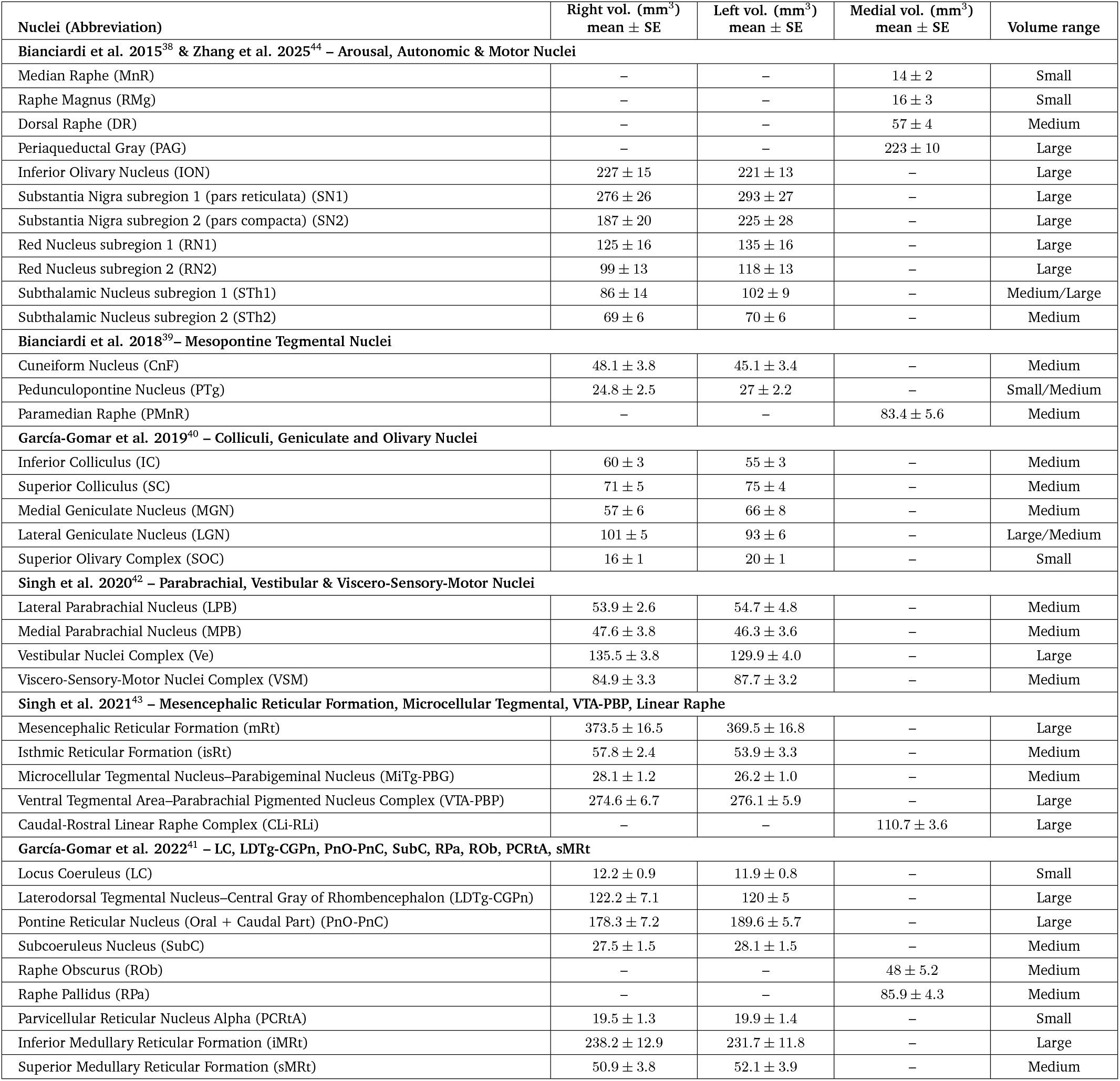
Volumetric Profile of Brainstem Navigator Nuclei at 7T DWI Resolution. Small (*<* 25 mm^3^): small relative to the 1.7 mm isotropic DWI voxels (voxel volume = 4.91 mm^3^)–partial volume effects likely. Medium (25–100 mm^3^): tractography feasible at this resolution. Large (>100 mm^3^): well-sampled by DWI at 1.7 mm isotropic resolution. Atlas is defined at 1 mm^3^ in MNI space from 1.1 mm isotropic 7T MRI (*N* = 12 subjects). In the current study, DWI tractography was performed at 1.7 mm isotropic resolution. Volumes reported as mean *±* SE in native space of the atlas participants (1.1 mm isotropic).

| Nuclei (Abbreviation) | Right vol. (mm<sup>3</sup>)<br>mean ± SE | Left vol. (mm<sup>3</sup>)<br>mean ± SE | Medial vol. (mm<sup>3</sup>)<br>mean ± SE | Volume range |
| --- | --- | --- | --- | --- |
| **Bianciardi et al. 2015<sup>38</sup> & Zhang et al. 2025<sup>44</sup> – Arousal, Autonomic & Motor Nuclei** |  |  |  |  |
| Median Raphe (MnR) | – | – | 14 ± 2 | Small |
| Raphe Magnus (RMg) | – | – | 16 ± 3 | Small |
| Dorsal Raphe (DR) | – | – | 57 ± 4 | Medium |
| Periaqueductal Gray (PAG) | – | – | 223 ± 10 | Large |
| Inferior Olivary Nucleus (ION) | 227 ± 15 | 221 ± 13 | – | Large |
| Substantia Nigra subregion 1 (pars reticulata) (SN1) | 276 ± 26 | 293 ± 27 | – | Large |
| Substantia Nigra subregion 2 (pars compacta) (SN2) | 187 ± 20 | 225 ± 28 | – | Large |
| Red Nucleus subregion 1 (RN1) | 125 ± 16 | 135 ± 16 | – | Large |
| Red Nucleus subregion 2 (RN2) | 99 ± 13 | 118 ± 13 | – | Large |
| Subthalamic Nucleus subregion 1 (STh1) | 86 ± 14 | 102 ± 9 | – | Medium/Large |
| Subthalamic Nucleus subregion 2 (STh2) | 69 ± 6 | 70 ± 6 | – | Medium |
| **Bianciardi et al. 2018<sup>39</sup> – Mesopontine Tegmental Nuclei** |  |  |  |  |
| Cuneiform Nucleus (CnF) | 48.1 ± 3.8 | 45.1 ± 3.4 | – | Medium |
| Pedunculopontine Nucleus (PTg) | 24.8 ± 2.5 | 27 ± 2.2 | – | Small/Medium |
| Paramedian Raphe (PMnR) | – | – | 83.4 ± 5.6 | Medium |
| **García-Gomar et al. 2019<sup>40</sup> – Colliculi, Geniculate and Olivary Nuclei** |  |  |  |  |
| Inferior Colliculus (IC) | 60 ± 3 | 55 ± 3 | – | Medium |
| Superior Colliculus (SC) | 71 ± 5 | 75 ± 4 | – | Medium |
| Medial Geniculate Nucleus (MGN) | 57 ± 6 | 66 ± 8 | – | Medium |
| Lateral Geniculate Nucleus (LGN) | 101 ± 5 | 93 ± 6 | – | Large/Medium |
| Superior Olivary Complex (SOC) | 16 ± 1 | 20 ± 1 | – | Small |
| **Singh et al. 2020<sup>42</sup> – Parabrachial, Vestibular & Viscero-Sensory-Motor Nuclei** |  |  |  |  |
| Lateral Parabrachial Nucleus (LPB) | 53.9 ± 2.6 | 54.7 ± 4.8 | – | Medium |
| Medial Parabrachial Nucleus (MPB) | 47.6 ± 3.8 | 46.3 ± 3.6 | – | Medium |
| Vestibular Nuclei Complex (Ve) | 135.5 ± 3.8 | 129.9 ± 4.0 | – | Large |
| Viscero-Sensory-Motor Nuclei Complex (VSM) | 84.9 ± 3.3 | 87.7 ± 3.2 | – | Medium |
| **Singh et al. 2021<sup>43</sup> – Mesencephalic Reticular Formation, Microcellular Tegmental, VTA-PBP, Linear Raphe** |  |  |  |  |
| Mesencephalic Reticular Formation (mRt) | 373.5 ± 16.5 | 369.5 ± 16.8 | – | Large |
| Isthmic Reticular Formation (isRt) | 57.8 ± 2.4 | 53.9 ± 3.3 | – | Medium |
| Microcellular Tegmental Nucleus–Parabigeminal Nucleus (MiTg-PBG) | 28.1 ± 1.2 | 26.2 ± 1.0 | – | Medium |
| Ventral Tegmental Area–Parabrachial Pigmented Nucleus Complex (VTA-PBP) | 274.6 ± 6.7 | 276.1 ± 5.9 | – | Large |
| Caudal-Rostral Linear Raphe Complex (CLi-RLi) | – | – | 110.7 ± 3.6 | Large |
| **García-Gomar et al. 2022<sup>41</sup> – LC, LDTg-CGPn, PnO-PnC, SubC, RPa, ROb, PCRTA, sMRt** |  |  |  |  |
| Locus Coeruleus (LC) | 12.2 ± 0.9 | 11.9 ± 0.8 | – | Small |
| Laterodorsal Tegmental Nucleus–Central Gray of Rhombencephalon (LDTg-CGPn) | 122.2 ± 7.1 | 120 ± 5 | – | Large |
| Pontine Reticular Nucleus (Oral + Caudal Part) (PnO-PnC) | 178.3 ± 7.2 | 189.6 ± 5.7 | – | Large |
| Subcoeruleus Nucleus (SubC) | 27.5 ± 1.5 | 28.1 ± 1.5 | – | Medium |
| Raphe Obscurus (ROb) | – | – | 48 ± 5.2 | Medium |
| Raphe Pallidus (RPa) | – | – | 85.9 ± 4.3 | Medium |
| Parvicellular Reticular Nucleus Alpha (PCRTA) | 19.5 ± 1.3 | 19.9 ± 1.4 | – | Small |
| Inferior Medullary Reticular Formation (iMRt) | 238.2 ± 12.9 | 231.7 ± 11.8 | – | Large |
| Superior Medullary Reticular Formation (sMRt) | 50.9 ± 3.8 | 52.1 ± 3.9 | – | Medium |

## References

[1] Jessica S Damoiseaux and Michael D Greicius. Greater than the sum of its parts: a review of studies combining structural connectivity and resting-state functional connectivity. Brain Structure and Function, 213(6):525–533, 2009.

[2] Patric Hagmann, Leila Cammoun, Xavier Gigandet, Reto Meuli, Christopher J Honey, Van J Wedeen, and Olaf Sporns. Mapping the structural core of human cerebral cortex. PLoS biology, 6(7):e159, 2008.

[3] Christopher J Honey, Olaf Sporns, Leila Cammoun, Xavier Gigandet, Jean-Philippe Thiran, Reto Meuli, and Patric Hagmann. Predicting human resting-state functional connectivity from structural connectivity. Proceedings of the National Academy of Sciences, 106(6):2035–2040, 2009.

[4] Olaf Sporns. Structure and function of complex brain networks. Dialogues in Clinical Neuroscience, 15(3):247–262, 2013.

[5] Hae-Jeong Park and Karl Friston. Structural and functional brain networks: from connections to cognition. Science, 342(6158):1238411, 2013.

[6] Ann M Hermundstad, Danielle S Bassett, Kevin S Brown, Elissa M Aminoff, David Clewett, Scott Freeman, Amy Frithsen, Arianne Johnson, Christine M Tipper, Michael B Miller, et al. Structural foundations of resting-state and task-based functional connectivity in the human brain. Proceedings of the National Academy of Sciences, 110(15):6169–6174, 2013.

[7] Bratislav Mišić, Richard F Betzel, Marcel A De Reus, Martijn P Van Den Heuvel, Marc G Berman, Anthony R McIntosh, and Olaf Sporns. Network-level structure-function relationships in human neocortex. Cerebral Cortex, 26 (7):3285–3296, 2016.

[8] Pawel Skudlarski, Kanchana Jagannathan, Vince D Calhoun, Michelle Hampson, Beata A Skudlarska, and Godfrey Pearlson. Measuring brain connectivity: diffusion tensor imaging validates resting state temporal correlations. Neuroimage, 43(3):554–561, 2008.

[9] Bertha Vázquez-Rodríguez, Laura E Suárez, Ross D Markello, Golia Shafiei, Casey Paquola, Patric Hagmann, Martijn P Van Den Heuvel, Boris C Bernhardt, R Nathan Spreng, and Bratislav Misic. Gradients of structure– function tethering across neocortex. Proceedings of the National Academy of Sciences, 116(42):21219–21227, 2019.

[10] Laura E Suárez, Ross D Markello, Richard F Betzel, and Bratislav Misic. Linking structure and function in macroscale brain networks. Trends in Cognitive Sciences, 24(4):302–315, 2020.

[11] Zhen-Qi Liu, Golia Shafiei, Sylvain Baillet, and Bratislav Misic. Spatially heterogeneous structure-function coupling in haemodynamic and electromagnetic brain networks. Neuroimage, 278:120276, 2023.

[12] Panagiotis Fotiadis, Linden Parkes, Kathryn A Davis, Theodore D Satterthwaite, Russell T Shinohara, and Dani S Bassett. Structure–function coupling in macroscale human brain networks. Nature Reviews Neuroscience, 25(10):688–704, 2024.

[13] Mandeep S Dagli, John E Ingeholm, and James V Haxby. Localization of cardiac-induced signal change in fmri. Neuroimage, 9(4):407–415, 1999.

[14] Dorit Merhof, Grzegorz Soza, Andreas Stadlbauer, Günther Greiner, and Christopher Nimsky. Correction of susceptibility artifacts in diffusion tensor data using nonlinear registration. Medical Image Analysis, 11(6):588–603, 2007.

[15] Ann K Harvey, Kyle TS Pattinson, Jonathan CW Brooks, Stephen D Mayhew, Mark Jenkinson, and Richard G Wise. Brainstem functional magnetic resonance imaging: disentangling signal from physiological noise. Journal of Magnetic Resonance Imaging: An Official Journal of the International Society for Magnetic Resonance in Medicine, 28(6):1337–1344, 2008.

[16] J Alvarez-Linera. Magnetic resonance techniques for the brainstem. In Seminars in Ultrasound, CT and MRI, volume 31, pages 230–245. Elsevier, 2010.

[17] Anastasia A Ford, Luis Colon-Perez, William T Triplett, Joseph M Gullett, Thomas H Mareci, and David B FitzGerald. Imaging white matter in human brainstem. Frontiers in Human Neuroscience, 7:400, 2013.

[18] Jonathan CW Brooks, Olivia K Faull, Kyle TS Pattinson, and Mark Jenkinson. Physiological noise in brainstem fmri. Frontiers in Human Neuroscience, 7:623, 2013.

[19] F Beissner. Functional mri of the brainstem: common problems and their solutions. Clinical Neuroradiology, 25(Suppl 2):251–257, 2015.

[20] Catie Chang, Erika P Raven, and Jeff H Duyn. Brain– heart interactions: challenges and opportunities with functional magnetic resonance imaging at ultra-high field. Philosophical Transactions of the Royal Society A: Mathematical, Physical and Engineering Sciences, 374 (2067):20150188, 2016.

[21] Roberta Sclocco, Florian Beissner, Marta Bianciardi, Jonathan R Polimeni, and Vitaly Napadow. Challenges and opportunities for brainstem neuroimaging with ultrahigh field mri. Neuroimage, 168:412–426, 2018.

[22] Sina Straub, Benjamin R Knowles, Sebastian Flassbeck, Ruth Steiger, Mark E Ladd, and Elke R Gizewski. Mapping the human brainstem: Brain nuclei and fiber tracts at 3 t and 7 t. NMR in Biomedicine, 32(9):e4118, 2019.

[23] M-Marsel Mesulam. Large-scale neurocognitive networks and distributed processing for attention, language, and memory. Annals of Neurology: Official Journal of the American Neurological Association and the Child Neurology Society, 28(5):597–613, 1990.

[24] Stephen L Foote and Jaime A Pineda. Extrathalamic modulation of cortical function. 1993.

[25] Juan Mena-Segovia, Hana M Sims, Peter J Magill, and J Paul Bolam. Cholinergic brainstem neurons modulate cortical gamma activity during slow oscillations. The Journal of Physiology, 586(12):2947–2960, 2008.

[26] Trevor W Robbins and AFT Arnsten. The neuropsychopharmacology of fronto-executive function: monoaminergic modulation. Annual Review of Neuroscience, 32(1):267–287, 2009.

[27] Melissa R Warden, Aslihan Selimbeyoglu, Julie J Mirzabekov, Maisie Lo, Kimberly R Thompson, Sung-Yon Kim, Avishek Adhikari, Kay M Tye, Loren M Frank, and Karl Deisseroth. A prefrontal cortex–brainstem neuronal projection that controls response to behavioural challenge. Nature, 492(7429):428–432, 2012.

[28] A Moses Lee, Jennifer L Hoy, Antonello Bonci, Linda Wilbrecht, Michael P Stryker, and Cristopher M Niell. Identification of a brainstem circuit regulating visual cortical state in parallel with locomotion. Neuron, 83 (2):455–466, 2014.

[29] Ruud L van den Brink, Thomas Pfeffer, and Tobias H Donner. Brainstem modulation of large-scale intrinsic cortical activity correlations. Frontiers in Human Neuroscience, 13:340, 2019.

[30] Peter Novak, Vera Novak, Allahyar Kangarlu, Amir M Abduljalil, Donald W Chakeres, and Pierre-Marie Robitaille. High resolution mri of the brainstem at 8 t. Journal of Computer Assisted Tomography, 25(2):242–246, 2001.

[31] Andreas Deistung, Andreas Schäfer, Ferdinand Schweser, Uta Biedermann, Daniel Güllmar, Robert Trampel, Robert Turner, and Jürgen R Reichenbach. High-resolution mr imaging of the human brainstem in vivo at 7 tesla. Frontiers in Human Neuroscience, 7:710, 2013.

[32] Birgit R Plantinga, Yasin Temel, Alard Roebroeck, Kâmil Uludağ, Dimo Ivanov, Mark L Kuijf, and Bart M ter Haar Romenij. Ultra-high field magnetic resonance imaging of the basal ganglia and related structures. Frontiers in Human Neuroscience, 8:876, 2014.

[33] Antonio Meola, Fang-Cheng Yeh, Wendy Fellows-Mayle, Jared Weed, and Juan C Fernandez-Miranda. Human connectome-based tractographic atlas of the brainstem connections and surgical approaches. Neurosurgery, 79 (3):437–455, 2016.

[34] Yuchun Tang, Wei Sun, Arthur W Toga, John M Ringman, and Yonggang Shi. A probabilistic atlas of human brainstem pathways based on connectome imaging data. Neuroimage, 169:227–239, 2018.

[35] Gary H Glover, Tie-Qiang Li, and David Ress. Imagebased method for retrospective correction of physiological motion effects in fmri: Retroicor. Magnetic Resonance in Medicine: An Official Journal of the International Society for Magnetic Resonance in Medicine, 44(1):162–167, 2000.

[36] Simone Cauzzo, Kavita Singh, Matthew Stauder, María Guadalupe García-Gomar, Nicola Vanello, Claudio Passino, Jeffrey Staab, Iole Indovina, and Marta Bianciardi. Functional connectome of brainstem nuclei involved in autonomic, limbic, pain and sensory processing in living humans from 7 tesla resting state fmri. Neuroimage, 250:118925, 2022.

[37] Kavita Singh, Simone Cauzzo, María Guadalupe García-Gomar, Matthew Stauder, Nicola Vanello, Claudio Passino, and Marta Bianciardi. Functional connectome of arousal and motor brainstem nuclei in living humans by 7 tesla resting-state fmri. Neuroimage, 249:118865, 2022.

[38] Marta Bianciardi, Nicola Toschi, Brian L Edlow, Cornelius Eichner, Kawin Setsompop, Jonathan R Polimeni, Emery N Brown, Hannah C Kinney, Bruce R Rosen, and Lawrence L Wald. Toward an in vivo neuroimaging template of human brainstem nuclei of the ascending arousal, autonomic, and motor systems. Brain Connectivity, 5(10):597–607, 2015.

[39] Marta Bianciardi, Christian Strong, Nicola Toschi, Brian L Edlow, Bruce Fischl, Emery N Brown, Bruce R Rosen, and Lawrence L Wald. A probabilistic template of human mesopontine tegmental nuclei from in vivo 7 t mri. Neuroimage, 170:222–230, 2018.

[40] María G García-Gomar, Christian Strong, Nicola Toschi, Kavita Singh, Bruce R Rosen, Lawrence L Wald, and Marta Bianciardi. In vivo probabilistic structural atlas of the inferior and superior colliculi, medial and lateral geniculate nuclei and superior olivary complex in humans based on 7 tesla mri. Frontiers in Neuroscience, 13: 764, 2019.

[41] María G García-Gomar, Aleksandar Videnovic, Kavita Singh, Matthew Stauder, Laura D Lewis, Lawrence L Wald, Bruce R Rosen, and Marta Bianciardi. Disruption of brainstem structural connectivity in rem sleep behavior disorder using 7 tesla magnetic resonance imaging. Movement Disorders, 37(4):847–853, 2022.

[42] Kavita Singh, Iole Indovina, Jean C Augustinack, Kimberly Nestor, María G García-Gomar, Jeffrey P Staab, and Marta Bianciardi. Probabilistic template of the lateral parabrachial nucleus, medial parabrachial nucleus, vestibular nuclei complex, and medullary viscerosensory-motor nuclei complex in living humans from 7 tesla mri. Frontiers in Neuroscience, 13:1425, 2020.

[43] Kavita Singh, María Guadalupe García-Gomar, and Marta Bianciardi. Probabilistic atlas of the mesencephalic reticular formation, isthmic reticular formation, microcellular tegmental nucleus, ventral tegmental area nucleus complex, and caudal–rostral linear raphe nucleus complex in living humans from 7 tesla magnetic resonance imaging. Brain Connectivity, 11(8):613–623, 2021.

[44] Jiahe Zhang, Danlei Chen, Philip Deming, Tara Srirangarajan, Jordan E Theriault, Philip A Kragel, Ludger Hartley, Kent M Lee, Kieran McVeigh, Tor D Wager, et al. Cortical and subcortical mapping of the human allostatic–interoceptive system using 7 tesla fmri. Nature Neuroscience, 28(11):2380–2391, 2025.

[45] Bram Stieltjes, Walter E Kaufmann, Peter CM Van Zijl, Kim Fredericksen, Godfrey D Pearlson, Meiyappan Solaiyappan, and Susumu Mori. Diffusion tensor imaging and axonal tracking in the human brainstem. Neuroimage, 14(3):723–735, 2001.

[46] Simon Levinson, Michelle Miller, Ahmed Iftekhar, Monica Justo, Daniel Arriola, Wenxin Wei, Saman Hazany, Josue M Avecillas-Chasin, Taylor P Kuhn, Andreas Horn, et al. A structural connectivity atlas of limbic brainstem nuclei. Frontiers in Neuroimaging, 1:1009399, 2023.

[47] Yu Zhang, Andrei A Vakhtin, Jennifer S Jennings, Payam Massaband, Max Wintermark, Patricia L Craig, J Wesson Ashford, J David Clark, and Ansgar J Furst. Diffusion tensor tractography of brainstem fibers and its application in pain. PloS one, 15(2):e0213952, 2020.

[48] Kavita Singh, María Guadalupe García-Gomar, Simone Cauzzo, Jeffrey P Staab, Iole Indovina, and Marta Bianciardi. Structural connectivity of autonomic, pain, limbic, and sensory brainstem nuclei in living humans based on 7 tesla and 3 tesla mri. Human Brain Mapping, 43 (10):3086–3112, 2022.

[49] Justine Y Hansen, Simone Cauzzo, Kavita Singh, María Guadalupe García-Gomar, James M Shine, Marta Bianciardi, and Bratislav Misic. Integrating brainstem and cortical functional architectures. Nature Neuroscience, 27(12):2500–2511, 2024.

[50] Constantin Economo et al. Die cytoarchitektonik der hirnrinde des erwachsenen menschen. Arch NeurPsyche, 1925.

[51] Constantin Freiherr von Economo, Georg N Koskinas, and Lazaros C Triarhou. Atlas of cytoarchitectonics of the adult human cerebral cortex, volume 10. Karger Basel, 2008.

[52] Lianne H Scholtens, Marcel A de Reus, Siemon C de Lange, Ruben Schmidt, and Martijn P van den Heuvel. An mri von economo–koskinas atlas. Neuroimage, 170:249–256, 2018.

[53] Alexander Schaefer, Ru Kong, Evan M Gordon, Timothy O Laumann, Xi-Nian Zuo, Avram J Holmes, Simon B Eickhoff, and BT Thomas Yeo. Local-global parcellation of the human cerebral cortex from intrinsic functional connectivity mri. Cerebral Cortex, 28(9):3095–3114, 2018.

[54] BT Yeo, Fenna M Krienen, Jorge Sepulcre, Mert R Sabuncu, Danial Lashkari, Marisa Hollinshead, Joshua L Roffman, Jordan W Smoller, Lilla Zöllei, Jonathan R Polimeni, et al. The organization of the human cerebral cortex estimated by intrinsic functional connectivity. Journal of Neurophysiology, 106(3):1125–1165, 2011.

[55] Tal Yarkoni, Russell A Poldrack, Thomas E Nichols, David C Van Essen, and Tor D Wager. Large-scale automated synthesis of human functional neuroimaging data. Nature Methods, 8(8):665–670, 2011.

[56] Simon B Eickhoff, Danilo Bzdok, Angela R Laird, Florian Kurth, and Peter T Fox. Activation likelihood estimation meta-analysis revisited. Neuroimage, 59(3):2349–2361, 2012.

[57] Richard F Betzel, Alessandra Griffa, Patric Hagmann, and Bratislav Mišić. Distance-dependent consensus thresholds for generating group-representative structural brain networks. Network Neuroscience, 3(2):475–496, 2019.

[58] Evan M Gordon, Roselyne J Chauvin, Andrew N Van, Aishwarya Rajesh, Ashley Nielsen, Dillan J Newbold, Charles J Lynch, Nicole A Seider, Samuel R Krimmel, Kristen M Scheidter, et al. A somato-cognitive action network alternates with effector regions in motor cortex. Nature, 617(7960):351–359, 2023.

[59] Ernesto Estrada and Naomichi Hatano. Communicability in complex networks. Physical Review E—Statistical, Nonlinear, and Soft Matter Physics, 77(3):036111, 2008.

[60] Jonathan J Crofts and Desmond J Higham. A weighted communicability measure applied to complex brain networks. Journal of the Royal Society Interface, 6(33):411–414, 2009.

[61] Caio Seguin, Olaf Sporns, Andrew Zalesky, Fernando Calamante, et al. Network communication models narrow the gap between the modular organization of structural and functional brain networks. Neuroimage, 257: 119323, 2022.

[62] Joaquín Goñi, Martijn P Van Den Heuvel, Andrea Avena-Koenigsberger, Nieves Velez de Mendizabal, Richard F Betzel, Alessandra Griffa, Patric Hagmann, Bernat Corominas-Murtra, Jean-Philippe Thiran, and Olaf Sporns. Resting-brain functional connectivity predicted by analytic measures of network communication. Proceedings of the National Academy of Sciences, 111(2): 833–838, 2014.

[63] Jennifer Andreotti, Kay Jann, Lester Melie-Garcia, Stéphanie Giezendanner, Eugenio Abela, Roland Wiest, Thomas Dierks, and Andrea Federspiel. Validation of network communicability metrics for the analysis of brain structural networks. PloS One, 9(12):e115503, 2014.

[64] Caio Seguin, Ye Tian, and Andrew Zalesky. Network communication models improve the behavioral and functional predictive utility of the human structural connectome. Network Neuroscience, 4(4):980–1006, 2020.

[65] Robert H Wurtz and Joanne E Albano. Visual-motor function of the primate superior colliculus. Annual Review of Neuroscience, 3(1):189–226, 1980.

[66] James M Sprague and Thomas H Meikle Jr. The role of the superior colliculus in visually guided behavior. Experimental Neurology, 11(1):115–146, 1965.

[67] Christine E Collins, David C Lyon, and Jon H Kaas. Distribution across cortical areas of neurons projecting to the superior colliculus in new world monkeys. The Anatomical Record Part A: Discoveries in Molecular, Cellular, and Evolutionary Biology: An Official Publication of the American Association of Anatomists, 285(1):619–627, 2005.

[68] Christina M Cerkevich, David C Lyon, Pooja Balaram, and Jon H Kaas. Distribution of cortical neurons projecting to the superior colliculus in macaque monkeys. Eye and Brain, 6(sup1):121–137, 2014.

[69] Jean-Pierre Hornung. The human raphe nuclei and the serotonergic system. Journal of Chemical Neuroanatomy, 26(4):331–343, 2003.

[70] S Mark Williams and PS Goldman-Rakic. Widespread origin of the primate mesofrontal dopamine system. Cerebral Cortex, 8(4):321–345, 1998.

[71] David B Carr and Susan R Sesack. Projections from the rat prefrontal cortex to the ventral tegmental area: target specificity in the synaptic associations with mesoaccumbens and mesocortical neurons. Journal of Neuroscience, 20(10):3864–3873, 2000.

[72] Ethan S Bromberg-Martin, Masayuki Matsumoto, and Okihide Hikosaka. Dopamine in motivational control: rewarding, aversive, and alerting. Neuron, 68(5):815–834, 2010.

[73] Marisela Morales and Elyssa B Margolis. Ventral tegmental area: cellular heterogeneity, connectivity and behaviour. Nature Reviews Neuroscience, 18(2):73–85, 2017.

[74] Russell A Poldrack, Aniket Kittur, Donald Kalar, Eric Miller, Christian Seppa, Yolanda Gil, D Stott Parker, Fred W Sabb, and Robert M Bilder. The cognitive atlas: toward a knowledge foundation for cognitive neuroscience. Frontiers in Neuroinformatics, 5:17, 2011.

[75] Richard F Betzel and Danielle S Bassett. Specificity and robustness of long-distance connections in weighted, interareal connectomes. Proceedings of the National Academy of Sciences, 115(21):E4880–E4889, 2018.

[76] Christopher J Honey, Jean-Philippe Thivierge, and Olaf Sporns. Can structure predict function in the human brain? Neuroimage, 52(3):766–776, 2010.

[77] Razia Azen and David V Budescu. The dominance analysis approach for comparing predictors in multiple regression. Psychological Methods, 8(2):129, 2003.

[78] Yusuke Adachi, Takahiro Osada, Olaf Sporns, Takamitsu Watanabe, Teppei Matsui, Kentaro Miyamoto, and Yasushi Miyashita. Functional connectivity between anatomically unconnected areas is shaped by collective network-level effects in the macaque cortex. Cerebral Cortex, 22(7):1586–1592, 2012.

[79] Kelly Shen, R Matthew Hutchison, Gleb Bezgin, Stefan Everling, and Anthony R McIntosh. Network structure shapes spontaneous functional connectivity dynamics. Journal of Neuroscience, 35(14):5579–5588, 2015.

[80] Qiang Gu. Neuromodulatory transmitter systems in the cortex and their role in cortical plasticity. Neuroscience, 111(4):815–835, 2002.

[81] Lisa A Briand, Howard Gritton, William M Howe, Damon A Young, and Martin Sarter. Modulators in concert for cognition: modulator interactions in the prefrontal cortex. Progress in Neurobiology, 83(2):69–91, 2007.

[82] David T George, Rezvan Ameli, and George F Koob. Periaqueductal gray sheds light on dark areas of psychopathology. Trends in Neurosciences, 42(5):349–360, 2019.

[83] Kenneth Marek, Danna Jennings, Shirley Lasch, Andrew Siderowf, Caroline Tanner, Tanya Simuni, Chris Coffey, Karl Kieburtz, Emily Flagg, Sohini Chowdhury, et al. The parkinson progression marker initiative (ppmi). Progress in Neurobiology, 95(4):629–635, 2011.

[84] Uğur Türe, M Gazi Yaşargil, Allan H Friedman, and Ossama Al-Mefty. Fiber dissection technique: lateral aspect of the brain. Neurosurgery, 47(2):417–427, 2000.

[85] Abhidha Shah, Sukhdeep Singh Jhawar, Maximilliano Nunez, Aimee Goel, and Atul Goel. Brainstem anatomy: A study on the basis of the pattern of fiber organization. World Neurosurgery, 134:e826–e846, 2020.

[86] John D French, FK Von Amerongen, and HW Magoun. An activating system in brain stem of monkey. AMA Archives of Neurology & Psychiatry, 68(5):577–590, 1952.

[87] K Hartmann von Monakow, K Akert, and H Künzle. Projections of precentral and premotor cortex to the red nucleus and other midbrain areas in macaca fascicularis. Experimental Brain Research, 34(1):91–105, 1979.

[88] Robert M Beckstead, Joel R Morse, and Ralph Norgren. The nucleus of the solitary tract in the monkey: projections to the thalamus and brain stem nuclei. Journal of Comparative Neurology, 190(2):259–282, 1980.

[89] Linda J Porrino and Patricia S Goldman-Rakic. Brainstem innervation of prefrontal and anterior cingulate cortex in the rhesus monkey revealed by retrograde transport of hrp. Journal of Comparative Neurology, 205 (1):63–76, 1982.

[90] Sten Grillner and Peter Wallen. Central pattern generators for locomotion, with special reference to vertebrates. Annual Review of Neuroscience, 1985.

[91] Christine A Livingston and Robert B Leonard. Locomotion evoked by stimulation of the brain stem in the atlantic stingray, dasyatis sabina. Journal of Neuroscience, 10(1):194–204, 1990.

[92] Schahram Akbarian, Otto-Joachim Grüsser, and Wolfgang O Guldin. Corticofugal connections between the cerebral cortex and brainstem vestibular nuclei in the macaque monkey. Journal of Comparative Neurology, 339(3):421–437, 1994.

[93] AD Craig. Distribution of brainstem projections from spinal lamina i neurons in the cat and the monkey. Journal of Comparative Neurology, 361(2):225–248, 1995.

[94] Patrick J Whelan. Control of locomotion in the decerebrate cat. Progress in Neurobiology, 49(5):481–515, 1996.

[95] Patricia Del Cerro, Ángel Rodríguez-De-Lope, and Jorge E Collazos-Castro. The cortical motor system in the domestic pig: origin and termination of the corticospinal tract and cortico-brainstem projections. Frontiers in Neuroanatomy, 15:748050, 2021.

[96] Seung Wook Oh, Julie A Harris, Lydia Ng, Brent Winslow, Nicholas Cain, Stefan Mihalas, Quanxin Wang, Chris Lau, Leonard Kuan, Alex M Henry, et al. A mesoscale connectome of the mouse brain. Nature, 508 (7495):207–214, 2014.

[97] B De Coene, JV Hajnal, JM Pennock, and GM Bydder. Mri of the brain stem using fluid attenuated inversion recovery pulse sequences. Neuroradiology, 35(5):327–331, 1993.

[98] María Guadalupe García-Gomar, Kavita Singh, Simone Cauzzo, and Marta Bianciardi. In vivo structural connectome of arousal and motor brainstem nuclei by 7 tesla and 3 tesla mri. Human Brain Mapping, 43(14):4397–4421, 2022.

[99] HGJM Kuypers. A new look at the organization of the motor system. Progress in Brain Research, 57:381–403, 1982.

[100] Akinsegun Akintunde and Donald F Buxton. Origins and collateralization of corticospinal, corticopontine, corticorubral and corticostriatal tracts: a multiple retrograde fluorescent tracing study. Brain Research, 586(2):208–218, 1992.

[101] D Purves and GJ Augustine. The primary motor cortex: upper motor neurons that initiate complex voluntary movements in: D, p., g, ja, fitzpatrick d, et al. Neuroscience. Sinauer Associates, Sunderland., 2001.

[102] RJ Nudo and RB Masterton. Descending pathways to the spinal cord: a comparative study of 22 mammals. Journal of Comparative Neurology, 277(1):53–79, 1988.

[103] Sten Grillner. The motor infrastructure: from ion channels to neuronal networks. Nature Reviews Neuroscience, 4(7):573–586, 2003.

[104] Quentin Welniarz, Isabelle Dusart, and Emmanuel Roze. The corticospinal tract: Evolution, development, and human disorders. Developmental Neurobiology, 77(7): 810–829, 2017.

[105] Robert D Oades and Glenda M Halliday. Ventral tegmental (a10) system: neurobiology. 1. anatomy and connectivity. Brain Research Reviews, 12(2):117–165, 1987.

[106] Jonas A Hosp, Volker Arnd Coenen, Michel Rijntjes, Karl Egger, Horst Urbach, Cornelius Weiller, and Marco Reisert. Ventral tegmental area connections to motor and sensory cortical fields in humans. Brain Structure and Function, 224(8):2839–2855, 2019.

[107] Muhammad Zubair, Sjoerd R Murris, Kaoru Isa, Hirotaka Onoe, Yoshinori Koshimizu, Kenta Kobayashi, Wim Vanduffel, and Tadashi Isa. Divergent whole brain projections from the ventral midbrain in macaques. Cerebral Cortex, 31(6):2913–2931, 2021.

[108] TM Lock, JS Baizer, and DB Bender. Distribution of corticotectal cells in macaque. Experimental Brain Research, 151(4):455–470, 2003.

[109] Richard J Krauzlis, Lee P Lovejoy, and Alexandre Zénon. Superior colliculus and visual spatial attention. Annual Review of Neuroscience, 36(1):165–182, 2013.

[110] OT Phillipson. Afferent projections to the ventral tegmental area of tsai and interfascicular nucleus: a horseradish peroxidase study in the rat. Journal of Comparative Neurology, 187(1):117–143, 1979.

[111] EJ Van Bockstaele and VM Pickel. Gaba-containing neurons in the ventral tegmental area project to the nucleus accumbens in rat brain. Brain Research, 682(1-2):215–221, 1995.

[112] Tsuyoshi Yamaguchi, Hui-Ling Wang, Xueping Li, Tsz H Ng, and Marisela Morales. Mesocorticolimbic glutamatergic pathway. Journal of Neuroscience, 31(23): 8476–8490, 2011.

[113] Blake L Elliott, Kimberlee D’Ardenne, Vishnu P Murty, Gene A Brewer, and Samuel M McClure. Midbrain– hippocampus structural connectivity selectively predicts motivated memory encoding. Journal of Neuroscience, 42(50):9426–9434, 2022.

[114] Fares JP Sayegh, Lionel Mouledous, Catherine Macri, Juliana Pi Macedo, Camille Lejards, Claire Rampon, Laure Verret, and Lionel Dahan. Ventral tegmental area dopamine projections to the hippocampus trigger longterm potentiation and contextual learning. Nature Communications, 15(1):4100, 2024.

[115] Klaus H Maier-Hein, Peter F Neher, Jean-Christophe Houde, Marc-Alexandre Côté, Eleftherios Garyfallidis, Jidan Zhong, Maxime Chamberland, Fang-Cheng Yeh, Ying-Chia Lin, Qing Ji, et al. The challenge of mapping the human connectome based on diffusion tractography. Nature Communications, 8(1):1349, 2017.

[116] Kurt G Schilling, Alessandro Daducci, Klaus Maier-Hein, Cyril Poupon, Jean-Christophe Houde, Vishwesh Nath, Adam W Anderson, Bennett A Landman, and Maxime Descoteaux. Challenges in diffusion mri tractography– lessons learned from international benchmark competitions. Magnetic Resonance Imaging, 57:194–209, 2019.

[117] Kurt G Schilling, Laurent Petit, Francois Rheault, Samuel Remedios, Carlo Pierpaoli, Adam W Anderson, Bennett A Landman, and Maxime Descoteaux. Brain connections derived from diffusion mri tractography can be highly anatomically accurate—if we know where white matter pathways start, where they end, and where they do not go. Brain Structure and Function, 225(8):2387–2402, 2020.

[118] Ben Jeurissen, Maxime Descoteaux, Susumu Mori, and Alexander Leemans. Diffusion mri fiber tractography of the brain. NMR in Biomedicine, 32(4):e3785, 2019.

[119] Stamatios N Sotiropoulos and Andrew Zalesky. Building connectomes using diffusion mri: why, how and but. NMR in Biomedicine, 32(4):e3752, 2019.

[120] Francois Rheault, Philippe Poulin, Alex Valcourt Caron, Etienne St-Onge, and Maxime Descoteaux. Common misconceptions, hidden biases and modern challenges of dmri tractography. Journal of Neural Engineering, 17 (1):011001, 2020.

[121] Bratislav Mišić, Richard F Betzel, Azadeh Nematzadeh, Joaquin Goni, Alessandra Griffa, Patric Hagmann, Alessandro Flammini, Yong-Yeol Ahn, and Olaf Sporns. Cooperative and competitive spreading dynamics on the human connectome. Neuron, 86(6):1518–1529, 2015.

[122] Bharat B Biswal and Lucina Q Uddin. The history and future of resting-state functional magnetic resonance imaging. Nature, 641(8065):1121–1131, 2025.

[123] Maria Giulia Preti and Dimitri Van De Ville. Decoupling of brain function from structure reveals regional behavioral specialization in humans. Nature Communications, 10(1):4747, 2019.

[124] Graham L Baum, Zaixu Cui, David R Roalf, Rastko Ciric, Richard F Betzel, Bart Larsen, Matthew Cieslak, Philip A Cook, Cedric H Xia, Tyler M Moore, et al. Development of structure–function coupling in human brain networks during youth. Proceedings of the National Academy of Sciences, 117(1):771–778, 2020.

[125] Vincent Bazinet, Reinder Vos de Wael, Patric Hagmann, Boris C Bernhardt, and Bratislav Misic. Multiscale communication in cortico-cortical networks. Neuroimage, 243:118546, 2021.

[126] Zijin Gu, Keith Wakefield Jamison, Mert Rory Sabuncu, and Amy Kuceyeski. Heritability and interindividual variability of regional structure-function coupling. Nature Communications, 12(1):4894, 2021.

[127] Zhen-Qi Liu, Bertha Vazquez-Rodriguez, R Nathan Spreng, Boris C Bernhardt, Richard F Betzel, and Bratislav Misic. Time-resolved structure-function coupling in brain networks. Communications Biology, 5(1):532, 2022.

[128] Farnaz Zamani Esfahlani, Joshua Faskowitz, Jonah Slack, Bratislav Mišić, and Richard F Betzel. Local structure-function relationships in human brain networks across the lifespan. Nature Communications, 13 (1):2053, 2022.

[129] Justine Y Hansen, Golia Shafiei, Katharina Voigt, Emma X Liang, Sylvia ML Cox, Marco Leyton, Sharna D Jamadar, and Bratislav Misic. Integrating multimodal and multiscale connectivity blueprints of the human cerebral cortex in health and disease. Plos biology, 21 (9):e3002314, 2023.

[130] Zhen-Qi Liu, Andrea I Luppi, Justine Y Hansen, Ye Ella Tian, Andrew Zalesky, BT Thomas Yeo, Ben D Fulcher, and Bratislav Misic. Benchmarking methods for mapping functional connectivity in the brain. Nature Methods, pages 1–10, 2025.

[131] Mark C Nelson, Wen Da Lu, Ilana R Leppert, Heather A Hansen, Christopher D Rowley, Bratislav Misic, and Christine L Tardif. The role of white matter myelin in structural-functional network coupling. bioRxiv, pages 2025–07, 2025.

[132] Martin A Koch, David G Norris, and Margret Hund-Georgiadis. An investigation of functional and anatomical connectivity using magnetic resonance imaging. Neuroimage, 16(1):241–250, 2002.

[133] Panagiotis Fotiadis, Amy FT Arnsten, Linden Parkes, Theodore D Satterthwaite, Russell T Shinohara, and Dani S Bassett. Biological substrates of structurefunction coupling in brain networks. Neuroscience & Biobehavioral Reviews, page 106581, 2026.

[134] Ed Bullmore and Olaf Sporns. The economy of brain network organization. Nature Reviews Neuroscience, 13 (5):336–349, 2012.

[135] Panagiotis Fotiadis, Matthew Cieslak, Xiaosong He, Lorenzo Caciagli, Mathieu Ouellet, Theodore D Satterthwaite, Russell T Shinohara, and Dani S Bassett. Myelination and excitation-inhibition balance synergistically shape structure-function coupling across the human cortex. Nature Communications, 14(1):6115, 2023.

[136] Gustavo Deco, Adrián Ponce-Alvarez, Dante Mantini, Gian Luca Romani, Patric Hagmann, and Maurizio Corbetta. Resting-state functional connectivity emerges from structurally and dynamically shaped slow linear fluctuations. Journal of Neuroscience, 33(27):11239–11252, 2013.

[137] Gustavo Deco, Josephine Cruzat, Joana Cabral, Gitte M Knudsen, Robin L Carhart-Harris, Peter C Whybrow, Nikos K Logothetis, and Morten L Kringelbach. Wholebrain multimodal neuroimaging model using serotonin receptor maps explains non-linear functional effects of lsd. Current Biology, 28(19):3065–3074, 2018.

[138] Peng Wang, Ru Kong, Xiaolu Kong, Raphaël Liégeois, Csaba Orban, Gustavo Deco, Martijn P Van Den Heuvel, and BT Thomas Yeo. Inversion of a large-scale circuit model reveals a cortical hierarchy in the dynamic resting human brain. Science Advances, 5(1):eaat7854, 2019.

[139] Laura E Suárez, Blake A Richards, Guillaume Lajoie, and Bratislav Misic. Learning function from structure in neuromorphic networks. Nature Machine Intelligence, 3(9): 771–786, 2021.

[140] Andrea I Luppi, S Parker Singleton, Justine Y Hansen, Keith W Jamison, Danilo Bzdok, Amy Kuceyeski, Richard F Betzel, and Bratislav Misic. Contributions of network structure, chemoarchitecture and diagnostic categories to transitions between cognitive topographies. Nature Biomedical Engineering, 8(9):1142–1161, 2024.

[141] Istvan Törk. Anatomy of the serotonergic system a. Annals of the New York Academy of Sciences, 600(1):9–34, 1990.

[142] M-Marsel Mesulam and Changiz Geula. Nucleus basalis (ch4) and cortical cholinergic innervation in the human brain: observations based on the distribution of acetylcholinesterase and choline acetyltransferase. Journal of Comparative Neurology, 275(2):216–240, 1988.

[143] Satoshi Ikemoto. Dopamine reward circuitry: two projection systems from the ventral midbrain to the nucleus accumbens–olfactory tubercle complex. Brain Research Reviews, 56(1):27–78, 2007.

[144] Susan J Sara. The locus coeruleus and noradrenergic modulation of cognition. Nature Reviews Neuroscience, 10(3):211–223, 2009.

[145] Golia Shafiei, Yashar Zeighami, Crystal A Clark, Jennifer T Coull, Atsuko Nagano-Saito, Marco Leyton, Alain Dagher, and Bratislav Mišić. Dopamine signaling modulates the stability and integration of intrinsic brain networks. Cerebral Cortex, 29(1):397–409, 2019.

[146] Eric G Ceballos, Asa Farahani, Zhen-Qi Liu, Filip Milisav, Justine Y Hansen, Alain Dagher, and Bratislav Misic. Mapping neuropeptide signaling in the human brain. bioRxiv, pages 2024–12, 2024.

[147] James M Shine. Neuromodulatory control of complex adaptive dynamics in the brain. Interface Focus, 13(3):20220079, 2023.

[148] Lea Tenenholz Grinberg, Udo Rueb, and Helmut Heinsen. Brainstem: neglected locus in neurodegenerative diseases. Frontiers in Neurology, 2:42, 2011.

[149] Kelvin C Luk, Victoria Kehm, Jenna Carroll, Bin Zhang, Patrick O’Brien, John Q Trojanowski, and Virginia M-Y Lee. Pathological α-synuclein transmission initiates parkinson-like neurodegeneration in nontransgenic mice. Science, 338(6109):949–953, 2012.

[150] Ying-Qiu Zheng, Yu Zhang, Yvonne Yau, Yashar Zeighami, Kevin Larcher, Bratislav Misic, and Alain Dagher. Local vulnerability and global connectivity jointly shape neurodegenerative disease propagation. PLoS biology, 17(11):e3000495, 2019.

[151] Christina Tremblay, Shady Rahayel, Andrew Vo, Filip Morys, Golia Shafiei, Nooshin Abbasi, Ross D Markello, Ziv Gan-Or, Bratislav Misic, and Alain Dagher. Brain atrophy progression in parkinson’s disease is shaped by connectivity and local vulnerability. Brain Communications, 3(4):fcab269, 2021.

[152] Andrew Vo, Christina Tremblay, Shady Rahayel, Golia Shafiei, Justine Y Hansen, Yvonne Yau, Bratislav Misic, and Alain Dagher. Network connectivity and local transcriptomic vulnerability underpin cortical atrophy progression in parkinson’s disease. NeuroImage: Clinical, 40:103523, 2023.

[153] Andrew Vo, Christina Tremblay, Shady Rahayel, Sarah Al-Bachari, Henk W Berendse, Joanna K Bright, Fernando Cendes, Emile d’Angremont, John C Dalrymplealford, Ines Debove, et al. Global network and local vulnerabilities underlie brain atrophy across parkinson’s disease stages. Brain, page awaf432, 2025.

[154] Joel C Watts, Kurt Giles, Abby Oehler, Lefkos Middleton, David T Dexter, Steve M Gentleman, Stephen J DeArmond, and Stanley B Prusiner. Transmission of multiple system atrophy prions to transgenic mice. Proceedings of the National Academy of Sciences, 110(48): 19555–19560, 2013.

[155] Lydia Chougar, Christina Tremblay, Aline Delva, Marie Filiatrault, Andrew Vo, Justine Y Hansen, Asa Farahani, Bratislav Misic, Parsa Khalafi, Charles-Etienne Castonguay, et al. Mri-derived atrophy in multiple system atrophy aligns with mitochondrial and glial gene expression patterns. npj Parkinson’s Disease, 2025.

[156] Sveva Fornari, Amelie Schäfer, Mathias Jucker, Alain Goriely, and Ellen Kuhl. Prion-like spreading of alzheimer’s disease within the brain’s connectome. Journal of the Royal Society Interface, 16(159):20190356, 2019.

[157] Tahmineh Taheri, Asa Farahani, Zhen-Qi Liu, Eric G Ceballos, Adil Harroud, Alain Dagher, and Bratislav Misic. Spatial organization of aqp4 water channel expression in the human brain. Science Advances, 12(39):eaeg8117, 2026.

[158] Golia Shafiei, Vincent Bazinet, Mahsa Dadar, Ana L Manera, D Louis Collins, Alain Dagher, Barbara Borroni, Raquel Sanchez-Valle, Fermin Moreno, Robert Laforce Jr, et al. Network structure and transcriptomic vulnerability shape atrophy in frontotemporal dementia. Brain, 146 (1):321–336, 2023.

[159] Magdalini Polymenidou and Don W Cleveland. The seeds of neurodegeneration: prion-like spreading in als. Cell, 147(3):498–508, 2011.

[160] Asa Farahani, Justine Y Hansen, Vincent Bazinet, Golia Shafiei, D Louis Collins, Mahsa Dadar, Sanjay Kalra, Alain Dagher, and Bratislav Misic. Network spreading and local biological vulnerability in amyotrophic lateral sclerosis. Communications Biology, 8(1):1153, 2025.

[161] Masami Masuda-Suzukake, Takashi Nonaka, Masato Hosokawa, Takayuki Oikawa, Tetsuaki Arai, Haruhiko Akiyama, David MA Mann, and Masato Hasegawa. Prion-like spreading of pathological α-synuclein in brain. Brain, 136(4):1128–1138, 2013.

[162] Jason D Warren, Jonathan D Rohrer, Jonathan M Schott, Nick C Fox, John Hardy, and Martin N Rossor. Molecular nexopathies: a new paradigm of neurodegenerative disease. Trends in Neurosciences, 36(10):561–569, 2013.

[163] Jing L Guo and Virginia MY Lee. Cell-to-cell transmission of pathogenic proteins in neurodegenerative diseases. Nature Medicine, 20(2):130–138, 2014.

[164] Per Borghammer, Jacob Horsager, Katrine Andersen, Nathalie Van Den Berge, Anna Raunio, Shigeo Murayama, Laura Parkkinen, and Liisa Myllykangas. Neuropathological evidence of body-first vs. brain-first lewy body disease. Neurobiology of Disease, 161:105557, 2021.

[165] Per Borghammer, Mie Kristine Just, Jacob Horsager, Casper Skjærbæk, Anna Raunio, Eloise H Kok, Sara Savola, Shigeo Murayama, Yuko Saito, Liisa Myllykangas, et al. A postmortem study suggests a revision of the dual-hit hypothesis of parkinson’s disease. npj Parkinson’s Disease, 8(1):166, 2022.

[166] Shady Rahayel, Bratislav Mišić, Ying-Qiu Zheng, Zhen-Qi Liu, Alaa Abdelgawad, Nooshin Abbasi, Anna Caputo, Bin Zhang, Angela Lo, Victoria Kehm, et al. Differentially targeted seeding reveals unique pathological alpha-synuclein propagation patterns. Brain, 145(5): 1743–1756, 2022.

[167] Yasser Iturria-Medina, Roberto C Sotero, Paule J Toussaint, Alan C Evans, and Alzheimer’s Disease Neuroimaging Initiative. Epidemic spreading model to characterize misfolded proteins propagation in aging and associated neurodegenerative disorders. PLoS computational biology, 10(11):e1003956, 2014.

[168] Ashish Raj, Amy Kuceyeski, and Michael Weiner. A network diffusion model of disease progression in dementia. Neuron, 73(6):1204–1215, 2012.

[169] Jacob W Vogel, Nick Corriveau-Lecavalier, Nicolai Franzmeier, Joana B Pereira, Jesse A Brown, Anne Maass, Hugo Botha, William W Seeley, Dani S Bassett, David T Jones, et al. Connectome-based modelling of neurodegenerative diseases: towards precision medicine and mechanistic insight. Nature Reviews Neuroscience, 24 (10):620–639, 2023.

[170] JG Greenfield and Frances D Bosanquet. The brain-stem lesions in parkinsonism. Journal of Neurology, Neurosurgery, and Psychiatry, 16(4):213, 1953.

[171] Heiko Braak, Kelly Del Tredici, Udo Rüb, Rob AI De Vos, Ernst NH Jansen Steur, and Eva Braak. Staging of brain pathology related to sporadic parkinson’s disease. Neurobiology of Aging, 24(2):197–211, 2003.

[172] Johannes Brettschneider, Kelly Del Tredici, Jon B Toledo, John L Robinson, David J Irwin, Murray Grossman, EunRan Suh, Vivianna M Van Deerlin, Elisabeth M Wood, Young Baek, et al. Stages of ptdp-43 pathology in amyotrophic lateral sclerosis. Annals of Neurology, 74(1): 20–38, 2013.

[173] Johannes Brettschneider, Kimihito Arai, Kelly Del Tredici, Jon B Toledo, John L Robinson, Edward B Lee, Satoshi Kuwabara, Kazumoto Shibuya, David J Irwin, Lubin Fang, et al. Tdp-43 pathology and neuronal loss in amyotrophic lateral sclerosis spinal cord. Acta Neuropathologica, 128(3):423–437, 2014.

[174] Matthew D Cykowski, Hidehiro Takei, Paul E Schulz, Stanley H Appel, and Suzanne Z Powell. Tdp-43 pathology in the basal forebrain and hypothalamus of patients with amyotrophic lateral sclerosis. Acta Neuropathologica Communications, 2(1):171, 2014.

[175] Kay Seidel, Josefine Mahlke, Sonny Siswanto, Reijko Krüger, Helmut Heinsen, Georg Auburger, Mohamed Bouzrou, Lea T Grinberg, Helmut Wicht, Horst-Werner Korf, et al. The brainstem pathologies of parkinson’s disease and dementia with lewy bodies. Brain Pathology, 25(2):121–135, 2015.

[176] Josef Parvizi, Gary W Van Hoesen, and Antonio Damasio. The selective vulnerability of brainstem nuclei to alzheimer’s disease. Annals of Neurology: Official Journal of the American Neurological Association and the Child Neurology Society, 49(1):53–66, 2001.

[177] Martina Bocchetta, Maura Malpetti, Emily G Todd, James B Rowe, and Jonathan D Rohrer. Looking beneath the surface: the importance of subcortical structures in frontotemporal dementia. Brain Communications, 3(3):fcab158, 2021.

[178] Derek K Jones. Challenges and limitations of quantifying brain connectivity in vivo with diffusion mri. Imaging in Medicine, 2(3):341, 2010.

[179] Derek K Jones. Studying connections in the living human brain with diffusion mri. Cortex, 44(8):936–952, 2008.

[180] Saad Jbabdi, Stamatios N Sotiropoulos, Suzanne N Haber, David C Van Essen, and Timothy E Behrens. Measuring macroscopic brain connections in vivo. Nature Neuroscience, 18(11):1546–1555, 2015.

[181] Robert E Smith, Jacques-Donald Tournier, Fernando Calamante, and Alan Connelly. Anatomically-constrained tractography: improved diffusion mri streamlines tractography through effective use of anatomical information. Neuroimage, 62(3):1924–1938, 2012.

[182] Robert E Smith, Jacques-Donald Tournier, Fernando Calamante, and Alan Connelly. Sift: Spherical-deconvolution informed filtering of tractograms. Neuroimage, 67:298–312, 2013.

[183] Marshall A Dalton, Arkiev D’Souza, Jinglei Lv, and Fernando Calamante. New insights into anatomical connectivity along the anterior–posterior axis of the human hippocampus using in vivo quantitative fibre tracking. elife, 11:e76143, 2022.

[184] Samuel R Krimmel, Timothy O Laumann, Roselyne J Chauvin, Tamara Hershey, Jarod L Roland, Joshua S Shimony, Jon T Willie, Scott A Norris, Scott Marek, Andrew N. Van, et al. The human brainstem’s red nucleus was upgraded to support goal-directed action. Nature communications, 16(1):3398, 2025.

[185] Kathy Y Liu, Freya Marijatta, Dorothea Hämmerer, Julio Acosta-Cabronero, Emrah Düzel, and Robert J Howard. Magnetic resonance imaging of the human locus coeruleus: A systematic review. Neuroscience & Biobehavioral Reviews, 83:325–355, 2017.

[186] Boris Keil, Christina Triantafyllou, Michael Hamm, and Lawrence L Wald. Design optimization of a 32-channel head coil at 7t. In Proceedings of the International Society for Magnetic Resonance in Medicine, volume 18, page 1493, 2010.

[187] José V Manjón, Pierrick Coupé, Luis Concha, Antonio Buades, D Louis Collins, and Montserrat Robles. Diffusion weighted image denoising using overcomplete local pca. PloS one, 8(9):e73021, 2013.

[188] Brian B Avants, Nicholas J Tustison, Jue Wu, Philip A Cook, and James C Gee. An open source multivariate framework for n-tissue segmentation with evaluation on public data. Neuroinformatics, 9(4):381–400, 2011.

[189] Christophe Destrieux, Bruce Fischl, Anders Dale, and Eric Halgren. Automatic parcellation of human cortical gyri and sulci using standard anatomical nomenclature. Neuroimage, 53(1):1–15, 2010.

[190] Wolfgang M Pauli, Amanda N Nili, and J Michael Tyszka. A high-resolution probabilistic in vivo atlas of human subcortical brain nuclei. Scientific Data, 5(1):1–13, 2018.

[191] Robert W Cox. Afni: software for analysis and visualization of functional magnetic resonance neuroimages. Computers and Biomedical Research, 29(3):162–173, 1996.

[192] Rasmus M Birn, Monica A Smith, Tyler B Jones, and Peter A Bandettini. The respiration response function: the temporal dynamics of fmri signal fluctuations related to changes in respiration. Neuroimage, 40(2):644–654, 2008.

[193] Catie Chang, John P Cunningham, and Gary H Glover. Influence of heart rate on the bold signal: the cardiac response function. Neuroimage, 44(3):857–869, 2009.

[194] Ajay B Satpute, Tor D Wager, Julien Cohen-Adad, Marta Bianciardi, Ji-Kyung Choi, Jason T Buhle, Lawrence L Wald, and Lisa Feldman Barrett. Identification of discrete functional subregions of the human periaqueductal gray. Proceedings of the National Academy of Sciences, 110(42):17101–17106, 2013.

[195] Marta Bianciardi, Nicola Toschi, Cornelius Eichner, Jonathan R Polimeni, Kawin Setsompop, Emery N Brown, Matti S Hämäläinen, Bruce R Rosen, and Lawrence L Wald. In vivo functional connectome of human brainstem nuclei of the ascending arousal, autonomic, and motor systems by high spatial resolution 7-tesla fmri. Magnetic Resonance Materials in Physics, Biology and Medicine, 29:451–462, 2016.

[196] G Paxinos, X Huang, G Sengul, and C Watson. Organization of brainstem nuclei. The Human Nervous System, pages 260–327, 2012.

[197] J-Donald Tournier, Fernando Calamante, and Alan Connelly. Mrtrix: diffusion tractography in crossing fiber regions. International Journal of Imaging Systems and Technology, 22(1):53–66, 2012.

[198] J-Donald Tournier, Robert Smith, David Raffelt, Rami Tabbara, Thijs Dhollander, Maximilian Pietsch, Daan Christiaens, Ben Jeurissen, Chun-Hung Yeh, and Alan Connelly. Mrtrix3: A fast, flexible and open software framework for medical image processing and visualisation. Neuroimage, 202:116137, 2019.

[199] J-Donald Tournier, Fernando Calamante, and Alan Connelly. Robust determination of the fibre orientation distribution in diffusion mri: non-negativity constrained super-resolved spherical deconvolution. Neuroimage, 35 (4):1459–1472, 2007.

[200] J-Donald Tournier, Fernando Calamante, and Alan Connelly. Determination of the appropriate b value and number of gradient directions for high-angular-resolution diffusion-weighted imaging. NMR in Biomedicine, 26(12):1775–1786, 2013.

[201] J Donald Tournier, Fernando Calamante, Alan Connelly, et al. Improved probabilistic streamlines tractography by 2nd order integration over fibre orientation distributions. In Proceedings of the international society for magnetic resonance in medicine, volume 1670. Stockholm, 2010.

[202] James A Roberts, Alistair Perry, Gloria Roberts, Philip B Mitchell, and Michael Breakspear. Consistency-based thresholding of the human connectome. Neuroimage, 145:118–129, 2017.

[203] Marcel A de Reus and Martijn P van den Heuvel. Estimating false positives and negatives in brain networks. Neuroimage, 70:402–409, 2013.

[204] Derek K Jones, Thomas R Knösche, and Robert Turner. White matter integrity, fiber count, and other fallacies: the do’s and don’ts of diffusion mri. Neuroimage, 73: 239–254, 2013.

[205] Cibu Thomas, Frank Q Ye, M Okan Irfanoglu, Pooja Modi, Kadharbatcha S Saleem, David A Leopold, and Carlo Pierpaoli. Anatomical accuracy of brain connections derived from diffusion mri tractography is inherently limited. Proceedings of the National Academy of Sciences, 111(46):16574–16579, 2014.

[206] Richard F Betzel, Andrea Avena-Koenigsberger, Joaquín Goñi, Ye He, Marcel A De Reus, Alessandra Griffa, Petra E Vértes, Bratislav Mišic, Jean-Philippe Thiran, Patric Hagmann, et al. Generative models of the human connectome. Neuroimage, 124:1054–1064, 2016.

[207] Mark Jenkinson, Christian F Beckmann, Timothy EJ Behrens, Mark W Woolrich, and Stephen M Smith. Fsl. Neuroimage, 62(2):782–790, 2012.

[208] Brian B Avants, Nicholas J Tustison, Gang Song, Philip A Cook, Arno Klein, and James C Gee. A reproducible evaluation of ants similarity metric performance in brain image registration. Neuroimage, 54(3):2033–2044, 2011.

[209] Bruce Fischl. Freesurfer. Neuroimage, 62(2):774–781, 2012.

[210] Shengwei Zhang and Konstantinos Arfanakis. Evaluation of standardized and study-specific diffusion tensor imaging templates of the adult human brain: Template characteristics, spatial normalization accuracy, and detection of small inter-group fa differences. Neuroimage, 172:40–50, 2018.

[211] W Evan Johnson, Cheng Li, and Ariel Rabinovic. Adjusting batch effects in microarray expression data using empirical bayes methods. Biostatistics, 8(1):118–127, 2007.

[212] Zhen-Qi Liu, Vincent Bazinet, Justine Y Hansen, Filip Milisav, Andrea I Luppi, Eric G Ceballos, Asa Farahani, Laura E Suarez, Golia Shafiei, Ross D Markello, et al. netneurotools: a trainee-oriented approach to network neuroscience. bioRxiv, pages 2025–09, 2025.

[213] František Váša and Bratislav Mišić. Null models in network neuroscience. Nature Reviews Neuroscience, 23(8): 493–504, 2022.

[214] David C Van Essen, Matthew F Glasser, Donna L Dierker, John Harwell, and Timothy Coalson. Parcellations and hemispheric asymmetries of human cerebral cortex analyzed on surface-based atlases. Cerebral Cortex, 22(10): 2241–2262, 2012.

[215] Rudolf Nieuwenhuys. The myeloarchitectonic studies on the human cerebral cortex of the vogt–vogt school, and their significance for the interpretation of functional neuroimaging data. Brain Structure and Function, 218: 303–352, 2013.

[216] Stefan Van Der Walt, S Chris Colbert, and Gael Varoquaux. The numpy array: a structure for efficient numerical computation. Computing in Science & Engineering, 13(2):22–30, 2011.

[217] Charles R Harris, K Jarrod Millman, Stéfan J Van Der Walt, Ralf Gommers, Pauli Virtanen, David Cournapeau, Eric Wieser, Julian Taylor, Sebastian Berg, Nathaniel J Smith, et al. Array programming with numpy. Nature, 585(7825):357–362, 2020.

[218] Pauli Virtanen, Ralf Gommers, Travis E Oliphant, Matt Haberland, Tyler Reddy, David Cournapeau, Evgeni Burovski, Pearu Peterson, Warren Weckesser, Jonathan Bright, et al. Scipy 1.0: fundamental algorithms for scientific computing in python. Nature Methods, 17(3): 261–272, 2020.

[219] Wes McKinney et al. Data structures for statistical computing in python. SciPy, 445(1):51–56, 2010.

[220] Michael L. Waskom. seaborn: statistical data visualization. Journal of Open Source Software, 6(60):3021, 2021. doi:10.21105/joss.03021. URL https://doi.org/10.21105/joss.03021.

[221] John D Hunter. Matplotlib: A 2d graphics environment. Computing in Science & Engineering, 9(03):90–95, 2007.

[222] Skipper Seabold and Josef Perktold. statsmodels: Econometric and statistical modeling with python. In 9th Python in Science Conference, 2010.

[223] Alexandre Abraham, Fabian Pedregosa, Michael Eickenberg, Philippe Gervais, Andreas Mueller, Jean Kossaifi, Alexandre Gramfort, Bertrand Thirion, and Gaël Varoquaux. Machine learning for neuroimaging with scikitlearn. Frontiers in Neuroinformatics, 8:14, 2014.

[224] Matthew Brett, Christopher J. Markiewicz, Michael Hanke, Marc-Alexandre Côté, Ben Cipollini, Paul McCarthy, Dorota Jarecka, Christopher P. Cheng, Yaroslav O. Halchenko, Michiel Cottaar, Eric Larson, Satrajit Ghosh, Demian Wassermann, Stephan Gerhard, Gregory R. Lee, Hao-Ting Wang, Erik Kastman, Jakub Kaczmarzyk, Roberto Guidotti, Jonathan Daniel, Or Duek, Ariel Rokem, Cindee Madison, Brendan Moloney, Félix C. Morency, Mathias Goncalves, Ross Markello, Cameron Riddell, Anibal Sólon, Christopher Burns, Jarrod Millman, Alexandre Gramfort, Jaakko Leppäkangas, Jasper J.F. van den Bosch, Robert D. Vincent, Henry Braun, Krish Subramaniam, Dimitri Papadopoulos Orfanos, Andrew Van, Krzysztof J. Gorgolewski, Pradeep Reddy Raamana, Julian Klug, B. Nolan Nichols, Eric M. Baker, Soichi Hayashi, Basile Pinsard, Christian Haselgrove, Mark Hymers, Oscar Esteban, Serge Koudoro, Fernando Pérez-García, Jérôme Dockès, Nikolaas N. Oosterhof, Bago Amirbekian, Ian Nimmo-Smith, Ly Nguyen, Samir Reddigari, Samuel St-Jean, Egor Panfilov, Eleftherios Garyfallidis, Gael Varoquaux, Jon Haitz Legarreta, Kevin S. Hahn, Lea Waller, Oliver P. Hinds, Bennet Fauber, Jacob Roberts, Jean-Baptiste Poline, Jon Stutters, Kesshi Jordan, Matthew Cieslak, Miguel Estevan Moreno, Tomáš Hrnčiar, Valentin Haenel, Yannick Schwartz, Zvi Baratz, Benjamin C Darwin, Bertrand Thirion, Carl Gauthier, Igor Solovey, Ivan Gonzalez, Jath Palasubramaniam, Justin Lecher, Katrin Leinweber, Konstantinos Raktivan, Markéta Calábková, Peter Fischer, Philippe Gervais, Syam Gadde, Thomas Ballinger, Thomas Roos, Venkateswara Reddy Reddam, and freec84. nipy/nibabel:, June 2022. URL 10.5281/zenodo.6658382.

[225] Michael F Huerta and John K Harting. Connectional organization of the superior colliculus. Trends in Neurosciences, 7(8):286–289, 1984.

[226] John K Harting. Descending pathways from the superior colliculus: an autoradiographic analysis in the rhesus monkey (macaca mulatta). Journal of Comparative Neurology, 173(3):583–612, 1977.

[227] Jeffrey G Mellott, Nichole L Beebe, and Brett R Schofield. Gabaergic and non-gabaergic projections to the superior colliculus from the auditory brainstem. Brain Structure and Function, 223(4):1923–1936, 2018.

[228] Alfonso Deichler, Denisse Carrasco, Luciana Lopez-Jury, Tomas Vega-Zuniga, Natalia Márquez, Jorge Mpodozis, and Gonzalo J Marín. A specialized reciprocal connectivity suggests a link between the mechanisms by which the superior colliculus and parabigeminal nucleus produce defensive behaviors in rodents. Scientific Reports, 10(1):16220, 2020.

[229] Alvin J Beitz. Possible origin of glutamatergic projections to the midbrain periaqueductal gray and deep layer of the superior colliculus of the rat. Brain Research Bulletin, 23(1-2):25–35, 1989.

[230] B Cohen and JA Büttner-Ennever. Projections from the superior colliculus to a region of the central mesencephalic reticular formation (cmrf) associated with horizontal saccadic eye movements. Experimental Brain Research, 57(1):167–176, 1984.

[231] Paul J May. The mammalian superior colliculus: laminar structure and connections. Progress in Brain Research, 151:321–378, 2006.

[232] Niping Wang, Susan Warren, and Paul J May. The macaque midbrain reticular formation sends side-specific feedback to the superior colliculus. Experimental Brain Research, 201(4):701–717, 2010.

[233] Niping Wang, Eddie Perkins, Lan Zhou, Susan Warren, and Paul J May. Reticular formation connections underlying horizontal gaze: the central mesencephalic reticular formation (cmrf) as a conduit for the collicular saccade signal. Frontiers in Neuroanatomy, 11:36, 2017.

[234] Michael F Huerta, Leah A Krubitzer, and Jon H Kaas. Frontal eye field as defined by intracortical microstimulation in squirrel monkeys, owl monkeys, and macaque monkeys: I. subcortical connections. Journal of Comparative Neurology, 253(4):415–439, 1986.

[235] GB Stanton, ME Goldberg, and CJ Bruce. Frontal eye field efferents in the macaque monkey: Ii. topography of terminal fields in midbrain and pons. Journal of Comparative Neurology, 271(4):493–506, 1988.

[236] Michael F Huerta and Jon H Kaas. Supplementary eye field as defined by intracortical microstimulation: connections in macaques. Journal of Comparative Neurology, 293(2):299–330, 1990.

[237] John K Harting and Bruce V Updyke. Oculomotor-related pathways of the basal ganglia. Progress in Brain Research, 151:441–460, 2006.

[238] Xue Liu, Hongren Huang, Terrance P Snutch, Peng Cao, Liping Wang, and Feng Wang. The superior colliculus: cell types, connectivity, and behavior. Neuroscience Bulletin, 38(12):1519–1540, 2022.

[239] A Jayaraman, Robert R Batton III, and Malcolm B Carpenter. Nigrotectal projections in the monkey: an autoradiographic study. Brain Research, 135(1):147–152, 1977.

[240] Katsuyuki Kaneda, Kaoru Isa, Yuchio Yanagawa, and Tadashi Isa. Nigral inhibition of gabaergic neurons in mouse superior colliculus. Journal of Neuroscience, 28 (43):11071–11078, 2008.

[241] Eliane Comoli, Véronique Coizet, Justin Boyes, J Paul Bolam, Newton S Canteras, Rachel H Quirk, Paul G Overton, and Peter Redgrave. A direct projection from superior colliculus to substantia nigra for detecting salient visual events. Nature Neuroscience, 6(9):974–980, 2003.

[242] GM Krauthamer, BS Grunwerg, and H Krein. Putative cholinergic neurons of the pedunculopontine tegmental nucleus projecting to the superior colliculus consist of sensory responsive and unresponsive populations which are functionally distinct from other mesopontine neurons. Neuroscience, 69(2):507–517, 1995.

[243] MN Wallace and K Fredens. Origin of high acetylcholinesterase activity in the mouse superior colliculus. Experimental Brain Research, 72(2):335–346, 1988.

[244] William C Hall, David Fitzpatrick, LL Klatt, and Denis Raczkowski. Cholinergic innervation of the superior colliculus in the cat. Journal of Comparative Neurology, 287 (4):495–514, 1989.

[245] P Redgrave, P Dean, IJ Mitchell, A Odekunle, and A Clark. The projection from superior colliculus to cuneiform area in the rat: I. anatomical studies. Experimental Brain Research, 72(3):611–625, 1988.

[246] IJ Mitchell, P Dean, and P Redgrave. The projection from superior colliculus to cuneiform area in the rat: Ii. defence-like responses to stimulation with glutamate in cuneiform nucleus and surrounding structures. Experimental Brain Research, 72(3):626–639, 1988.

[247] Nora L Benavidez, Michael S Bienkowski, Muye Zhu, Luis H Garcia, Marina Fayzullina, Lei Gao, Ian Bowman, Lin Gou, Neda Khanjani, Kaelan R Cotter, et al. Organization of the inputs and outputs of the mouse superior colliculus. Nature Communications, 12(1):4004, 2021.

[248] Aloisio Tortelly and Fernando Reinoso-Suárez. Projections to the superior colliculus from the dorsal hypothalamic area and other prosencephalic structures derived from the embryonic subthalamic longitudinal band of the diencephalon. Neuroscience Letters, 18(3):257–260, 1980.

[249] Ada Eban-Rothschild, Gideon Rothschild, William J Giardino, Jeff R Jones, and Luis De Lecea. Vta dopaminergic neurons regulate ethologically relevant sleep–wake behaviors. Nature neuroscience, 19(10):1356–1366, 2016.

[250] Brian L Edlow, Emi Takahashi, Ona Wu, Thomas Benner, Guangping Dai, Lihong Bu, Patricia Ellen Grant, David M Greer, Steven M Greenberg, Hannah C Kinney, et al. Neuroanatomic connectivity of the human ascending arousal system critical to consciousness and its disorders. Journal of Neuropathology & Experimental Neurology, 71(6):531–546, 2012.

[251] Wei Tang, Olexiy Kochubey, Michael Kintscher, and Ralf Schneggenburger. A vta to basal amygdala dopamine projection contributes to signal salient somatosensory events during fear learning. Journal of Neuroscience, 40 (20):3969–3980, 2020.

[252] Roy A Wise. Dopamine, learning and motivation. Nature Reviews Neuroscience, 5(6):483–494, 2004.

[253] Qi Wu, Maureen P Boyle, and Richard D Palmiter. Loss of gabaergic signaling by agrp neurons to the parabrachial nucleus leads to starvation. Cell, 137(7):1225–1234, 2009.

[254] William Gordon Frankle, Mark Laruelle, and Suzanne N Haber. Prefrontal cortical projections to the midbrain in primates: evidence for a sparse connection. Neuropsychopharmacology, 31(8):1627–1636, 2006.

[255] Susan R Sesack and Virginia M Pickel. Prefrontal cortical efferents in the rat synapse on unlabeled neuronal targets of catecholamine terminals in the nucleus accumbens septi and on dopamine neurons in the ventral tegmental area. Journal of Comparative Neurology, 320 (2):145–160, 1992.

[256] David B Carr and Susan R Sesack. Gaba-containing neurons in the rat ventral tegmental area project to the prefrontal cortex. Synapse, 38(2):114–123, 2000.

[257] Kim Wager and Sue Cox. Chapter 6—the limbic (emotional) system. Auricular acupuncture & addiction, pages 57–67, 2009.

[258] Ali Charara, Yoland Smith, and André Parent. Glutamatergic inputs from the pedunculopontine nucleus to midbrain dopaminergic neurons in primates: Phaseolus vulgaris-leucoagglutinin anterograde labeling combined with postembedding glutamate and gaba immunohistochemistry. Journal of Comparative Neurology, 364(2): 254–266, 1996.

[259] Ericka C Holmstrand and Susan R Sesack. Projections from the rat pedunculopontine and laterodorsal tegmental nuclei to the anterior thalamus and ventral tegmental area arise from largely separate populations of neurons. Brain Structure and Function, 216(4):331–345, 2011.

[260] Juan Mena-Segovia. Structural and functional considerations of the cholinergic brainstem. Journal of Neural Transmission, 123(7):731–736, 2016.

[261] Hitoshi Morikawa and Carlos A Paladini. Dynamic regulation of midbrain dopamine neuron activity: intrinsic, synaptic, and plasticity mechanisms. Neuroscience, 198: 95–111, 2011.

[262] Jia Qi, Shiliang Zhang, Hui-Ling Wang, Huikun Wang, Jose de Jesus Aceves Buendia, Alexander F Hoffman, Carl R Lupica, Rebecca P Seal, and Marisela Morales. A glutamatergic reward input from the dorsal raphe to ventral tegmental area dopamine neurons. Nature Communications, 5(1):5390, 2014.

[263] Mitsuko Watabe-Uchida, Lisa Zhu, Sachie K Ogawa, Archana Vamanrao, and Naoshige Uchida. Whole-brain mapping of direct inputs to midbrain dopamine neurons. Neuron, 74(5):858–873, 2012.

[264] Sile An, Xiangning Li, Lei Deng, Peilin Zhao, Zhangheng Ding, Yutong Han, Yue Luo, Xin Liu, Anan Li, Qingming Luo, et al. A whole-brain connectivity map of vta and snc glutamatergic and gabaergic neurons in mice. Frontiers in Neuroanatomy, 15:818242, 2021.

[265] Nao Chuhma, Susana Mingote, Holly Moore, and Stephen Rayport. Dopamine neurons control striatal cholinergic neurons via regionally heterogeneous dopamine and glutamate signaling. Neuron, 81(4):901–912, 2014.

[266] Stefanie Geisler and Daniel S Zahm. Afferents of the ventral tegmental area in the rat-anatomical substratum for integrative functions. Journal of Comparative Neurology, 490(3):270–294, 2005.

[267] Robert M Beckstead, Valerie B Domesick, and Walle JH Nauta. Efferent connections of the substantia nigra and ventral tegmental area in the rat. Brain Research, 175 (2):191–217, 1979.

[268] Larry W. Swanson. The projections of the ventral tegmental area and adjacent regions: a combined fluorescent retrograde tracer and immunofluorescence study in the rat. Brain Research Bulletin, 9(1-6):321–353, 1982.

[269] Yu Qiao, Chun-Kui Zhang, Zhi-Hong Li, Ze-Hao Niu, Jin Li, and Jin-Lian Li. Collateral projections from the lateral parabrachial nucleus to the central amygdaloid nucleus and the ventral tegmental area in the rat. The Anatomical Record, 302(7):1178–1186, 2019.

[270] Maria Inês Nogueira, Briseida Dogo de Rezende, Luiz Eduardo Ribeiro do Vale, and Jackson Cioni Bittencourt. Afferent connections of the caudal raphe pallidus nucleus in rats: a study using the fluorescent retrograde tracers fluorogold and true-blue. Annals of Anatomyanatomischer Anzeiger, 182(1):35–45, 2000.

[271] Masaki Tanaka, Hitoshi Okamura, Yoshitaka Tamada, Ikuko Nagatsu, Yoshifumi Tanaka, and Yasuhiko Ibata. Catecholaminergic input to spinally projecting serotonin neurons in the rostral ventromedial medulla oblongata of the rat. Brain Research Bulletin, 35(1):23–30, 1994.

[272] Dirk M Hermann, Pierre-Hervé Luppi Christelle Peyron, Peter Hinckel, and Michel Jouvet. Afferent projections to the rat nuclei raphe magnus, raphe pallidus and reticularis gigantocellularis pars α demonstrated by iontophoretic application of choleratoxin (subunit b). Journal of Chemical Neuroanatomy, 13(1):1–21, 1997.

[273] AL Halberstadt and CD Balaban. Organization of projections from the raphe nuclei to the vestibular nuclei in rats. Neuroscience, 120(2):573–594, 2003.

[274] Deanna M Barch, Gregory C Burgess, Michael P Harms, Steven E Petersen, Bradley L Schlaggar, Maurizio Corbetta, Matthew F Glasser, Sandra Curtiss, Sachin Dixit, Cindy Feldt, et al. Function in the human connectome: task-fmri and individual differences in behavior. Neuroimage, 80:169–189, 2013.

[275] Neal H Barmack. Inferior olive and oculomotor system. Progress in Brain Research, 151:269–291, 2006.

[276] Tao Liu, Duo Xu, James Ashe, and Khalafalla Bushara. Specificity of inferior olive response to stimulus timing. Journal of Neurophysiology, 100(3):1557–1561, 2008.

